# Primum: a decision benchmark for selecting cryo-EM re-refinements

**DOI:** 10.64898/2026.10.31.763284

**Authors:** Xingyu Yi, Xinya Zhang, Lizhe Zhang, Jinhua Yi

**Affiliations:** INTOSYN AI Labs; Teachers College, Columbia University; Department of Pharmacology and Toxicology, University of Toronto

## Abstract

Map fit is widely used in cryo-EM re-refinement. We benchmarked selection against retaining the deposited model (no operation), using packed-side-chain agreement with higher-resolution references. In a prespecified 3.0–3.6 Å cohort (167 entries, 131 sequence groups), highest-correlation selection increased *χ*_1_ disagreement by 2.4° (95% interval 2.1–2.7°), with positive loss in 91% of groups. Group-weighted fractions with >20° increases and decreases were 7.2% and 3.7%; the fraction exceeding 1 Å local side-chain RMSD rose by 2.4 percentage points. Selection by a half map withheld from re-refinement increased disagreement (2.29°; 1.70–2.97°; 99 entries). In 53 prespecified within-study low/high-resolution pairs, loss was 3.43° (2.85–4.02°), positive in 50 pairs; half-map loss was 2.35° (1.60–3.21°; 44 pairs). Primum provides fixed cohorts, residue sets and scoring tools to benchmark refiners and selection rules against no operation.

Cryo-electron microscopy (cryo-EM) atomic models are built and refined against experimental density under stereochemical restraints^1^. At the resolutions typical of many deposited maps, local resolvability varies and sidechain conformations are often weakly determined by the data^2–5^. Deposited models are nevertheless increasingly re-refined: dedicated services re-refine archived entries⁶, new refinement methods are evaluated by re-refining deposited models^7–12^, and automated building and prediction now supply candidate coordinates at scale^13–14^. Each re-refinement implies a decision to replace or keep the deposited coordinates, usually assessed through validation scores, including agreement with the map.

The field has long recognized that agreement with the data used for refinement cannot stand alone. Independent half-map and cross-validation tests were introduced to detect over-fitting^15–19^, and current practice combines map fit with geometry, local resolvability, difference maps and other complementary scores^20–27^. Method developers have noted that deposited and refined models can be over-fitted to cryo-EM density^28–30^, that stronger map restraints degrade model geometry^31^, that refinement changes models more as resolution worsens^32^, and that treating correlated map voxels as independent rewards fitting noise^7^. Community challenges and benchmarking suites compare models with reference structures^33–35^, and benchmarking studies in other fields have shown that rankings depend on the metrics and the task definition^36–37^. These advances assess model quality and validation scores. We ask how acting on a score changes a model relative to keeping the deposited model. Crystallography anchors that decision in Rfree, computed from data withheld from refinement^38–39^; cryo-EM re-refinement workflows report geometry and map-fit scores^6–40^.

To measure this consequence directly, we treat re-refinement as a decision in which the deposited model remains a scored no-operation action alongside the refined candidates. A stated rule, such as taking the candidate with the highest same-map correlation, selects one action in each case, and the selected model is scored against a separately determined, higher-resolution structure of the same protein^30^. Cases that share a comparison structure or sequence group are averaged before independent units are weighted equally, and selection loss is the change in disagreement produced by the rule relative to no operation.

We quantify side-chain effects in the prespecified 3.0–3.6 Å cohort (167 entries), alongside 26 MEDIC models, 11 refinement configurations, the earlier 156-entry replication and 53 within-study pairs. Primum combines the cohorts, fixed scoring protocol and reference strategies into a reusable benchmark for refiners and selection rules.

## Results

### Selection loss at 3.0–3.6 Å

The prespecified cohort contained single-particle cryo-EM entries at 3.0 ≤ *d* < 3.6 Å, paired with later-released, separately determined structures at least 0.5 Å better in resolution. The 200-entry roster yielded 168 eligible and 167 analysed entries in 131 sequence groups (Methods). Every case offers no operation alongside Phenix ×1, Phenix ×2, Servalcat and CryoNet.Refine (Fig. 1a). The endpoint is chemical-symmetry-aware *χ*_1_ disagreement with the comparison structure over packed side chains (at least 18 C*α* neighbours within 10 Å), scored on common residue support across actions. Sequence groups receive equal weight.

**Figure 1.**
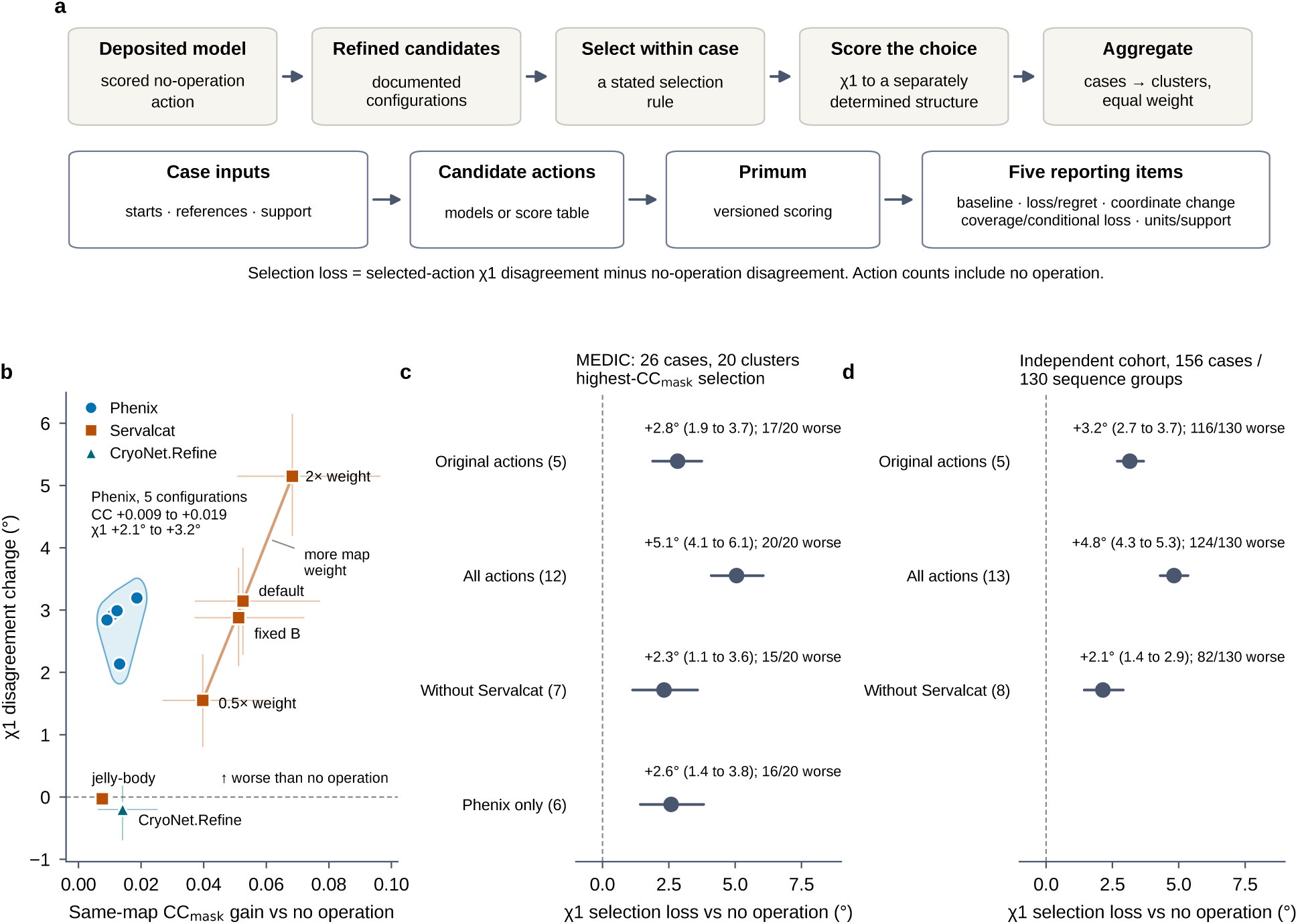
A no-operation decision benchmark and the configuration sweep. **a**, The deposited model is kept as a scored no-operation action next to refined candidates; a rule selects within each case; the choice is scored by *χ*_1_ agreement with a separately determined structure; cases are averaged within clusters that receive equal weight. The Primum workflow links case inputs and candidate actions to the five reporting items described in the Discussion. **b**, Same-map CC_mask_ gain against *χ*_1_ change relative to no operation for all 11 refinement configurations (MEDIC, 26 cases, 20 clusters); bars are 95% cluster-bootstrap intervals (Phenix intervals in Extended Data Fig. 3); the line joins Servalcat at 0.5×, 1× and 2× automatic map weight. **c**, Selection loss of the highest-CC_mask_ rule for the original five actions, all 12 actions, all actions without Servalcat and Phenix only (action counts include no operation), with clusters worse than no operation. **d**, The same rules in the prespecified independent replication (156 entries, 130 sequence groups), with sequence groups worse than no operation; no Phenix-only set was prespecified for this cohort. Positive values are worse than no operation.

Relative to no operation, Phenix ×1, Phenix ×2 and Servalcat increased *χ*_1_ disagreement by 1.38°, 1.68° and 2.63°, whereas CryoNet.Refine produced no detected mean change (−0.09°; Table 1). Their mean same-map CC_mask_ gains were +0.0098, +0.0116, +0.0265 and +0.0085, respectively (Fig. 2c).

**Figure 2.**
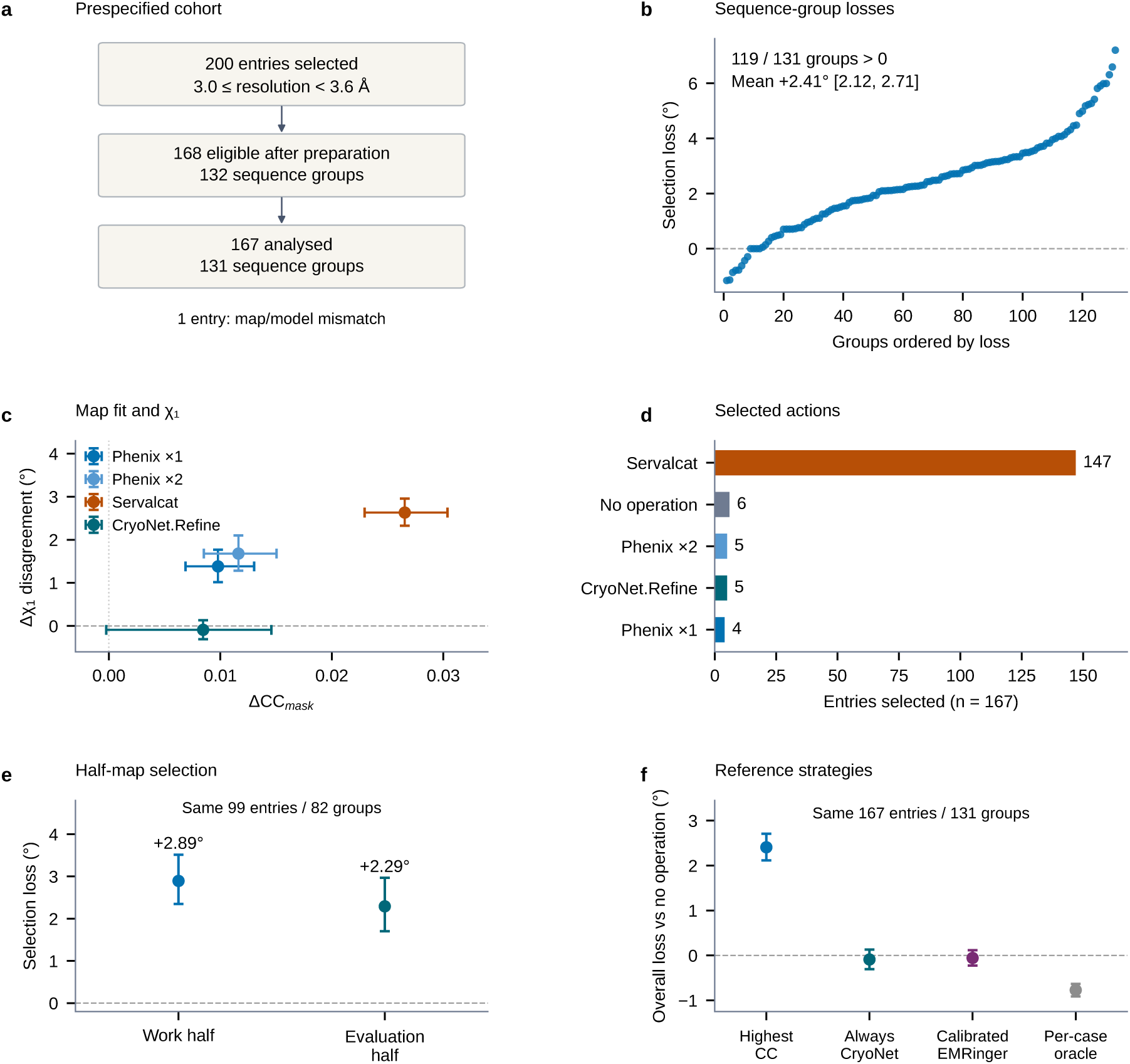
Selection by map fit at 3.0–3.6 Å. **a**, Cohort selection and analysis membership. **b**, Selection loss in all 131 sequence groups, ordered by loss, for the highest same-map CC_mask_ rule over five actions. **c**, Changes in map fit and packed-side-chain disagreement for the four refiners, on the same 167 entries. **d**, Counts of selected actions. **e**, Selection by work-half or evaluation-half CC_mask_ on the same 99 entries in 82 groups; candidates comprise no operation and three Phenix/Servalcat configurations; each configuration’s scores and endpoints are averaged over both directions before selection. **f**, Overall loss for the five-action map-fit rule, always CryoNet.Refine, the calibrated EMRinger acceptance rule and the per-case oracle on the full 167-entry sample; retained no-operation choices contribute zero loss. Points and bars are equal-group means and 95% percentile bootstrap intervals (10,000 draws, seed 20261101); positive loss indicates greater disagreement with the independent reference.

**Table 1.** Refinement effects at 3.0–3.6 Å, 167 entries and 131 sequence groups. Changes are relative to no operation, whose group-weighted mean *χ*_1_ disagreement is 27.16°. Values are group-weighted means with unadjusted 95% percentile bootstrap intervals (10,000 draws, seed 20261101).

| Action | Δχ<sub>1</sub> [95% interval] (°) | ΔCC<sub>mask</sub> [95% interval] |
|---|---|---|
| No operation | 0.00 [0.00, 0.00] | 0.0000 [0.0000, 0.0000] |
| Phenix ×1 | +1.38 [+1.02, +1.77] | +0.0098 [+0.0069, +0.0130] |
| Phenix ×2 | +1.68 [+1.28, +2.10] | +0.0116 [+0.0085, +0.0150] |
| Servalcat | +2.63 [+2.33, +2.96] | +0.0265 [+0.0229, +0.0304] |
| CryoNet.Refine | −0.09 [−0.31, +0.13] | +0.0085 [−0.0002, +0.0146] |

The rule selects the highest same-map CC_mask_ on common side-chain atoms at occupancy 1 and B = 30 Å^2^; ties favour no operation (Methods). The rule chose Servalcat in 147 of 167 cases and increased *χ*_1_ disagreement by 2.41° (95% group-bootstrap interval 2.12–2.71°); 119 of 131 groups (91%) had positive loss (Fig. 2b,d). Its observed regret, the selected action’s disagreement minus the lowest among the available actions, was 3.18°. Comparison-structure clustering gave +2.41° (2.12–2.70°; 135 clusters). Accepting each refiner only when its own CC_mask_ increased gave overall losses of 1.28°, 1.54°, 2.42° and 0.04° for Phenix ×1, Phenix ×2, Servalcat and CryoNet.Refine, respectively (Supplementary Table S39).

Among 37,504 scored side chains, disagreement increased by more than 20° in 2,806 and decreased by that amount in 1,339 (group-weighted fractions 7.2% and 3.7%). The fraction exceeding 1 Å local side-chain RMSD increased by 2.4 percentage points (95% interval 1.5–3.3), and *χ*_2_ disagreement increased by 1.97° (1.60–2.34°). There were 316 *χ*_1_-class changes among 33,153 non-valine residues (group-weighted fraction 0.87%; Supplementary Table S37).

### Lowand high-resolution pairs from the same study

An initial 22-pair cohort included five nearly superposable starting/reference pairs, motivating a further cohort with prespecified coordinate and model-building criteria (Supplementary Tables S42 and S43). Of 90 roster pairs, 53 from 53 studies met these criteria before refinement; 52 used separate acquisitions and one shared a dataset. Starting-model *χ*_1_ disagreement was at least 13.1°, and neither entry named its partner as a starting model. Low-map resolutions ranged from 3.0 to 5.15 Å (median 3.7 Å).

Highest-CC_mask_ selection increased disagreement by 3.43◦ (2.85–4.02°), with positive loss in 50/53 pairs (94%; Extended Data Fig. 10 and Supplementary Table S41). All 52 selected refinements improved CC_mask_ over no operation. Evaluation-half selection lost 2.35° (1.60–3.21°; 44 pairs). Restricting the primary analysis to the 52 separate-acquisition pairs gave 3.50° (2.93–4.10°; Supplementary Table S42).

CryoNet.Refine had the smallest mean loss among the four refiners, 0.57° (0.23–0.93°), a positive loss in contrast to the 3.0–3.6 Å cohort (−0.09°). The per-case oracle improved agreement by 0.44°, leaving observed regret of 3.87° for highest-CC_mask_ selection.

### Selection losses persist across programs and refinement settings

The original MEDIC cohort contained 26 cases and 20 reference clusters^30^; its five-action map-fit rule increased disagreement by 2.83° (1.85–3.74°; Extended Data Table 1). Protocol effects and residue-level diagnostics are reported in Supplementary Table S33, Supplementary Fig. S7 and Supplementary Note 9.

Because Servalcat dominated the selections, the next question was whether the loss persisted with other configurations and without that program. The original Servalcat protocol used the full map, whereas its developers recommend half-map input. We prespecified a sweep of documented refinement configurations (Methods): for Servalcat, jelly-body restraints, half and double its automatic map weight, fixed B factors and half-map input; for Phenix, ADP refinement, reference-model restraints, tighter geometry targets and simulated annealing. One simulated-annealing run exceeded the 4-h limit, so that configuration was excluded as prespecified; all 192 runs of the remaining eight configurations passed the technical checks.

With all 12 actions available (no operation and 11 refinement configurations), the highest-CC_mask_ rule chose Servalcat at double map weight in 24 of 26 cases and increased disagreement by 5.05° (4.10–6.06°), with all 20 clusters worse than no operation (Fig. 1c). Across the selected models, 1,132 packed side chains increased in disagreement by more than 20° and 450 decreased by that threshold (17.1% versus 6.7%, cluster-weighted). Removing every Servalcat refinement configuration left five Phenix refinement configurations and CryoNet.Refine; the rule then distributed its choices among them and still lost 2.32° (1.14–3.58°; 15/20 clusters worse); these Phenix-dominated choices changed *χ*_1_-class agreement more often and lost the comparison structure’s class more often than they gained it (206 versus 153 side chains). Phenix alone lost 2.59° (1.43–3.81°). The full-set and without-Servalcat intervals both lie above zero, meeting the prespecified criterion for a selection loss that persists after excluding Servalcat. Removing Servalcat from the original five actions alone had given a weaker 2.13° (0.68–3.85°; 11/20).

Within Servalcat, disagreement tracked map weight (Fig. 1b and Extended Data Fig. 3): halving the automatic weight increased disagreement by 1.55° (0.81–2.29°), the default by 3.15° (2.28–4.00°) and doubling it by 5.15° (4.19–6.15°), as CC_mask_ gains rose from 0.040 to 0.053 to 0.068. Jelly-body restraints, which damp coordinate shifts, gave the smallest CC_mask_ gain (0.008) and no detected mean *χ*_1_ change (−0.03°; −0.15 to 0.09°). Across programs, however, larger CC_mask_ gains did not consistently correspond to larger increases in disagreement: the five Phenix refinement configurations raised CC_mask_ by only 0.009–0.019 yet increased disagreement by 2.1–3.2°. No tested MEDIC configuration combined a CC_mask_ gain with a *χ*_1_ interval entirely at or below zero, the prespecified counterexample. Servalcat’s recommended half-map input reduced the loss by 0.53° (0.13–1.09°) in the ten cases with half maps but still increased disagreement by 3.19° (2.33–3.97°).

In a developer-generated public AQuaRef cohort (40 cases, 23 connected reference components), analysed separately because its baseline is a regularized start rather than the deposited model, AQuaRef reduced samemap CC_mask_ (−0.009; −0.013 to −0.004; Holm-adjusted P = 0.0013) while reducing *χ*_1_ disagreement (−0.93°; −1.36 to −0.51°; P = 0.0020), whereas public Servalcat increased both measures (CC_mask_ +0.038; *χ*_1_ +3.17°, 2.01–4.22°; Extended Data Fig. 5). Across the tested protocols, a gain in map fit was therefore neither necessary nor sufficient for improved mean agreement.

### Validation signals and reference support

EMRinger rewards side chains whose density peaks lie at rotameric positions²¹ and might be expected to protect side chains better than map correlation. On the 24 MEDIC cases (19 clusters) at ≤4.5 Å where it is defined, the highest-EMRinger rule refined 17 cases and lost 1.52° (0.56–2.64°), against 2.75° (1.78–3.73°) for the highestCC_mask_ rule on the same cases (Fig. 3b,c). The two signals had a cluster-weighted action-agreement fraction of 0.18 (0.03–0.37), and both increased disagreement. Geometry-based rules avoided loss chiefly by abstaining: the strictest rule returned no operation in all 24 cases, and a Pareto rule accepted three cases at +0.15° (0.00–0.40°; Fig. 3e).

**Figure 3.**
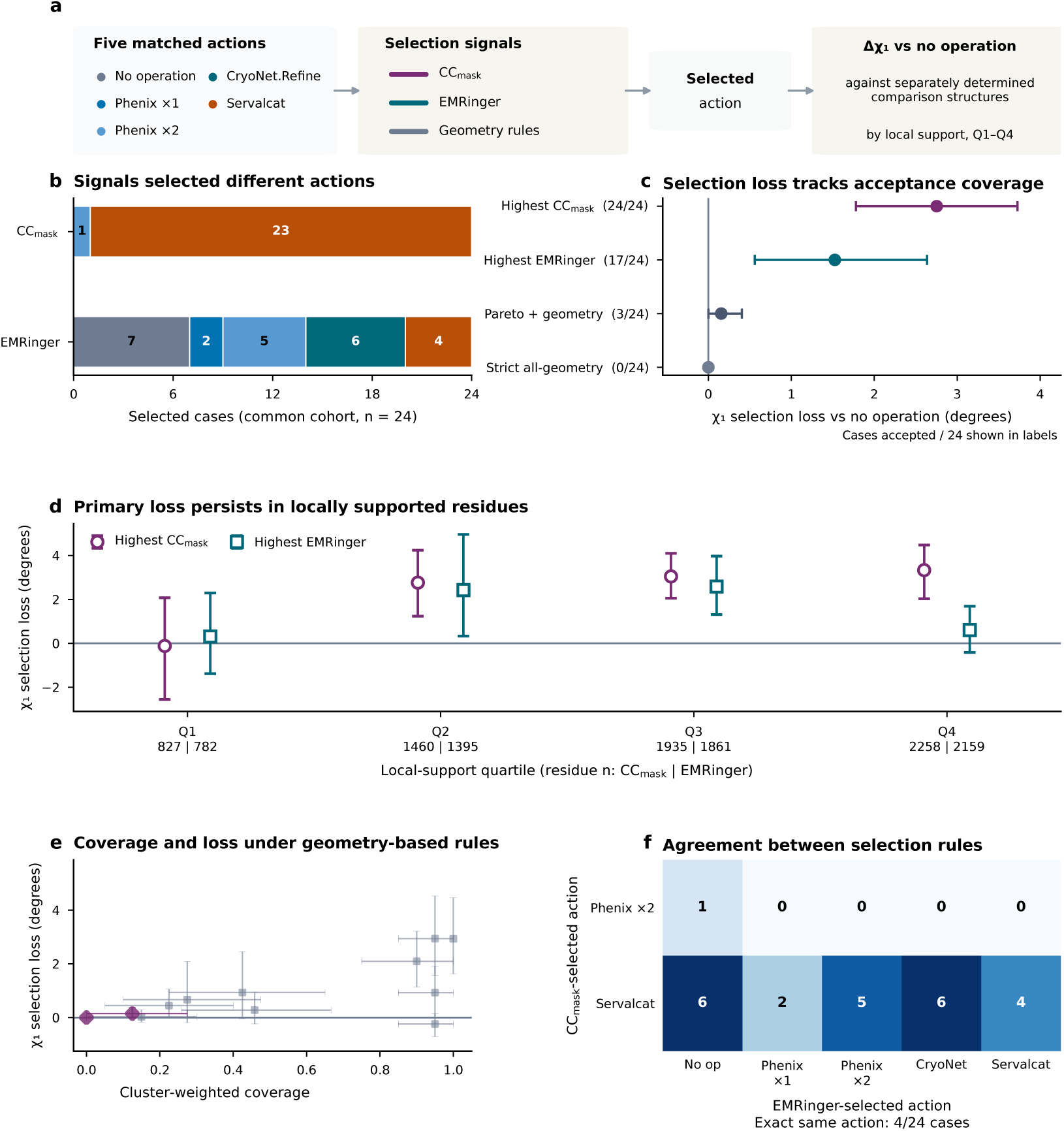
Validation signals, selection loss and reference support. **a**, Actions selected separately by CC_mask_, EMRinger or geometry. **b**, Selected actions for the highest-CC_mask_ and highest-EMRinger rules on the common 24-case, 19-cluster cohort. **c**, Selection loss of four rules (accepted cases out of 24). **d**, Selection loss across within-structure local-support quartiles of the comparison structures (residue counts for CC_mask_ and EMRinger). **e**, Coverage and loss for 12 geometry and combined selectors. **f**, Agreement between CC_mask_and EMRinger-selected actions. Points and bars are cluster-weighted means and 95% intervals.

A second question was whether the loss was concentrated in poorly supported comparison residues. Each MEDIC comparison structure’s per-residue support was computed from its own experimental data, independently of the refinement outcomes, and converted to within-structure quartiles, separately for 16 cryo-EM and four Xray comparisons (Methods). The highest-CC_mask_ rule’s loss was −0.12° (−2.55 to 2.08°) in the weakest quartile but positive in the other three, and largest in the best-supported quartile (3.33°, 2.03–4.48°; Fig. 3d). It remained positive in that quartile when only cryo-EM comparisons were used (2.83°, 1.42–4.18°). For the EMRinger rule the best-supported quartile crossed zero (0.61°, −0.41 to 1.69°).

### A prespecified independent replication

To test generality beyond MEDIC, we selected new deposited models from archive metadata alone and prespecified the predictions (Methods). Low-resolution single-particle entries (3.6–6.5 Å, released 2014–2023, protein only, at most 4,000 residues) were paired with the best-resolved structure of the same protein (95% sequenceidentity group) at ≤3.2 Å and at least 1 Å better, with a public release date later than that of the low-resolution entry. Sequence groups represented in MEDIC were excluded. A fixed identifier-derived order (Methods) accepted 200 entries (at most two per group) from 1,237 eligible pairs, and 156 entries in 130 sequence groups met the MEDIC eligibility rules (≥95% identity and ≥20 packed *χ*_1_ residues). Each was refined with the same Phenix, Servalcat and CryoNet.Refine protocols and the eight sweep configurations, and scored exactly as in MEDIC with the 95% sequence group as the independent unit.

We prespecified four predictions: positive loss for the highest-CC_mask_ rule over the five original actions (P1), over all actions (P2) and over all actions without Servalcat (P3), and 95% intervals above zero for Phenix ×1, Phenix ×2 and Servalcat, with CryoNet.Refine’s interval including zero (P4). No eligible entry or configuration was excluded from the full-map replication analysis. The highest-CC_mask_ rule over the five original actions increased disagreement by 3.16° (2.68–3.68°; 116 of 130 sequence groups worse), reproducing P1; as in MEDIC, it chose Servalcat in most entries (135 of 156). Offered all 13 actions, it chose Servalcat at double map weight in 134 entries and lost 4.82° (4.30–5.35°; 124/130), and with every Servalcat refinement configuration removed it still lost 2.14° (1.44–2.91°; 82/130), reproducing P2 and P3 (Fig. 1d and Fig. 4b). Phenix ×1, Phenix ×2 and Servalcat increased disagreement as predicted (2.84°, 3.56° and 3.02°), but CryoNet.Refine reduced it (−0.39°,

**Figure 4.**
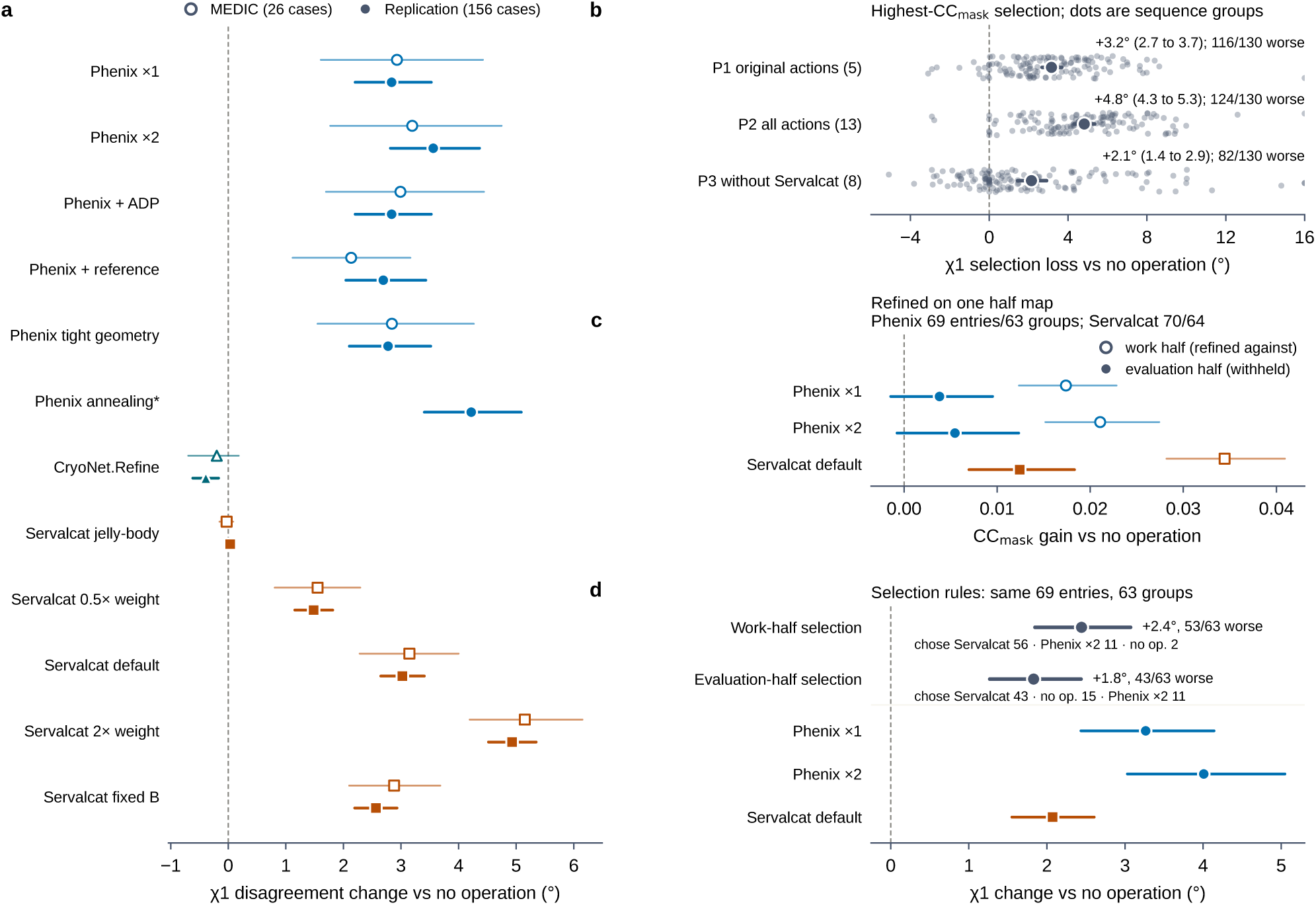
Prespecified independent replication. **a**, *χ*_1_ change relative to no operation for every refinement configuration in the replication cohort (filled symbols; 156 entries, 130 sequence groups) beside MEDIC (open symbols; 26 cases, 20 clusters); the asterisk marks the configuration excluded from MEDIC after one run exceeded the time limit. **b**, Selection loss of the highest-CC_mask_ rule for the prespecified action sets P1–P3 (action counts include no operation); dots are sequencegroup values, and values beyond the axis are drawn at its edge. **c**, In the replication entries with two deposited half maps, CC_mask_ gain of each action refined against one half map, scored on that half (open) and on the opposite evaluation half (filled); both directions are averaged within entry. Phenix ×1 and ×2, 69 entries and 63 sequence groups; Servalcat, 70 entries and 64 groups. **d**, Selection loss of choosing by evaluation-half CC_mask_ (P6) and by work-half CC_mask_ on the same 69 entries (63 groups), with one configuration selected per entry using direction-averaged scores, and the *χ*_1_ changes of the half-map-refined actions (n as in c). Cluster-weighted means with 95% cluster-bootstrap intervals; positive values are worse than no operation.

−0.62 to −0.17°) with its interval below zero, so P4 was not met (Fig. 4a). CryoNet.Refine also raised mean CC_mask_ (+0.021, 0.015–0.026). Although it improved agreement on average, the rule selected it in only 6 of 156 entries and selected Servalcat in 135, consistent with Servalcat’s larger mean gain in fit (+0.047). The rule’s observed regret was 4.09°. As in the MEDIC sweep, disagreement within Servalcat rose from 1.48° at half to 3.02° at the automatic and 4.93° at double map weight, jelly-body restraints produced no detected mean change (+0.03°, −0.02 to 0.09°), and simulated annealing, which completed in this cohort, gave the largest Phenix loss (4.22°; Fig. 4a, Extended Data Fig. 9 and Supplementary Table S32). The direction and magnitude were similar with comparison structures as the clustering unit (P1, 3.13°, 2.65–3.65°; Extended Data Fig. 9c).

### Half maps withheld from re-refinement

In the 3.0–3.6 Å cohort, 103 eligible entries listed two half maps; 99 entries in 82 groups had all candidates needed for both selection rules. Selecting by evaluation-half CC_mask_ increased disagreement by 2.29° (1.70– 2.97°), compared with 2.89° (2.35–3.51°) for work-half selection on the same entries (Fig. 2e; per-action effects in Supplementary Table S36).

Ten MEDIC cases (nine clusters) have deposited half maps. Refining against one half and evaluating on the other, in both directions, the tested Servalcat protocol increased evaluation-half CC_mask_ by 0.046 (0.017–0.082; Holm-adjusted P = 0.023) together with *χ*_1_ disagreement by 2.77° (1.90–3.60°; P = 0.012), and Phenix ×1 increased disagreement by 4.02° (P = 0.016), whereas CryoNet.Refine raised evaluation-half CC_mask_ (0.024; P = 0.023) without a detected *χ*_1_ change (Fig. 5). Deposited models may already have been built against both halves, so the evaluation half was withheld only from re-refinement.

**Figure 5.**
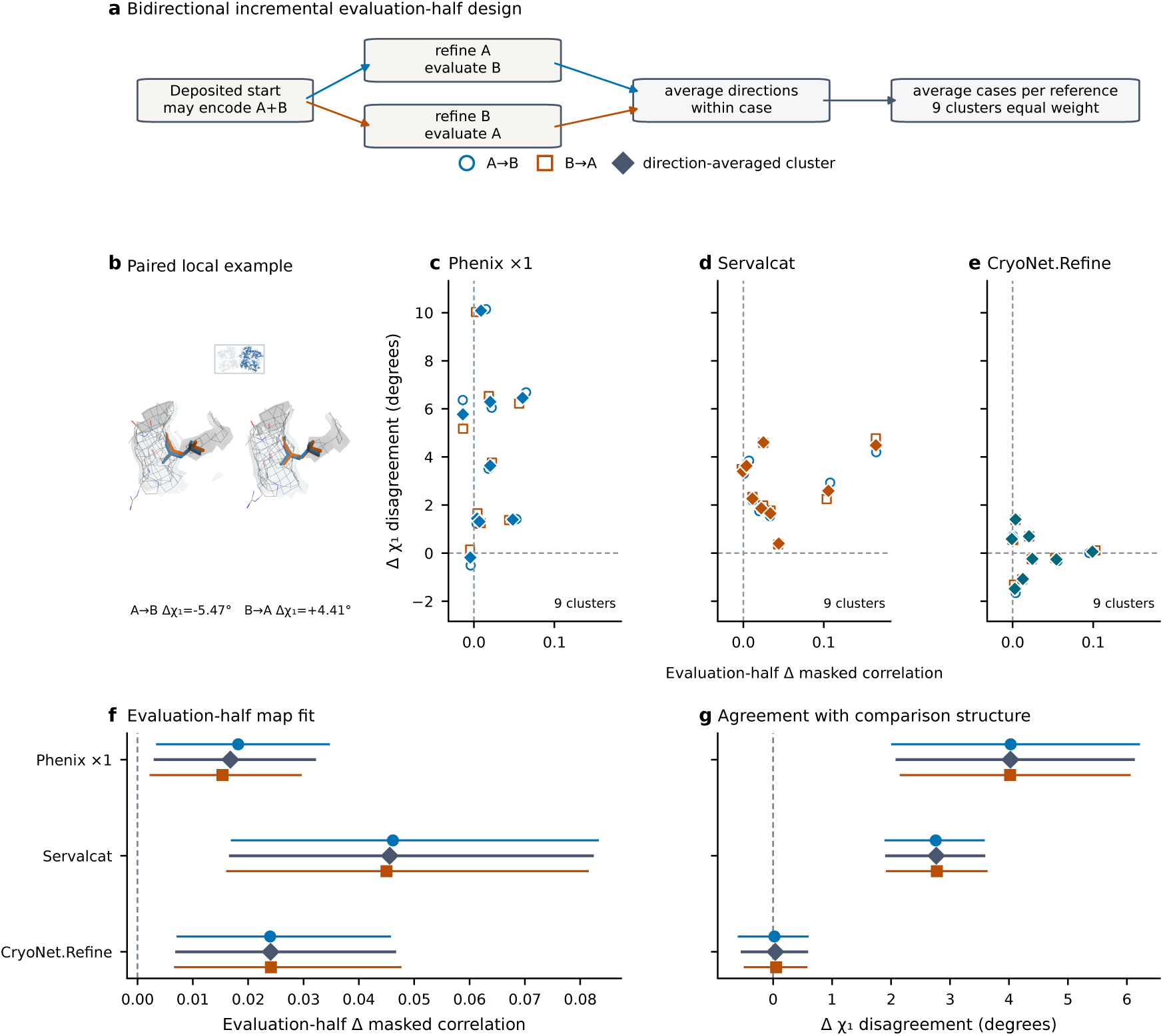
Opposite half maps in MEDIC. **a**, Refinement against one half map and evaluation on the other, in both directions, averaged within case. **b**, A rank-selected example (6B3Q LEU712; comparison 7K1F; Supplementary Fig. S3d) with the two evaluation halves. **c–e**, Directional and averaged values for nine clusters. **f,g**, Evaluation-half CC_mask_ change (**f**) and *χ*_1_ change (**g**) for each protocol in the A→B direction (circles), the B→A direction (squares) and averaged over both (diamonds), with 95% intervals.

For half-map selection¹⁵–¹⁷ in the earlier replication, each configuration’s evaluation-half CC_mask_ was averaged over both refinement directions before selection; its averaged *χ*_1_ disagreement supplied the outcome (Methods; Fig. 4c,d). As predicted (P6), selecting the action with the highest evaluation-half CC_mask_ increased disagreement by 1.83° (1.27–2.44°; 43 of 63 sequence groups worse), compared with 2.44° (1.85–3.07°) for work-half selection on the same 69 entries (one entry without valid Phenix half-map refinements is excluded from this rule). Selection using the evaluation half reduced the loss by about a quarter but did not eliminate it. Phenix’s gains in fit were smaller on the evaluation half (+0.004 and +0.005 for one and two passes, both intervals including zero), so the rule mostly rejected them; Servalcat retained a positive mean evaluation-half gain (+0.012, 0.007–0.018; 70 entries, 64 groups) while increasing disagreement by 2.07° (1.55–2.61°), and was still chosen in 43 of the 69 entries used for the selection rules. The direction and magnitude were similar with comparison structures as the clustering unit (1.82°, 1.26–2.41°; Supplementary Table S31).

### A reusable benchmark for refinement decisions

Primum scores refiners on fixed residue support and selection rules on case-level action tables (Extended Data Table 2). An example EMRinger rule calibrated on MEDIC and the earlier replication required a gain above 0.88 (rounded; exact value in Supplementary Table S38). In the 3.0–3.6 Å cohort, it accepted 107/167 entries (65.3% group-weighted coverage), with accepted-set loss −0.09° (−0.34 to +0.18°), overall loss −0.06° and regret 0.71° versus 3.18° for map-fit selection. It chose CryoNet.Refine 106 times and Servalcat once (Fig. 2f; Supplementary Table S38). Always CryoNet.Refine gave −0.09° at 100% coverage, with a 0.68° gap to the oracle. On the 53 within-study pairs, the unchanged calibrated EMRinger rule had accepted-set loss 1.00° at 45.3% coverage, failing both prespecified loss and coverage criteria (Supplementary Table S44).

## Discussion

Selection by support-matched same-map CC_mask_ increased side-chain disagreement at 3.0–3.6 Å, in the earlier cohorts and in pairs from the same study. In the earlier cohorts, the loss persisted without Servalcat and rose with its map weight. Evaluation-half correlation reduced the loss compared with work-half correlation, but did not eliminate it.

CryoNet.Refine improved mean map fit and mean side-chain agreement in the 156-entry replication, yet was rarely selected by the highest-CC_mask_ rule. The calibrated EMRinger rule mainly selected CryoNet.Refine at 3.0–3.6 Å and failed its loss and coverage criteria in the within-study test.

We propose five minimum reporting items for re-refinement benchmarks: (1) the no-operation endpoint value; (2) the candidate set and selection rule, with selection loss and observed regret; (3) coordinate agreement against a separately determined comparison, alongside map fit and geometry; (4) refinement coverage and the loss among accepted cases; and (5) the independent statistical units and scored residue support. Primum computes the decision summaries from a case-level table. Proposed acceptance rules should report coverage and loss on cases separate from those used for calibration. We recommend including no-operation comparisons in community model challenges, such as EMDataResource, and in validation reports that accompany re-refined models.

In the new within-study cohort screened before refinement for near-identical coordinates and recorded direct starting-model links, selection loss was 3.43° (2.85–4.02°). Within-study references are the depositors’ models of a second map, and state equivalence was assessed from archive metadata. The endpoints concern packed side chains; refinement used documented configurations without manual rebuilding, and map correlations used common side-chain atoms, fixed occupancy and B factors, and coordinate-derived masks (Methods; Supplementary Note 3).

## Online Methods

### Study design and estimand

The unit of analysis is a decision: for each case, a rule selects one action from a set that always contains the prepared deposited model (no operation). The primary estimand is the cluster-weighted mean change in chemicalsymmetry-aware *χ*_1_ disagreement of the selected action relative to no operation (selection loss). Observed regret is the selected action’s disagreement minus the smallest disagreement among the actions offered. Cases sharing a comparison structure (MEDIC) or a 95% sequence-identity group (both replication cohorts) are averaged before clusters receive equal weight. Within-study estimates weight pairs equally, with one pair per study. The configuration sweep, the two replication cohorts, the half-map selection analyses and both within-study cohorts were prespecified; post hoc analyses are labelled as such. Prespecification refers to protocols recorded internally before the corresponding analysis (Supplementary Note 8).

### Cohort at 3.0–3.6 Å

The cohort protocol and 200-entry roster were fixed before refinement. Single-particle entries released in 2014– 2023 with 3.0 ≤ *d* < 3.6 Å were selected using the replication rules below, with a comparison structure at least 0.5 Å better in resolution and released later. Sequence groups represented in MEDIC or in the full earlier replication roster, together with a further prespecified list of groups, were excluded. Entries were accepted in a fixed pseudo-random order derived from the entry identifier (Supplementary Note 8), at most two per 95% group. Preparation yielded 168 eligible entries in 132 groups. All five original actions were generated with the refinement commands of the earlier replication. Entry 8EAP had a map/model mismatch and no valid nooperation correlation, leaving 167 entries in 131 groups. Among the 103 entries with half maps, 6HBJ, 6TTF and 6WXH also lacked valid Phenix half-map results: both selection rules use 99 entries in 82 groups; individual Servalcat effects use 102 entries in 85 groups. Bootstrap intervals use 10,000 draws with seed 20261101. The roster and analysis membership are in Supplementary Table S35 and the technical record in Supplementary Note 8; the protocol document and archive query will be made available through the Primum repository (Code availability) and are available from the corresponding author on request.

### MEDIC cohort, residue support and endpoints

MEDIC validation pairs³⁰ provide a low-resolution deposited model, its map and a separately determined higherresolution comparison structure. One of 27 cases fails the prespecified minimum of 20 scored residues, leaving 26 cases and 20 clusters. The longest modelled chain of the starting model is globally aligned to each comparison chain and the chain with the most identical aligned residues is used. A residue is scored when it has the same residue type in both structures, a defined *χ*_1_ in both, and at least 18 starting-model C*α* atoms within 10 Å; the packing criterion is evaluated from the starting model alone, and a residue is reported only if every action in the analysis defines it. Circular angular distance is min(|*a* − *b*| mod 360°, 360°−(|*a* − *b*| mod 360°)); valine uses the minimum over CG1/CG2-equivalent definitions. Terminal atom-label alternatives account for LEU, ASP, PHE and TYR symmetry, while separate flip-aware alternatives are used for ASN and HIS; GLN uses a flip-aware heavy-atom comparison (Supplementary Table S1). GLU *χ*_2_ is not folded. Local side-chain RMSD uses a five-residue N, C*α*, C and O Kabsch frame⁴¹; canonical *χ*_1_ classes⁴²–⁴⁴ are assigned to the nearest of −60°, +60° and 180°. This three-class measure is distinct from full side-chain rotamer validation.

### Refinement protocols

Phenix 2.2-6143 real_space_refine used minimization_global+local_grid_search+nqh_flips with automatic weight for one pass (×1) and a second pass from its output (×2)¹. Servalcat 0.4.142 refine_spa_norefmac used the full map, ten conjugate-gradient cycles and automatic weight⁸. CryoNet.Refine used the official configuration (–recycles 300 –max_tokens 1000, seed 11, Phenix geostd monomer library)⁹; it writes mmCIF, which was converted to PDB with gemmi (read_structure, setup_entities, write_pdb) before scoring. Release-date sensitivities use the Boltz-2 pretraining cutoff; CryoNet.Refine’s refinement-training cutoff remains undocumented (Supplementary Table S40). KNexPHENIX, Phenix-after-MEDIC and the MEDIC-distributed Rosetta pipeline are included as contextual comparisons in Supplementary Table S26.

### Support-matched same-map CC_mask_

Within the target chain, heavy-atom identities (chain, residue, alternate location, atom name) are intersected across no operation, Phenix ×1, Phenix ×2, CryoNet.Refine and Servalcat; the side-chain atoms of *χ*_1_-scored residues form the primary support. Every other action must contain all of these atoms. Models are rewritten with occupancy 1.00 and B = 30 Å^2^ to isolate the coordinate contribution and scored with phenix.map_correlations at the map resolution. The voxel mask is generated separately from each model’s coordinates.

### Decision rules

The highest-CC_mask_ rule selects, within each case, the action with the largest support-matched CC_mask_; ties within 10^−12^ favour no operation, then the lexicographically first action. Action-set dependence re-applies the identical rule to reduced sets; a single-action acceptance rule offers one refiner and accepts it only when its own CC_mask_ gain exceeds 10^−12^, using the same no-operation tie tolerance. The post hoc application to the 3.0–3.6 Å cohort reports all four refiners on the same 167 entries (Supplementary Table S39). Supplementary Fig. S7f separately applies a descriptive rule to cluster-averaged gains and outcomes. Risk–coverage accepts the best refined action only above each observed gain threshold; the full surface is reported without choosing a threshold. EMRinger and twelve geometry or combined selectors (six geometry metrics, four structural metrics, two combined CC_mask_– geometry rules) were applied in the same way on the completed metric tables (EMRinger on the 24 cases at ≤4.5 Å).

### Residue-level summaries

For each rule, residues whose symmetry-aware *χ*_1_ disagreement with the comparison structure increased or decreased by more than 20° relative to no operation, and residues whose canonical *χ*_1_ class (nearest of −60°, +60° and 180°) gained or lost agreement with the comparison structure, were counted over the 6,480 scored residues and summarized as cluster-weighted fractions (10,000 bootstrap draws, seed 20261016). Per-protocol residue distributions are reported in Supplementary Tables S18 and S19. Post hoc summaries for the 3.0–3.6 Å cohort additionally report residue-level changes, *χ*_1_-class transitions, *χ*_2_ and local side-chain RMSD on common support, with 10,000 group-bootstrap draws and seed 20261101 (Supplementary Table S37). No P values were computed for these summaries.

### Configuration sweep

From each deposited model we ran, with every unlisted option at its original value (Supplementary Table S28): Servalcat with –jellybody; –weight 0.5× or 2× the per-case automatic weight recorded in the original Servalcat log with –no_weight_adjust; –adp fix; and –halfmaps instead of –map in the ten cases with deposited half maps; and Phenix with ADP refinement added, reference-model restraints to the starting model, target bond and angle r.m.s.d. 0.005 Å and 0.5°, and morphing plus simulated annealing. A run was technically valid if it exited normally, wrote finite coordinates and contained every support atom and every *χ*_1_ atom of the scored residues; a configuration was included only if all its runs were valid. One simulated-annealing run (6XH8) exceeded the 4-h limit, excluding that configuration; the other 192 runs were valid. Prespecified analyses were per-configuration changes, the highest-CC_mask_ rule on the full set and without each program family, within-family rules, the relation between map-fit gain and *χ*_1_ change across configurations and the half-map subset (Supplementary Table S29), with the criterion for a loss persisting beyond Servalcat met only if both the full-set and the without-Servalcat intervals lay entirely above zero.

### Comparison-structure local support

Per-residue support was computed from the comparison data alone: native map–model correlation for 16 cryoEM comparison structures on their full maps, and real-space correlation against observed data for four X-ray structures (alternate conformers by highest occupancy). Residues were assigned to within-structure quartiles by midrank percentile; cryo-EM and X-ray values were not pooled.

### Bidirectional half maps (MEDIC)

Ten cases were refined against one half map and evaluated on the other in both directions; the primary fit endpoint is evaluation-half CC_mask_ on the *χ*_1_ side-chain support at B = 30 Å^2^, with B = 10, 20, 50 and 80 Å^2^ as sensitivities. Directions are averaged within case before clustering.

### Independent replication cohort

The protocol, including predictions P1–P4, was specified before cohort selection. Candidates were queried once from the RCSB and EMDB archives: electron microscopy, single-particle reconstruction, 3.6–6.5 Å, initial release 2014–2023, no nucleic-acid polymer, at most 4,000 deposited residues, one primary map of at most 512 MiB and legacy PDB-format coordinates; the target entity is the longest protein entity (≥100 residues). A comparison structure had to be X-ray or electron microscopy, ≤3.2 Å and at least 1.0 Å better, contain an entity in the same RCSB 95% sequence-identity group, be released strictly after the low-resolution entry and have PDB-format coordinates; the best-resolved such entry was chosen (ties: earliest release, then identifier). Groups containing any MEDIC entity were excluded. Entries were taken in a fixed pseudo-random order derived from the entry identifier (Supplementary Note 8) and accepted up to two per group, 200 entries or an 80-GB download budget; 2,758 search hits gave 1,237 eligible pairs and 200 accepted entries in 164 groups. Starting models were reduced to the 20 standard amino acids (selenomethionine to methionine), first model, highest-occupancy alternate location, no hydrogens, waters, ions or ligands; this prepared model is the no-operation action and the input to every arm. Eligibility followed MEDIC (≥95% aligned identity to the comparison chain, ≥20 packed *χ*_1_ residues), leaving 156 entries in 130 groups. Protocols were the prespecified Phenix ×1/×2, Servalcat and CryoNet.Refine protocols, the eight sweep configurations and Servalcat with half-map input where two half maps were listed. In this cohort ADP refinement left coordinates identical to Phenix ×1 in every entry (only B factors, which are fixed for scoring, differed), so the two arms have identical values. A case was excluded from any analysis containing an arm that failed its technical checks; a configuration failing in more than 10% of cases was excluded. The 95% sequence group is the independent unit (sensitivity: comparison structure); intervals use 10,000 draws with seed 20261015.

### Half-map selection

For the 70 eligible replication entries listing two half maps, Phenix ×1/×2 and Servalcat were refined from the prepared model against each half (directions A and B). One entry lacked valid Phenix half-map refinements; Phenix estimates and both selection rules therefore use 69 entries (63 sequence groups), and Servalcat estimates use all 70 entries (64 groups). Each model was scored on the opposite (evaluation) half and on its own (work) half with the same support rule; no operation was scored on both halves. In both replication cohorts, each configuration’s work-half and evaluation-half scores and *χ*_1_ disagreement are first averaged over the two directions. The rule then selects one configuration per entry using its averaged score and reports its averaged disagreement relative to no operation. Selected-action counts therefore count configurations per entry. The prespecified prediction P6 is positive loss for selecting the highest evaluation-half CC_mask_ among no operation, Phenix ×1, Phenix ×2 and Servalcat; work-half selection on the same cases is reported for comparison.

### Within-study low/high-resolution pairs

The 22-pair pilot motivated a further within-study cohort whose protocol and 90-study roster were fixed before downloading coordinates or maps. Pairs came from a metadata search for shared study and reconstruction relationships, with 3.0 ≤ *d*_low_ < 6.5 Å, *d*_high_ ≤ 3.5 Å and a gap of at least 0.5 Å. Automatic checks compared title state labels, major ligands and composite-map status. Pilot studies and entries were excluded; one pair per study was selected by shared modelled fraction, resolution gap, reference resolution and identifiers, in that order. Starting models contained at most 60,000 atoms, with at least half their modelled residues present in the reference. Before refinement, pairs required downloadable coordinates and maps, at least 95% sequence identity, at least 20 packed *χ*_1_ residues, scored-residue C*α* RMSD of 0.1–3.0 Å, baseline *χ*_1_ disagreement of at least 12.95° and no recorded direct starting-model link. These criteria retained 53 pairs (52 separate acquisitions, one shared dataset); 44 had usable half maps. Acquisition relationships and state screening were assessed from archive metadata. The five actions used the protocols above; the half-map rule refined against half 1 and selected by half 2. Intervals used 10,000 pair-bootstrap draws, seed 20261101; 95% sequence groups formed a sensitivity analysis. Protocols, exclusions and pilot analyses are in Supplementary Tables S41–S44 and Supplementary Note 8.

### External cohorts

The developer-generated AQuaRef public cohort (41 cases, 23 connected reference components; 40 under the primary mapping rule)¹⁰ and 19 CERES service results⁶ were analysed as separate tracks with their own baselines, endpoints and multiplicity families (Supplementary Tables S14–S17, S20 and S21; Supplementary Figs. S5 and S6; Supplementary Notes 2 and 5); they are not pooled with MEDIC or the replication cohort.

### Statistics

Intervals are unadjusted 95% percentile bootstrap intervals that resample the independent unit (reference cluster, 95% sequence group, within-study pair or AQuaRef reference component) with replacement⁴⁵. Bootstrap settings were fixed within each analysis protocol and differ between analysis stages: the original endpoint analyses (Supplementary Table S33, Fig. 5f,g, Extended Data Figs. 1, 2, 6 and 7, Supplementary Figs. S4 and S5 and the corresponding Supplementary Tables) used 8,000 draws with seed 0; the AQuaRef analyses used 10,000 draws with seed 0; and later analyses used 10,000 draws with protocol-specific seeds (for example, 20260902 for the original decision analyses, 20261014 for the sweep and 20261015 for the earlier replication and its half-map selection; 20261101 for the 3.0–3.6 Å cohort, both within-study cohorts and calibrated acceptance analyses). The same five-action selection therefore has slightly different intervals in the original analysis (1.85–3.74°) and the sweep (1.89–3.74°). For signed protocol effects, two-sided Wilcoxon signed-rank tests on cluster values⁴⁶ were Holm-adjusted within each stated family (Supplementary Table S33: 7 arms; CC_mask_: 4; half maps: 3 per endpoint; AQuaRef: 6 per endpoint and rule; CERES: 8)⁴⁷; no P values were computed for any decision analysis or for the 3.0–3.6 Å and within-study cohorts. Leave-oneand leave-two-cluster deletions were used to assess the influence of individual clusters.

### Calibration of an example acceptance rule

The rule family and thresholds were fixed on 26 MEDIC and 156 earlier replication entries (149 sequence groups), before scoring the 3.0–3.6 Å cohort. Candidate rules used movement caps, EMRinger, Q-score, masked FSC and combined geometry vetoes; a logistic rule was secondary. Calibration selected the largest target coverage in 0.30–0.80 with a one-sided 90% bootstrap upper bound on accepted-set loss at or below zero, breaking ties by regret and parameter count. The selected rule takes the refined candidate with the highest full-map EMRinger score²¹ and accepts it when its gain over no operation exceeds the calibrated threshold (0.88 when rounded; exact value in Supplementary Table S38); otherwise it retains no operation. Unavailable candidate scores are ignored; an unavailable baseline score yields no operation. Tied candidate scores use lexicographic program order. Application uses the map and candidate models. Thresholds were unchanged when applied to this cohort. Accepted-set loss divides group-weighted accepted loss by group-weighted coverage; overall loss also includes retained no-operation choices. Supplementary Table S38 reports calibration, test results and the prespecified test criteria.

### Primum

Primum (Python, NumPy only) implements the decision analysis (selection within each case, selection loss, observed regret, leave-one-action-out sets, single-action acceptance rules, risk–coverage and the cluster bootstrap) for any long table of case, cluster, action, selection score and outcome, and the coordinate endpoint using the same functions as the analysis code; CC_mask_ calls phenix.map_correlations when available. Its test suite reproduces every entry of Extended Data Table 1 from the case-level analysis table to within 10^−9^.

Primum v1.0 identifies the original cohort, endpoint and selection specifications; v2 adds the 3.0–3.6 Å cohort, the 22-pair pilot and the prespecified 53-pair within-study cohort using the same coordinate endpoint. Each case was indexed by its starting and comparison structures, chain correspondence, scored residue support and independent unit. Scoring used the same retained support across valid actions within each analysis; technical validity and candidate availability were recorded separately. The earlier 200-entry replication roster identifies the 156 eligible entries; the separate 3.0–3.6 Å cohort list identifies 168 eligible and 167 analysed entries. The cohortselection record specifies the archive query, release window, 95% sequence groups, exclusions, entry ordering and acceptance limits. These settings, together with the input identifiers, define the current cohort version and the selection rule for its reuse.

## Data availability

Coordinates, maps and half maps are public in the PDB and EMDB^48–49^. The 26 analysed low-resolution deposited models are 3JD3, 5GJW, 5YWD, 6B3Q, 6B70, 6DMR, 6F36, 6HRB, 6M66, 6M68, 6PBE, 6PBF, 6PVM, 6PVP, 6RNY, 6S5T, 6T24, 6T7S, 6WBI, 6X17, 6XH7, 6XH8, 7CCS, 7E5Z, 7K3A, 7L2J. The 20 comparison structures are 5K12, 6JPA, 6RD5, 6TA5, 6W7V, 6WBF, 6XL5, 7ADU, 7BGY, 7DWB, 7K1F, 7LP9, 7MIJ, 7MIR, 7P9V, 7QA4, 7RCP, 7T6O, 7VW6, 7W4O. The 26 corresponding EMDB maps are EMD-0258, EMD-4176, EMD-4960, EMD-6632, EMD-6853, EMD-7041, EMD-7062, EMD-7965, EMD-9515, EMD-10100, EMD-10366, EMD-10371, EMD-20291, EMD-20292, EMD-20493, EMD-20496, EMD-21590, EMD-21991, EMD-22184, EMD-22185, EMD-22654, EMD-23130, EMD-30114, EMD-30116, EMD-30341, EMD-30995.

The 200 replication entries, their comparison structures, maps and selection funnel are listed in Supplementary Table S27 (Supplementary Data 1); the 3.0–3.6 Å cohort list and analysis membership are in Supplementary Table S35 (Supplementary Data 2); the within-study rosters and analysis membership are in Supplementary Table S43. Supplementary Data 3 contains the within-study case-level results and exclusions; Supplementary Data 4 contains study identifiers, map identifiers and reconstruction metadata. Cohort protocols will be made available through the Primum repository. The Primum data-access entry point is the repository listed under Code availability. Dataset documentation and links to source tables, starting and comparison coordinates, re-refined models, residue supports, analysis protocols will be indexed there. These materials are available from the corresponding author on request and will be publicly archived on publication. Licensed third-party software and model weights are not redistributed.

## Code availability

The Primum software repository is https://github.com/INTOSYN/primum. Primum and the analysis code will be released under an open-source licence on publication and are available from the corresponding author on request. The repository will link each software version to its data inventory, scoring protocol and reference results.

## Acknowledgments

Computations were performed on INTOSYN AI Labs infrastructure.

---

**Extended Data Fig. 1.**
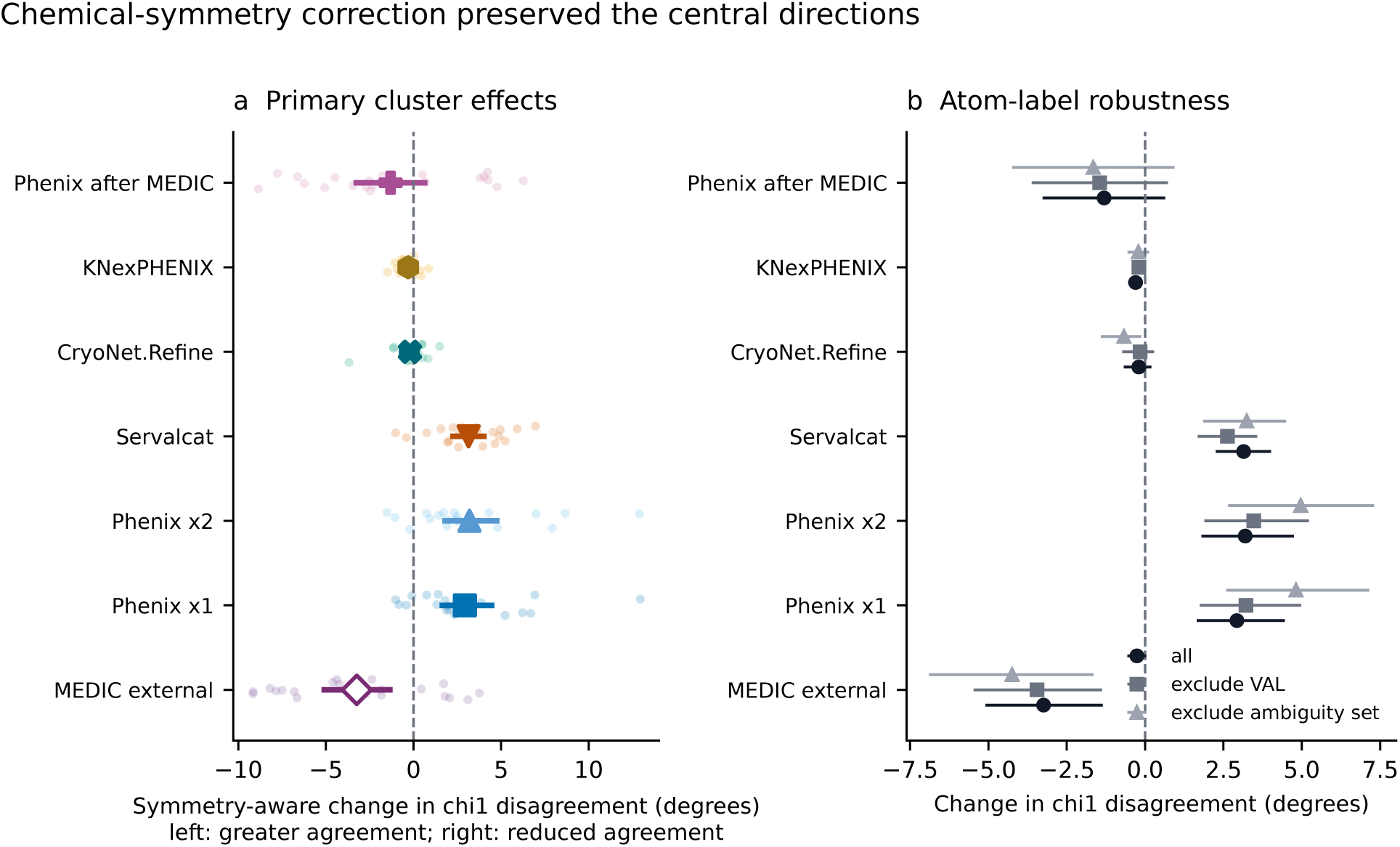
Chemical-symmetry and atom-label robustness. Primary, valine-excluded and ambiguity-set-excluded *χ*_1_ changes for the seven MEDIC arms; cluster-weighted means across 20 clusters with unadjusted 95% cluster-bootstrap intervals. Contextual arms are distinguished by their input and provenance (Supplementary Table S26).

**Extended Data Fig. 2.**
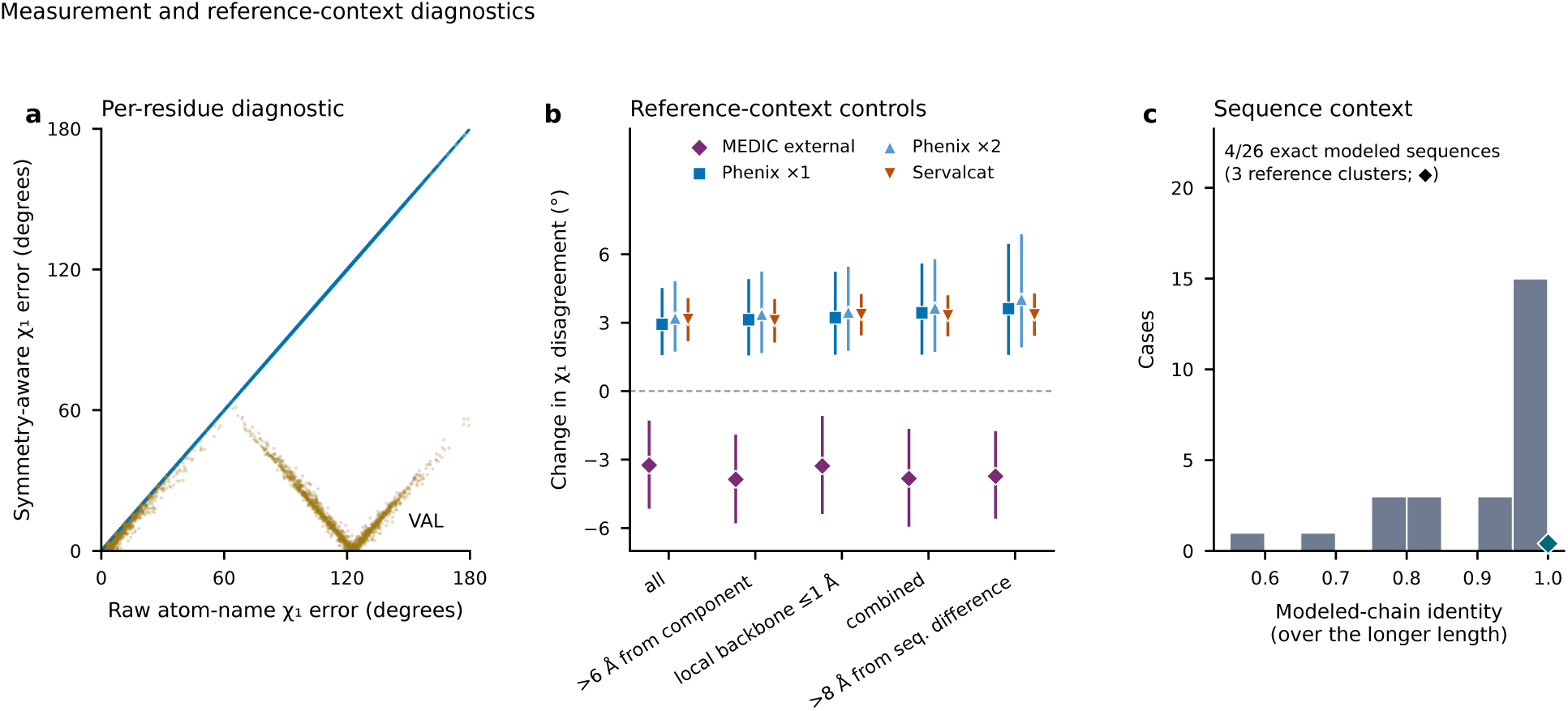
Measurement and reference-context diagnostics. Raw versus symmetry-aware *χ*_1_; exclusion of residues near non-polymer components, with poor local backbone agreement or near sequence differences; and two chain-identity denominators. The diagnostics assess sensitivity to residue matching and local structural context.

**Extended Data Fig. 3.**
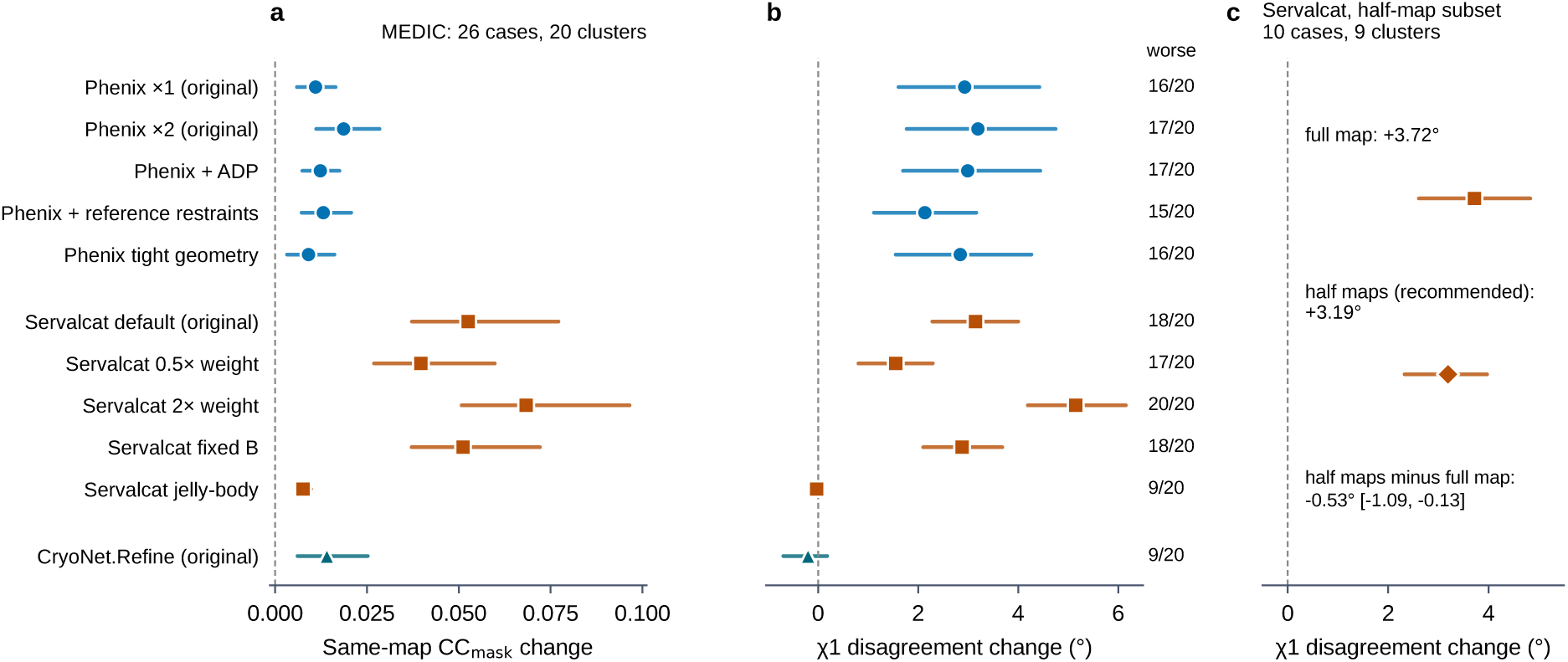
Every refinement configuration of the sweep. **a**, Same-map CC_mask_ change and **b**, *χ*_1_ change relative to no operation for the 11 refinement configurations (MEDIC, 26 cases, 20 clusters), with the number of clusters worse than no operation; cluster-weighted means and 95% cluster-bootstrap intervals. **c**, Servalcat with its recommended half-map input and with the full map in the ten cases with deposited half maps. Numerical values are in Supplementary Table S29.

**Extended Data Fig. 4.**
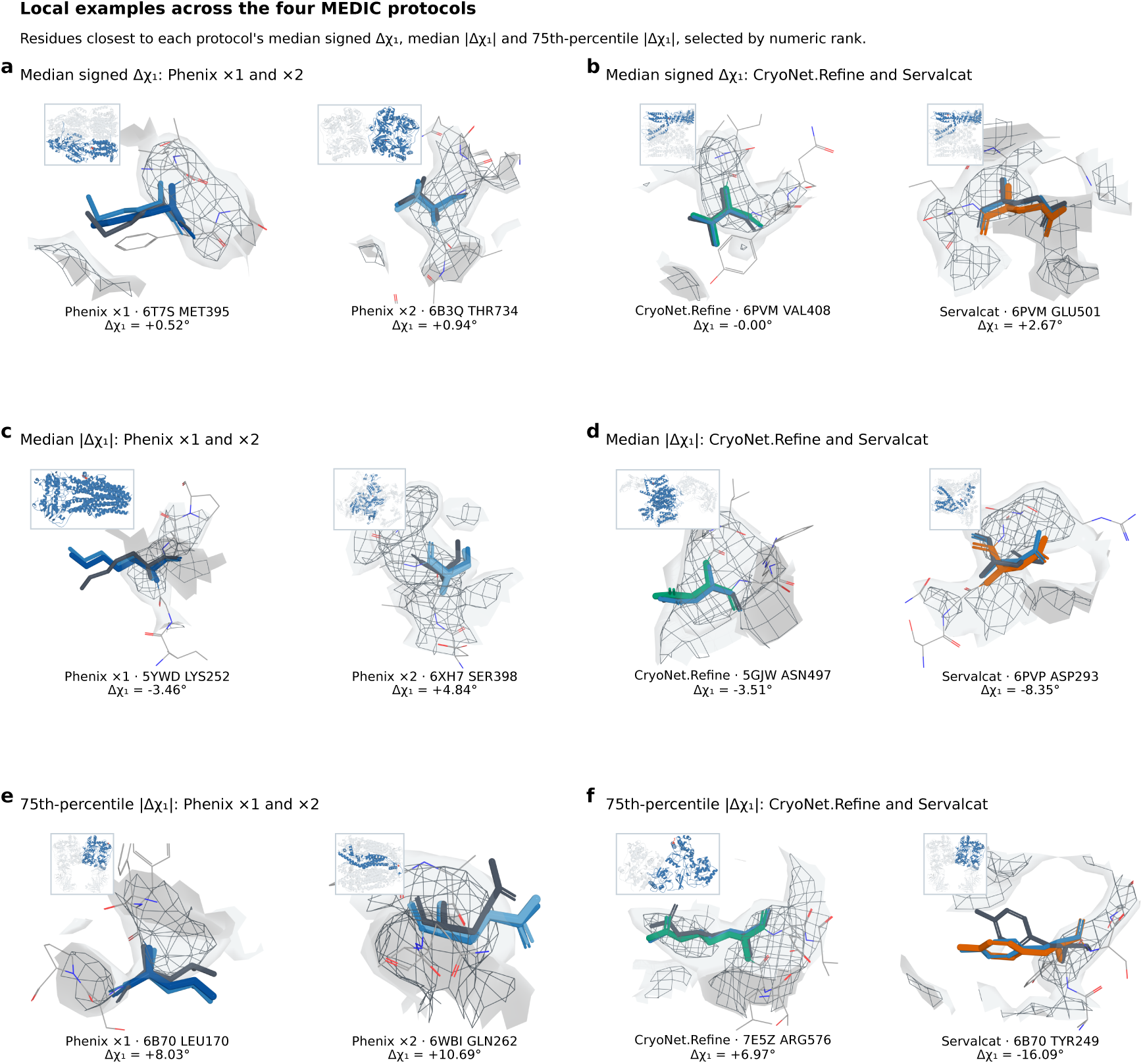
Molecular examples across the four MEDIC protocols. **a,b**, Residues closest to each protocol’s median signed Δ*χ*_1_, paired as Phenix ×1/×2 and CryoNet.Refine/Servalcat; **c,d**, residues closest to the median absolute Δ*χ*_1_; **e,f**, residues closest to the 75th-percentile absolute Δ*χ*_1_. Each cell shows the starting residue (blue), the refined residue (method colour), the superposed comparison residue (charcoal; five-residue Kabsch frame), the deposited map and a whole-model locator. Values are signed Δ*χ*_1_ relative to no operation (positive, reduced agreement). Examples were selected by numeric rank (Extended Data Fig. 8).

**Extended Data Fig. 5.**
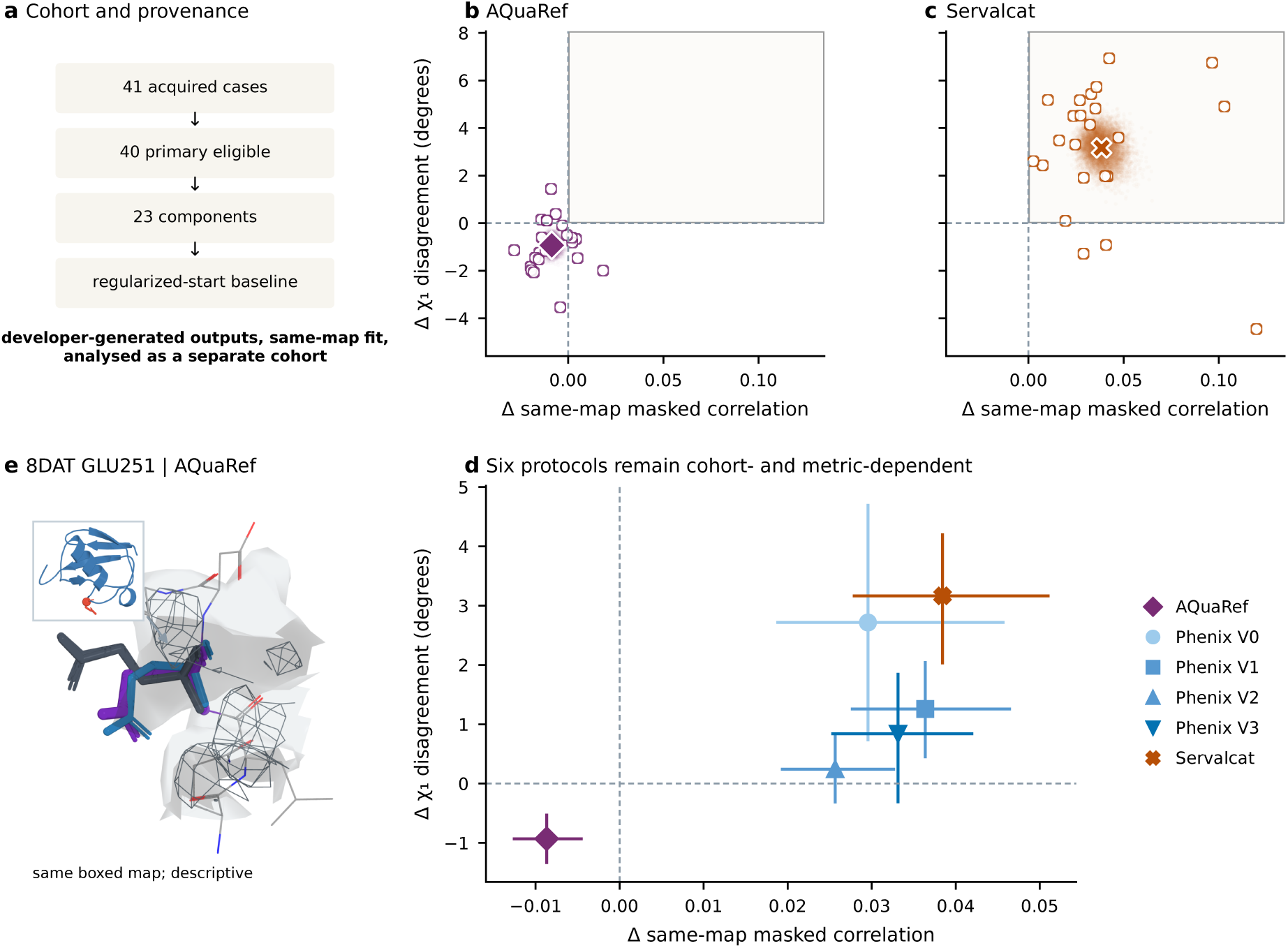
A developer-generated external cohort with the opposite combination. **a**, The public AQuaRef cohort retains 40 of 41 cases in 23 connected reference components on a regularized-start baseline. **b,c**, All 23 component values for AQuaRef and public Servalcat; large symbols are equal-component means and pale clouds paired bootstrap means. **d**, Six refinement configurations with 95% component-bootstrap intervals; the baseline and residue support are shared within this cohort. **e**, Example 8DAT GLU251 in the refinement map. Same-map, coordinate-derived CC_mask_; developer-generated outputs, analysed separately from the other cohorts.

**Extended Data Fig. 6.**
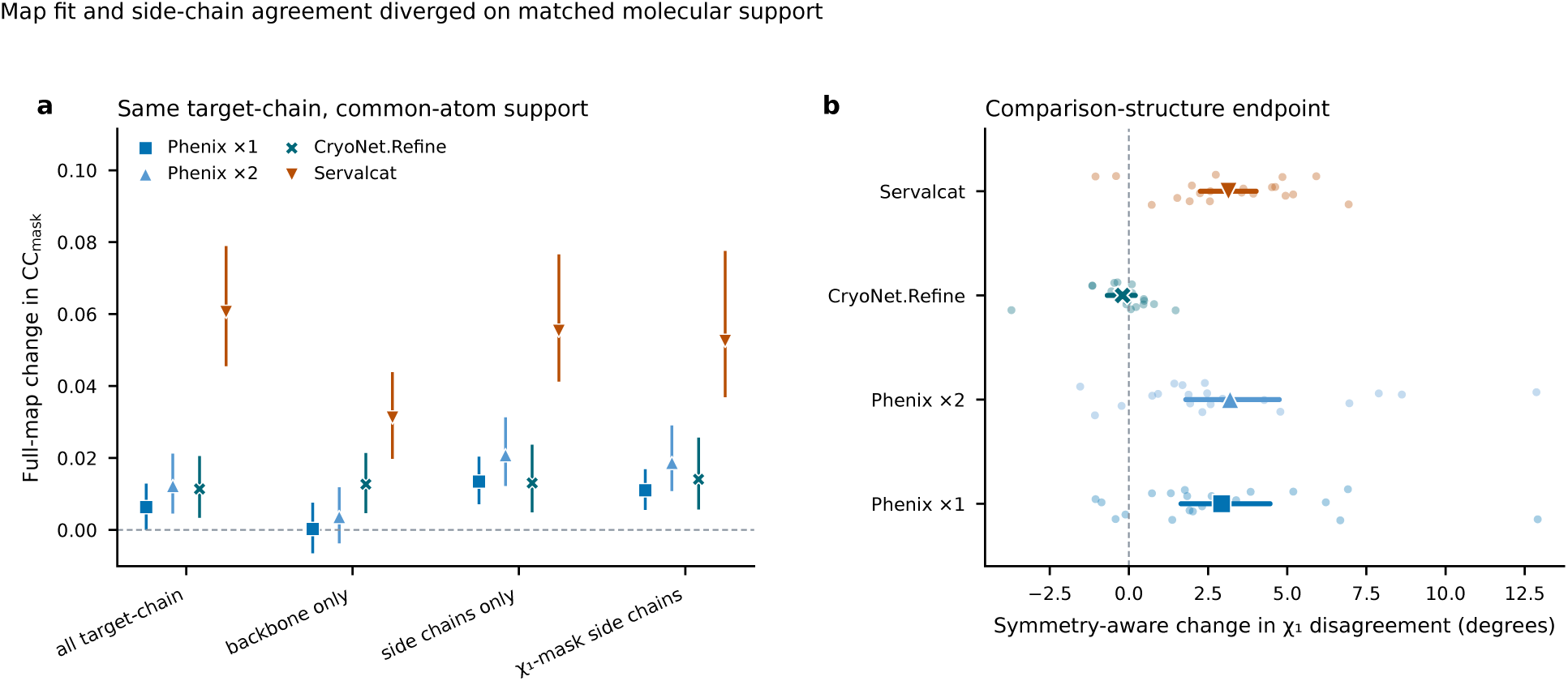
Same-map CC_mask_ across atom subsets. Target-chain, backbone, side-chain and *χ*_1_residue side-chain supports for the four MEDIC protocols. The mask is derived from each model’s coordinates.

**Extended Data Fig. 7.**
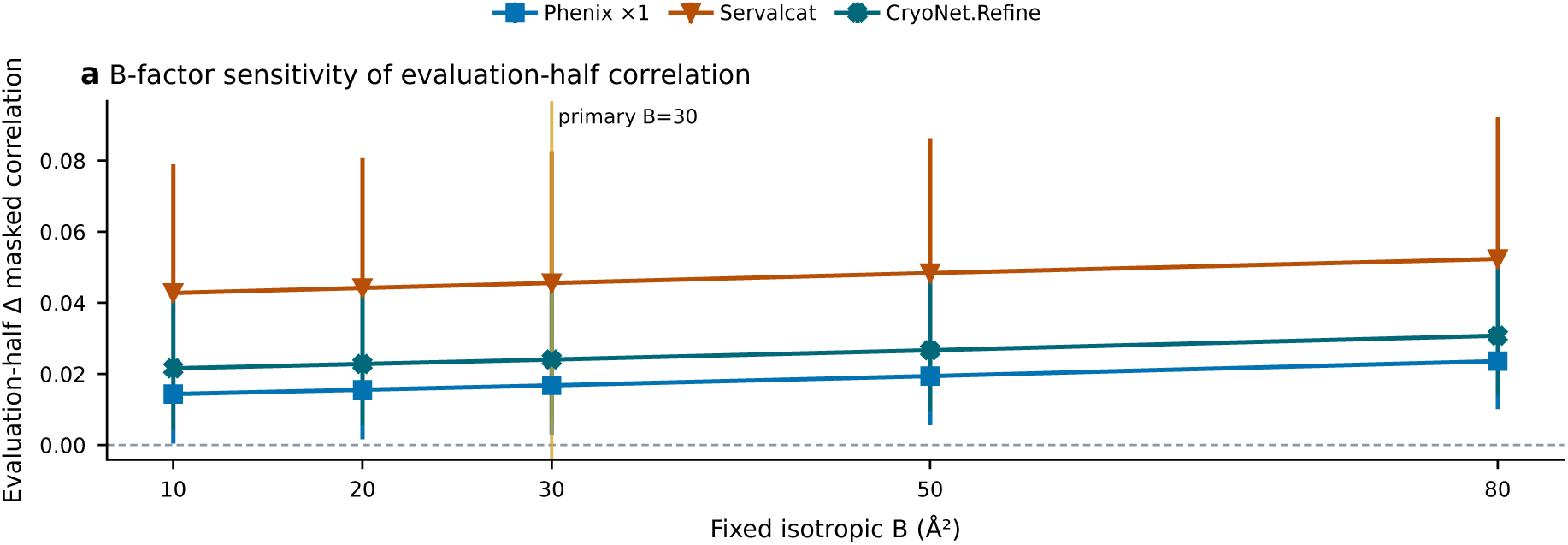
Half-map fixed-B sensitivity. Evaluation-half CC_mask_ changes at B = 10, 20, 30, 50 and 80 Å^2^ for three protocols (B = 30 Å^2^ primary); cluster-weighted means with unadjusted 95% intervals.

**Extended Data Fig. 8.**
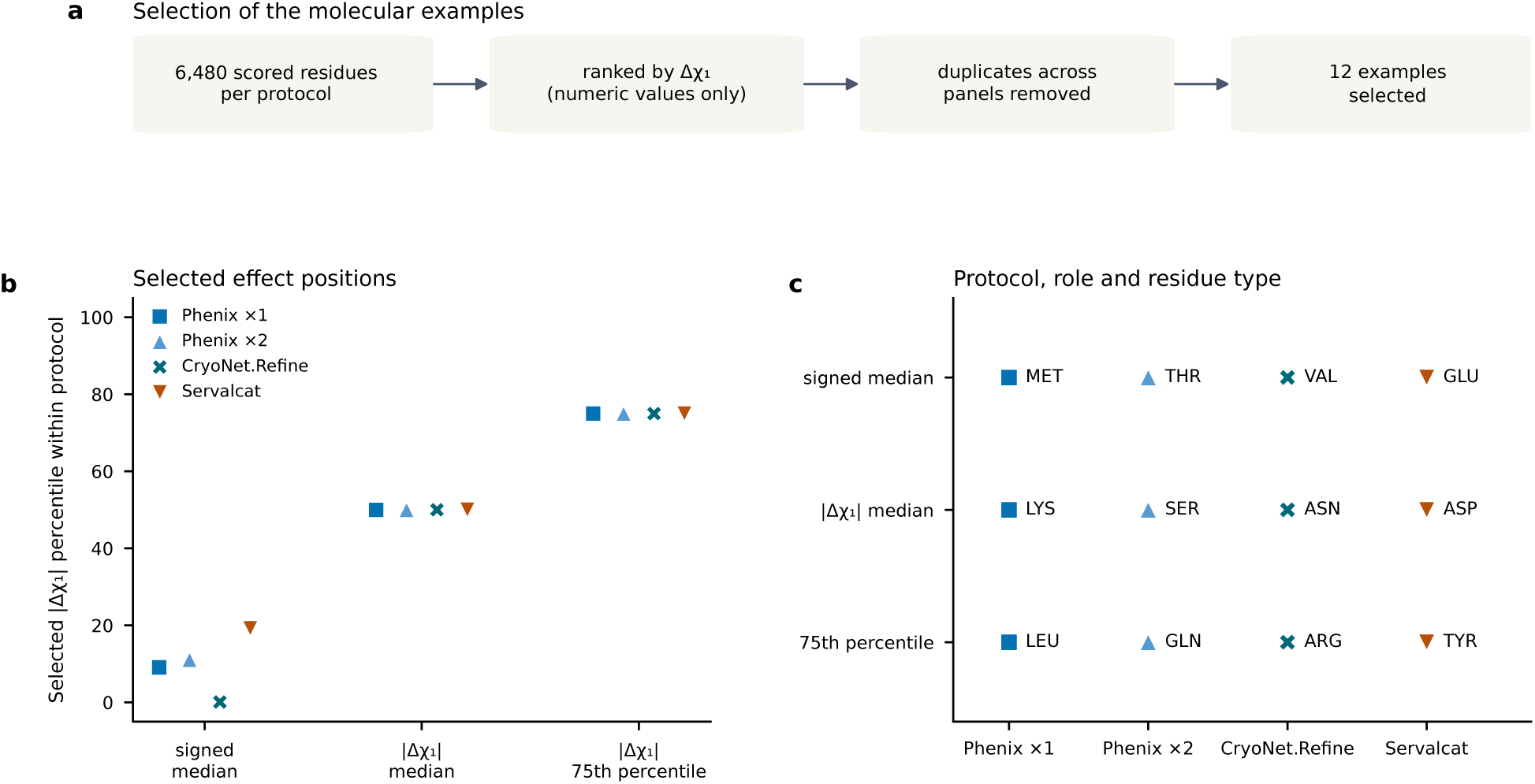
Selection of the molecular examples. **a**, Twelve examples were selected from the 6,480 scored residues by numeric ranking. **b**, Percentiles of the selected absolute effects within each protocol. **c**, Protocol, selection role and residue type of each example (each the top-ranked candidate).

**Extended Data Fig. 9.**
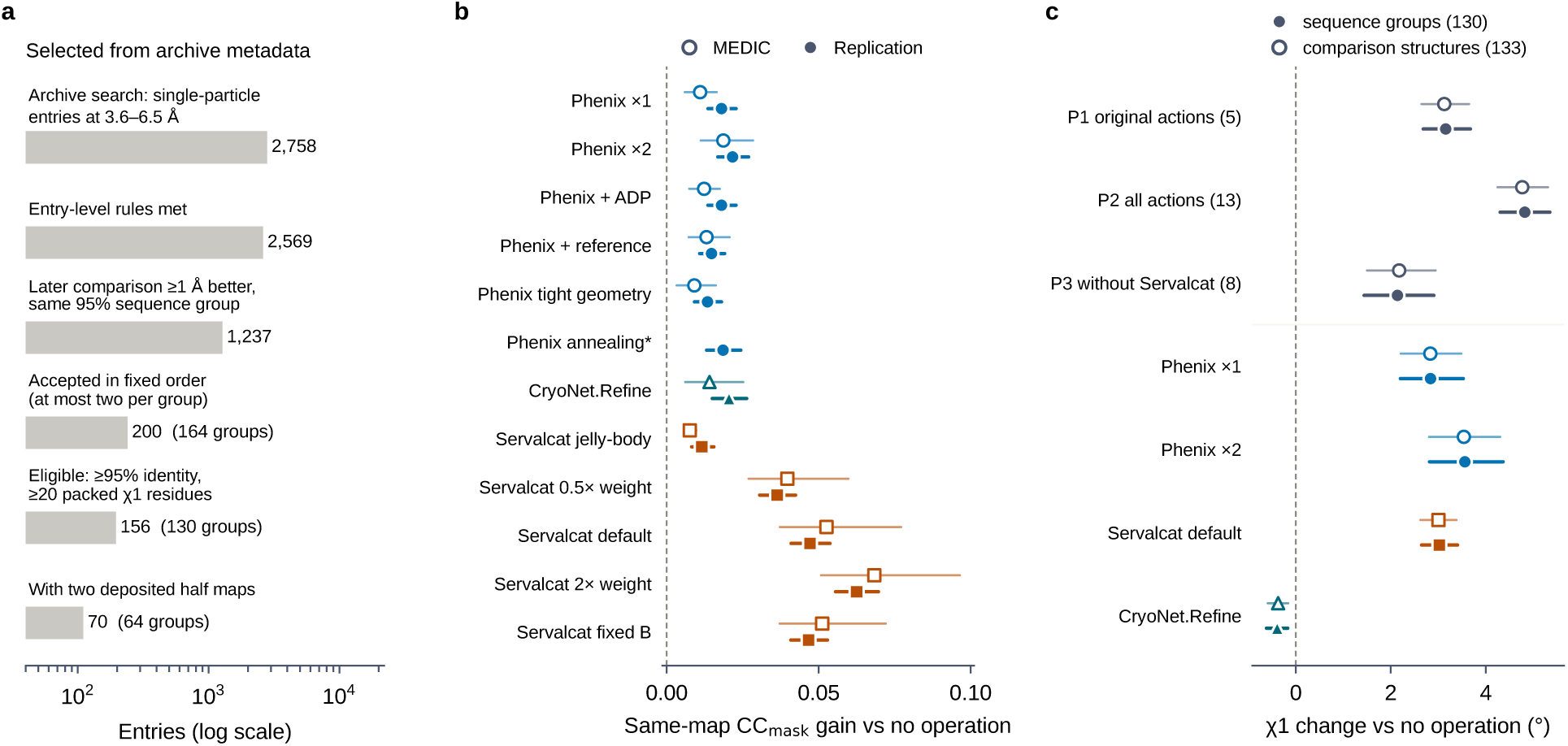
The replication cohort. **a**, Selection from archive metadata; of the 200 accepted entries, 35 had less than 95% aligned sequence identity to their comparison structure and 9 had fewer than 20 packed *χ*_1_ residues after preparation. **b**, Same-map CC_mask_ gain relative to no operation for every refinement configuration, replication (filled) beside MEDIC (open). **c**, Selection loss for P1–P3 and *χ*_1_ changes of the four original programs with sequence groups (filled; primary unit) or comparison structures (open) as the clustering unit. Cluster-weighted means with 95% cluster-bootstrap intervals.

**Extended Data Fig. 10.**
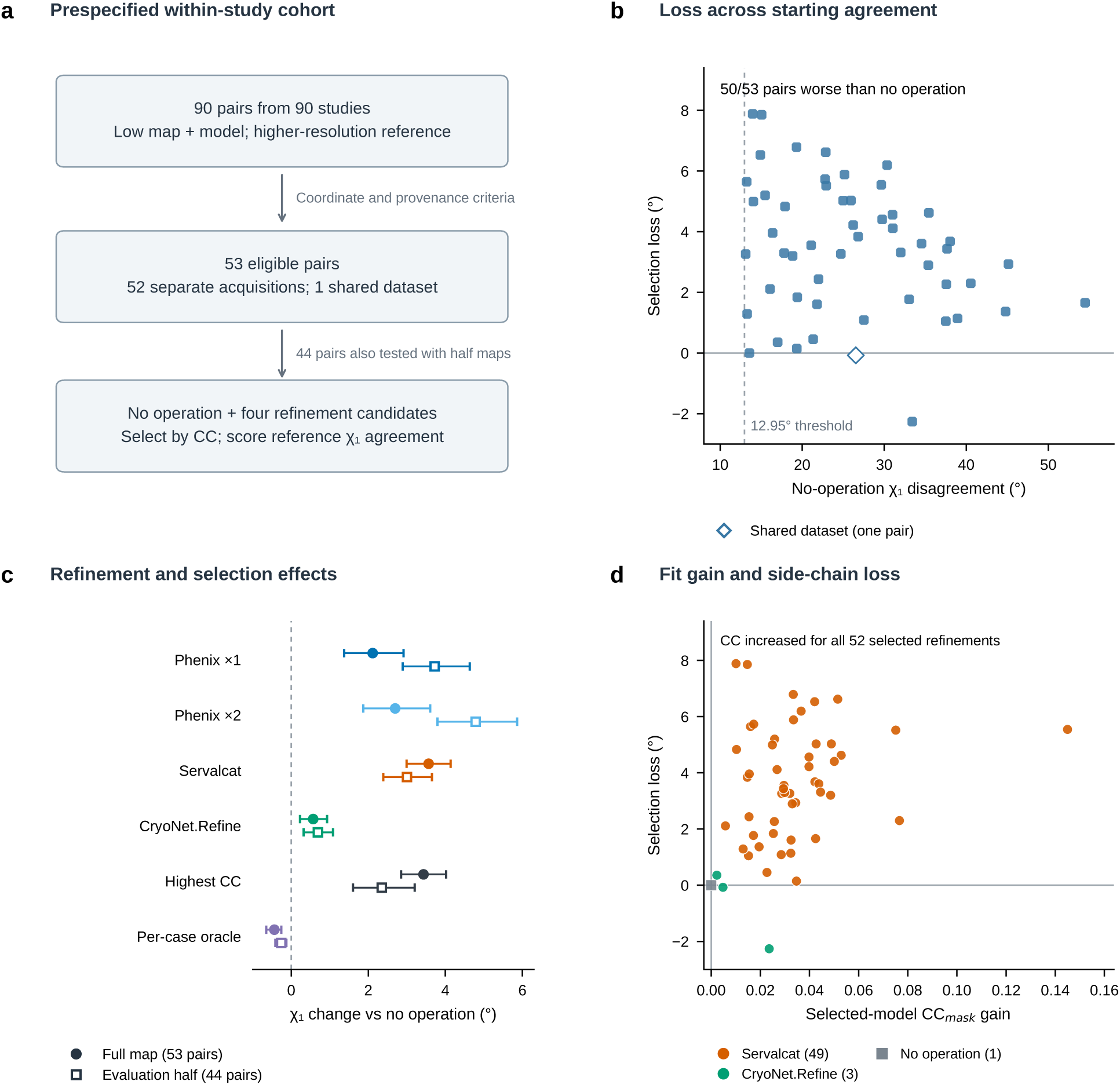
Selection loss in the prespecified within-study cohort. **a**, Cohort and five-action design. Screening before refinement retained 53 of 90 pairs, with 52 separate-acquisition pairs and one shared-dataset pair; 44 pairs had usable half maps. **b**, Full-map selection loss against baseline *χ*_1_ disagreement for all 53 pairs. The vertical dashed line marks the prespecified 12.95° baseline threshold; the diamond identifies the shared-dataset pair. **c**, Individual refinement effects, highest-CC_mask_ selection and the per-case oracle. Filled circles indicate full-map results (53 pairs), open squares indicate evaluation-half results after refinement against half 1 (44 pairs). Bars are 95% pair-bootstrap intervals (10,000 draws, seed 20261101). **d**, Full-map CC_mask_ gain and selection loss for the selected action in each pair. All 52 selected refinements increased CC_mask_; one pair retained no operation. Colour identifies the selected action. Positive selection loss is increased *χ*_1_ disagreement relative to no operation. Pair identifiers, baseline and provenance checks, exclusions and fixed-rule evaluation are in Supplementary Tables S41–S44.

**Extended Data Table 1.** Action-set dependence of the original five-action rule. The identical selection rule was re-applied after removing one refiner from the five original actions. Values are cluster-weighted mean *χ*_1_ selection loss relative to no operation with percentile cluster-bootstrap intervals (10,000 draws, seed 20260902). “Clusters worse” is the number of the 20 clusters with a positive estimate.

| Action set | Loss (°) | 95% interval | Clusters worse | Selected actions |
|---|---|---|---|---|
| All four refiners (MEDIC) | +2.83 | [+1.85, +3.74] | 17/20 | Servalcat 25, Phenix ×2 1 |
| Drop Phenix ×1 | +2.83 | [+1.85, +3.74] | 17/20 | Servalcat 25, Phenix ×2 1 |
| Drop Phenix ×2 | +3.15 | [+2.26, +4.01] | 18/20 | Servalcat 26 |
| Drop CryoNet.Refine | +2.83 | [+1.85, +3.74] | 17/20 | Servalcat 25, Phenix ×2 1 |
| **Drop Servalcat** | **+2.13** | **[+0.68, +3.85]** | **11/20** | Phenix ×2 14, CryoNet 10, Phenix ×1 2 |

**Extended Data Table 2.** Primum reference strategies at 3.0–3.6 Å. Loss is the change in *χ*_1_ disagreement relative to no operation; lower values indicate better agreement. The no-operation endpoint is 27.16°. Each block fixes its candidate set and evaluation sample. Earlier MEDIC and replication results are retained in Extended Data Table 1 and Supplementary Table S34.

| Strategy | Loss [95% interval] (°) | Groups worse | Coverage (%) | Regret (°) |
|---|---|---|---|---|
| **a, Full-map selection: five actions, 167 entries and 131 sequence groups** |  |  |  |  |
| No operation | 0.00 [0.00, 0.00] | 0/131 | 0.0 | 0.77 |
| Highest full-map CC<sub>mask</sub> | +2.41 [+2.12, +2.71] | 119/131 | 96.2 | 3.18 |
| Always CryoNet.Refine | −0.09 [−0.31, +0.13] | 65/131 | 100.0 | 0.68 |
| Per-case oracle | −0.77 [−0.91, −0.63] | 0/131 | 64.5 | 0.00 |
| **b, Half-map subset: four actions, 99 entries and 82 groups** |  |  |  |  |
| Highest work-half CC | +2.89 [+2.35, +3.51] | 74/82 | — | — |
| Highest evaluation-half CC | +2.29 [+1.70, +2.97] | 56/82 | — | — |

Group-weighted means and 95% percentile bootstrap intervals use 10,000 draws and seed 20261101. Coverage is the group-weighted fraction assigned a refinement. Regret is measured against the per-case minimum within the candidate set. The oracle uses the observed reference outcome to give a hindsight lower bound. An em dash denotes an unreported value. Full-map candidates are no operation, Phenix ×1/×2, Servalcat and CryoNet.Refine; half-map candidates exclude CryoNet.Refine.

