## Supplementary Information for "Primum: a decision benchmark for selecting cryo-EM re-refinements"

### Supplementary Table S1 — Symmetry and flip handling

| endpoint/residue | comparison convention |
|---|---|
| VAL $\chi_1$ | minimum over CG1/CG2 definitions |
| LEU $\chi_2$ | minimum over CD1/CD2 definitions |
| ASP $\chi_2$ | minimum over OD1/OD2 definitions |
| PHE/TYR $\chi_2$ | ring-label equivalent definitions |
| ASN $\chi_2$ | OD1/ND2 flip-aware comparison |
| HIS $\chi_2$ | ND1/CD2 alternative; heavy-atom comparison also maps CE1/NE2 |
| GLN heavy atoms | terminal OE1/NE2 flip-aware heavy-atom mapping; $\chi_2$ is not folded |
| GLU | $\chi_2$ is not 180° folded because terminal carboxylate symmetry is at $\chi_3$ |
| canonical $\chi_1$ class | nearest $\chi_1$ center among −60°, +60° and 180° |

### Supplementary Table S2 — Symmetry and atom-name robustness

| arm | all $\Delta\chi_1$ | Holm-adjusted P | no-VAL $\Delta\chi_1$ | Holm-adjusted P | no-ambiguity $\Delta\chi_1$ | Holm-adjusted P |
|---|---|---|---|---|---|---|
| MEDIC external | -3.24 | 0.01688 | -3.45 | 0.01944 | -4.24 | 0.02556 |
| Phenix ×1 | +2.93 | 0.0006676 | +3.22 | 0.001259 | +4.81 | 0.0061 |
| Phenix ×2 | +3.19 | 0.0006294 | +3.47 | 0.001307 | +4.97 | 0.004099 |
| Phenix after MEDIC | -1.31 | 0.5547 | -1.45 | 0.6236 | -1.65 | 0.4323 |
| CryoNet.Refine | -0.20 | 0.6742 | -0.16 | 0.8983 | -0.67 | 0.1322 |
| Servalcat | +3.15 | 9.346e-05 | +2.63 | 0.0007343 | +3.24 | 0.0061 |
| KNexPHENIX | -0.30 | 0.0577 | -0.20 | 0.2691 | -0.21 | 0.4323 |

### Supplementary Table S3 — Complementary side-chain measures

| arm | symmetry-aware $\Delta\chi_2$ (°) | Holm-adjusted P | local side-chain ΔRMSD (Å) | Holm-adjusted P | $\Delta\chi_1$-class agreement | Holm-adjusted P |
|---|---|---|---|---|---|---|
| MEDIC external | -2.77 | 0.07247 | -0.110 | 0.02193 | +0.019 | 0.2913 |
| Phenix ×1 | +4.08 | 0.0005245 | +0.050 | 0.09617 | -0.019 | 0.06694 |
| Phenix ×2 | +4.57 | 0.001045 | +0.064 | 0.09617 | -0.019 | 0.06742 |
| Phenix after MEDIC | +0.73 | 0.8983 | -0.064 | 0.1063 | +0.012 | 0.6073 |
| CryoNet.Refine | -1.75 | 0.0002861 | -0.006 | 0.7562 | +0.000 | 0.6346 |
| Servalcat | +3.70 | 2.67e-05 | +0.158 | 1.335e-05 | +0.000 | 0.925 |
| KNexPHENIX | -0.53 | 0.04594 | -0.026 | 0.09617 | +0.000 | 0.5765 |

### Supplementary Table S4 — Sequence context of the selected chains

| case | reference | target len | reference len | matches | substitutions | target-only | reference-only | identity/longer | exact | scored residues within 8 Å |
|---|---|---|---|---|---|---|---|---|---|---|
| 3JD3 | 5K12 | 496 | 294 | 293 | 1 | 202 | 0 | 0.591 | no | 75 |
| 5GJW | 6JPA | 1291 | 1274 | 1267 | 1 | 23 | 6 | 0.981 | no | 34 |
| 5YWD | 7W4O | 1154 | 1411 | 1152 | 2 | 0 | 257 | 0.816 | no | 76 |
| 6B3Q | 7K1F | 948 | 956 | 948 | 0 | 0 | 8 | 0.992 | no | 26 |
| 6B70 | 7K1F | 949 | 956 | 949 | 0 | 0 | 7 | 0.993 | no | 27 |
| 6DMR | 7T6O | 606 | 606 | 606 | 0 | 0 | 0 | 1.000 | yes | 0 |
| 6F36 | 6RD5 | 228 | 217 | 215 | 0 | 13 | 2 | 0.943 | no | 2 |
| 6HRB | 7BGY | 673 | 674 | 673 | 0 | 0 | 1 | 0.999 | no | 25 |
| 6M66 | 6WBF | 265 | 344 | 265 | 0 | 0 | 79 | 0.770 | no | 19 |
| 6M68 | 6WBF | 265 | 344 | 265 | 0 | 0 | 79 | 0.770 | no | 14 |
| 6PBE | 7T6O | 604 | 606 | 604 | 0 | 0 | 2 | 0.997 | no | 6 |
| 6PBF | 7T6O | 598 | 606 | 598 | 0 | 0 | 8 | 0.987 | no | 7 |
| 6PVM | 7MIJ | 644 | 639 | 639 | 0 | 5 | 0 | 0.992 | no | 5 |
| 6PVP | 7MIJ | 661 | 639 | 627 | 1 | 33 | 11 | 0.949 | no | 40 |
| 6RNY | 7ADU | 338 | 367 | 338 | 0 | 0 | 29 | 0.921 | no | 3 |
| 6S5T | 7MIR | 752 | 750 | 750 | 0 | 2 | 0 | 0.997 | no | 0 |
| 6T24 | 6W7V | 370 | 358 | 357 | 0 | 13 | 1 | 0.965 | no | 41 |
| 6T7S | 6TA5 | 1030 | 1030 | 1030 | 0 | 0 | 0 | 1.000 | yes | 0 |
| 6WBI | 7DWB | 329 | 421 | 316 | 1 | 12 | 104 | 0.751 | no | 20 |
| 6X17 | 7RCP | 415 | 416 | 413 | 2 | 0 | 1 | 0.993 | no | 59 |
| 6XH7 | 6XL5 | 1340 | 1340 | 1340 | 0 | 0 | 0 | 1.000 | yes | 0 |
| 6XH8 | 6XL5 | 1340 | 1340 | 1340 | 0 | 0 | 0 | 1.000 | yes | 0 |
| 7CCS | 7P9V | 469 | 464 | 464 | 0 | 5 | 0 | 0.989 | no | 0 |
| 7E5Z | 7VW6 | 775 | 913 | 775 | 0 | 0 | 138 | 0.849 | no | 101 |
| 7K3A | 7QA4 | 318 | 488 | 318 | 0 | 0 | 170 | 0.652 | no | 37 |
| 7L2J | 7LP9 | 529 | 634 | 529 | 0 | 0 | 105 | 0.834 | no | 27 |

Only coordinate-modeled residues are counted. The extracted selected chains contain no usable SEQRES records; full construct and arbitrary PTM equality are unavailable.

### Supplementary Table S5 — Reference-context filters

| filter | retained residues | cases/clusters | Phenix ×1 Δ (Holm) | Phenix ×2 Δ (Holm) | Servalcat Δ (Holm) |
|---|---|---|---|---|---|
| all | 6480 | 26/20 | +2.93 (0.000668) | +3.19 (0.000629) | +3.15 (9.35e-05) |
| component >4 Å | 6120 | 26/20 | +2.91 (0.00161) | +3.18 (0.00126) | +3.00 (0.000134) |
| component >6 Å | 5879 | 26/20 | +3.13 (0.00193) | +3.37 (0.00197) | +3.09 (9.35e-05) |
| component >8 Å | 5284 | 26/20 | +3.28 (0.00197) | +3.54 (0.00157) | +3.11 (4.01e-05) |
| local backbone ≤0.5 Å | 4869 | 26/20 | +4.24 (0.000668) | +4.55 (0.000492) | +3.79 (2.67e-05) |
| local backbone ≤1.0 Å | 6193 | 26/20 | +3.22 (0.000839) | +3.46 (0.000492) | +3.36 (2.67e-05) |
| >6 Å and local ≤1.0 Å | 5610 | 26/20 | +3.43 (0.00241) | +3.65 (0.00193) | +3.32 (1.34e-05) |
| >8 Å from modeled sequence difference | 5836 | 26/20 | +3.61 (0.00197) | +4.03 (0.000286) | +3.34 (2.67e-05) |

### Supplementary Table S6 — Descriptive reference strata

| stratum | cases | clusters | Phenix ×1 Δ | Phenix ×2 Δ | Servalcat Δ |
|---|---|---|---|---|---|
| cryo-EM reference | 21 | 16 | +2.61 | +2.80 | +2.60 |
| X-ray reference | 5 | 4 | +4.22 | +4.79 | +5.33 |
| nonwater component present | 20 | 16 | +2.63 | +2.91 | +3.40 |
| nonwater component absent | 6 | 4 | +4.11 | +4.33 | +2.14 |
| exact modeled-chain sequence | 4 | 3 | +1.13 | +1.23 | +2.30 |

Small strata are descriptive and are not interaction or equivalence tests.

### Supplementary Table S7 — Outcome completeness

| arm | missing $\chi_1$ outcomes on the prespecified residues evaluable in all included models |
|---|---|
| CryoNet.Refine | 0 |
| KNexPHENIX | 0 |
| Phenix after MEDIC | 0 |
| Phenix ×1 | 0 |
| Phenix ×2 | 0 |
| No operation | 0 |
| MEDIC external | 0 |
| Servalcat | 0 |

### Supplementary Table S8 — Full-map support-matched CC

| atom support | arm | cluster-weighted $\Delta CC_{mask}$ | clusters |
|---|---|---:|---:|
| all target-chain heavy atoms | Phenix ×1 | +0.00638 | 20 |
| all target-chain heavy atoms | Phenix ×2 | +0.01223 | 20 |
| all target-chain heavy atoms | CryoNet.Refine | +0.01141 | 20 |
| all target-chain heavy atoms | Servalcat | +0.06061 | 20 |
| backbone only | Phenix ×1 | +0.00027 | 20 |
| backbone only | Phenix ×2 | +0.00362 | 20 |
| backbone only | CryoNet.Refine | +0.01273 | 20 |
| backbone only | Servalcat | +0.03126 | 20 |
| side chains only | Phenix ×1 | +0.01347 | 20 |
| side chains only | Phenix ×2 | +0.02079 | 20 |
| side chains only | CryoNet.Refine | +0.01305 | 20 |
| side chains only | Servalcat | +0.05536 | 20 |
| prespecified $\chi_1$-residue side chains | Phenix ×1 | +0.01102 | 20 |
| prespecified $\chi_1$-residue side chains | Phenix ×2 | +0.01865 | 20 |
| prespecified $\chi_1$-residue side chains | CryoNet.Refine | +0.01410 | 20 |
| prespecified $\chi_1$-residue side chains | Servalcat | +0.05257 | 20 |

All rows use one target-chain global common-heavy-atom identity set across five arms, occupancy 1.00 and B=30 Å$^2$. Fixing B and occupancy isolates coordinate contributions to correlation and therefore does not quantify the total workflow contribution of refined atomic displacement parameters. The spatial mask remains coordinate-derived.

### Supplementary Table S9 — Per-case atom support and chain composition

| case | reference | starting chain count | target chain | global common heavy atoms | prespecified $\chi_1$ side-chain atoms |
|---|---|---|---|---|---|
| 3JD3 | 5K12 | 6 | A | 3880 | 540 |
| 5GJW | 6JPA | 5 | A | 10261 | 1963 |
| 5YWD | 7W4O | 1 | B | 8941 | 2391 |
| 6B3Q | 7K1F | 4 | A | 7743 | 2117 |
| 6B70 | 7K1F | 8 | A | 7748 | 2152 |
| 6DMR | 7T6O | 4 | A | 4853 | 650 |
| 6F36 | 6RD5 | 12 | M | 1730 | 207 |
| 6HRB | 7BGY | 4 | B | 4998 | 1236 |
| 6M66 | 6WBF | 7 | A | 2134 | 258 |
| 6M68 | 6WBF | 7 | A | 2134 | 156 |
| 6PBE | 7T6O | 4 | A | 4797 | 824 |
| 6PBF | 7T6O | 4 | A | 4588 | 694 |
| 6PVM | 7MIJ | 4 | A | 5251 | 872 |
| 6PVP | 7MIJ | 4 | A | 5356 | 947 |
| 6RNY | 7ADU | 12 | K | 2686 | 536 |
| 6S5T | 7MIR | 2 | A | 5977 | 1512 |
| 6T24 | 6W7V | 5 | A | 2888 | 764 |
| 6T7S | 6TA5 | 3 | J | 7812 | 1734 |
| 6WBI | 7DWB | 7 | E | 1870 | 162 |
| 6X17 | 7RCP | 3 | A | 3086 | 740 |
| 6XH7 | 6XL5 | 8 | C | 10569 | 2197 |
| 6XH8 | 6XL5 | 8 | C | 10569 | 2192 |
| 7CCS | 7P9V | 2 | A | 3650 | 1130 |
| 7E5Z | 7VW6 | 2 | A | 5951 | 1453 |
| 7K3A | 7QA4 | 12 | A | 2454 | 638 |
| 7L2J | 7LP9 | 4 | A | 4276 | 858 |

### Supplementary Table S10 — Bidirectional half-map results

Directions are averaged within each case before cases are averaged into nine equal-weight reference clusters.

| arm | bidirectional evaluation-half $\Delta CC_{mask}$ | 95% interval | Holm-adjusted P | bidirectional continuous $\Delta\chi_1$ (°) | 95% interval | Holm-adjusted P | A→B-minus-B→A $\chi_1$ Holm-adjusted P |
|---|---|---|---|---|---|---|---|
| Phenix ×1 | +0.01678 | [+0.00293, +0.03227] | 0.07422 | +4.023 | [+2.077, +6.132] | 0.01562 | 1.000 |
| Servalcat | +0.04558 | [+0.01654, +0.08244] | 0.02344 | +2.767 | [+1.899, +3.598] | 0.01172 | 1.000 |
| CryoNet.Refine | +0.02405 | [+0.00684, +0.04672] | 0.02344 | +0.0368 | [−0.5469, +0.5923] | 0.9102 | 1.000 |

The direction-difference p values are not equivalence tests. Each evaluation half is withheld only from the incremental run in the opposite direction, not from construction of the deposited starting model.

The prespecified B-factor sensitivity below uses the same nine equal-weight reference clusters and reruns only the fixed isotropic B value in the evaluation-half $\chi_1$-side-chain $CC_{mask}$ calculation. All point estimates remain positive from B=10 to 80 Å$^2$. Holm-adjusted threshold decisions vary for Phenix and CryoNet.Refine. B=30 Å$^2$ is the prespecified primary analysis; the other values are sensitivities.

| fixed B (Å$^2$) | arm | bidirectional evaluation-half $\Delta CC_{mask}$ | unadjusted 95% interval | Holm-adjusted P |
|---|---|---|---|---|
| 10 | Phenix ×1 | +0.01435 | [+0.00045, +0.02998] | 0.19531 |
| 10 | Servalcat | +0.04277 | [+0.01444, +0.07892] | 0.01172 |
| 10 | CryoNet.Refine | +0.02157 | [+0.00428, +0.04492] | 0.19531 |
| 20 | Phenix ×1 | +0.01554 | [+0.00167, +0.03112] | 0.07422 |
| 20 | Servalcat | +0.04418 | [+0.01548, +0.08067] | 0.02344 |
| 20 | CryoNet.Refine | +0.02280 | [+0.00554, +0.04587] | 0.03906 |
| **30 (primary)** | Phenix ×1 | **+0.01678** | **[+0.00293, +0.03227]** | **0.07422** |
| **30 (primary)** | Servalcat | **+0.04558** | **[+0.01654, +0.08244]** | **0.02344** |
| **30 (primary)** | CryoNet.Refine | **+0.02405** | **[+0.00684, +0.04672]** | **0.02344** |
| 50 | Phenix ×1 | +0.01939 | [+0.00563, +0.03464] | 0.03906 |
| 50 | Servalcat | +0.04837 | [+0.01856, +0.08626] | 0.01562 |
| 50 | CryoNet.Refine | +0.02666 | [+0.00956, +0.04852] | 0.01172 |
| 80 | Phenix ×1 | +0.02359 | [+0.01014, +0.03819] | 0.01953 |
| 80 | Servalcat | +0.05236 | [+0.02144, +0.09216] | 0.01562 |
| 80 | CryoNet.Refine | +0.03078 | [+0.01398, +0.05148] | 0.01172 |

### Supplementary Table S11 — Software and analysis scope

| component | version/configuration | analysis scope |
|---|---|---|
| Phenix local V0-matched arm | 2.2-6143; `minimization_global+local_grid_search+nqh_flips`, no ADP | one prespecified coordinate-update refinement configuration plus a second pass; not a default/optimal parameter survey |
| Servalcat | 0.4.142, 10 CG cycles | one prespecified local refinement configuration |
| CryoNet.Refine | official checkpoint, recycles 300, max_tokens 1000 | one output per case; multi-seed variability not tested |
| KNexPHENIX | Workflow 1 compatibility mapping | later version-mapped sensitivity workflow |
| MEDIC-distributed external pipeline | distributed multi-stage output | external non-matched comparator |
| $CC_{mask}$ | phenix.map_correlations | molecular support/B/occupancy fixed; spatial mask coordinate-derived |
| bidirectional half maps | 10 cases, 2 directions, 9 clusters, 880 evaluations | direction averaged within case before cluster-level inference |
| AQuaRef public structural analysis | 40-case primary; 41-case mandatory sensitivity; 23 components | developer-generated external cohort, not local execution or MEDIC arm |
| AQuaRef public same-map CC analysis | 574 tasks; 40-case primary and 41-case sensitivity; 23 components | seven-arm atom-support-matched, coordinate-derived $CC_{mask}$; same refinement map, not withheld |
| public Phenix V0 | global minimization + local grid search + N/Q/H flips; no ADP | public AQuaRef-cohort comparator; same strategy as the local Phenix arm |
| public Phenix V1 | global minimization; $C\beta$, Ramachandran and secondary-structure restraints disabled | public AQuaRef-cohort comparator |
| public Phenix V2 | global minimization; Ramachandran enabled; $C\beta$ and secondary-structure restraints disabled | public AQuaRef-cohort comparator |
| public Phenix V3 | global minimization; secondary structure enabled; $C\beta$ and Ramachandran restraints disabled | public AQuaRef-cohort comparator |
| CERES sensitivity | 19 cases / 15 clusters; nine-arm common primary support | developer-operated Phenix service workflow; 13/19 inputs contain additional components |
| practical-effect summaries | 26 cases / 20 clusters; 7 nonbaseline arms | exploratory thresholds; no new confirmatory family |

### Supplementary Table S12 — Analysis units and baselines

| analysis | statistical unit | baseline | interpretation |
|---|---|---|---|
| MEDIC primary | 20 equal-weight reference clusters | unrefined deposited model | signed incremental protocol effects |
| bidirectional half maps | 9 equal-weight reference clusters after within-case direction averaging | deposited model in each assignment | evaluation-half fit and agreement with comparison coordinates |
| AQuaRef public cohort | 23 equal-weight reference components | public regularized starting model | map fit and side-chain agreement relative to a common starting model |
| CERES service sensitivity | 15 equal-weight reference clusters | raw target-chain model on the available subset | workflow-level sensitivity with input-composition limitation |
| practical distributions | 20 equal-weight reference clusters | arm-minus-start change or signed residue threshold | exploratory magnitude and direction summaries without P values |

### Supplementary Table S13 — Two modeled-chain identity denominators

| name | formula | interpretation | observed minimum |
|---|---|---|---|
| shorter-chain identity | `matches / min(L_target, L_reference)` | identity over the shorter coordinate-modeled chain | 0.960 |
| longer-length-normalized identity | `matches / max(L_target, L_reference)` | identity penalized by unmatched modeled-chain length; plotted in Extended Data Fig. 2c | approximately 0.591 |

The denominators answer different questions: shorter-chain identity describes the overlapping modelled sequence, whereas longer-length normalization also penalizes unmatched modelled length. Neither includes unresolved construct sequence, post-translational modifications, biological interfaces or crystal contacts.

### Supplementary Table S14 — Developer-generated AQuaRef public external cohort, primary absolute-30 rule

All seven public arms use identical endpoint-specific paired residue support. Values are changes in equal-component mean continuous symmetry-aware $\chi_1$ disagreement relative to the public regularized start. Lower is better.

| public output arm | $\Delta\chi_1$ (°) | unadjusted 95% component-bootstrap interval | Holm-adjusted P |
|---|---|---|---|
| AQuaRef | **−0.9307** | [−1.3560, −0.5079] | **0.00204** |
| Phenix V0 | +2.7159 | [+0.7120, +4.7164] | 0.02744 |
| Phenix V1 | +1.2590 | [+0.4263, +2.0630] | 0.01736 |
| Phenix V2 | +0.2431 | [−0.3330, +0.8599] | 0.53992 |
| Phenix V3 | +0.8394 | [−0.3323, +1.8677] | 0.03911 |
| Servalcat | **+3.1653** | [+2.0123, +4.2190] | **0.000362** |

Primary eligibility retains 40 cases and 23 reference components. The public cohort was selected and generated by the AQuaRef developers; we re-scored their published outputs.

### Supplementary Table S15 — Length-adaptive AQuaRef sensitivity analysis

| public output arm | $\Delta\chi_1$ (°) | unadjusted 95% component-bootstrap interval | Holm-adjusted P |
|---|---|---|---|
| AQuaRef | **−0.9274** | [−1.3532, −0.5025] | **0.00204** |
| Phenix V0 | +2.6804 | [+0.6571, +4.6865] | 0.03033 |
| Phenix V1 | +1.2149 | [+0.3950, +2.0103] | 0.01736 |
| Phenix V2 | +0.2154 | [−0.3501, +0.8311] | 0.58034 |
| Phenix V3 | +0.8010 | [−0.3589, +1.8274] | 0.04274 |
| Servalcat | **+3.1499** | [+2.0057, +4.1983] | **0.000362** |

The sensitivity retains 41 cases and 23 reference components. It was prespecified alongside the primary analysis.

### Supplementary Table S16 — AQuaRef eligibility flow

| stage | absolute-30 primary | length-adaptive sensitivity |
|---|---|---|
| acquired cases | 41 | 41 |
| acquired official mappings | 64 | 64 |
| mappings passing rule | 63 | 64 |

| stage | absolute-30 primary | length-adaptive sensitivity |
|---|---|---|
| retained cases | 40 | 41 |
| analysis mappings | 60 | 64 |
| reference components | 23 | 23 |

The requirement of at least 30 aligned residues excludes 8G94 because official mapping G→3HUP chain A has 27 aligned residues. The adaptive rule requires at least 20 aligned residues, identity ≥0.80 and aligned coverage of the shorter chain ≥0.80.

### Supplementary Table S17 — Sensitivity analysis using CERES-refined models

All nine displayed arms are recomputed on identical primary support for 19 cases and 15 clusters.

| arm | cluster-weighted $\chi_1$ mean (°) | Δ versus raw (°) | unadjusted 95% interval | Holm-adjusted P |
|---|---|---|---|---|
| developer-operated public CERES service workflow | 37.91 | **+3.81** | [+2.24, +5.57] | **0.00427** |
| CryoNet.Refine | 33.68 | −0.41 | [−1.01, +0.06] | 0.416 |
| KNexPHENIX v1 | 33.64 | −0.46 | [−0.77, −0.14] | 0.0452 |
| Phenix from relaxed | 32.37 | −1.72 | [−3.90, +0.52] | 0.416 |
| Phenix ×1 | 37.32 | +3.22 | [+1.72, +5.08] | 0.00299 |
| Phenix ×2 | 37.49 | +3.40 | [+1.85, +5.27] | 0.00244 |
| no operation | 34.09 | — | — | — |
| Rosetta-relaxed | 30.38 | −3.71 | [−5.71, −1.69] | 0.0215 |
| Servalcat | 36.66 | +2.57 | [+1.55, +3.54] | 0.00299 |

For CERES itself, the unadjusted two-sided P value is 0.000854 and the full-precision change is +3.8148°. Six cases have exact full input composition; 13 service inputs include additional components from the full PDB entry. CERES uses a Phenix service workflow. Map fit was not evaluated for CERES.

### Supplementary Table S18 — Exploratory fractions of side chains with reduced or increased $\chi_1$ disagreement

All values are equal-reference-cluster mean percentages with unadjusted 95% cluster-bootstrap intervals. Disagreement decrease means arm-minus-start $\chi_1$ disagreement below the negative threshold; disagreement increase means above the positive threshold. The endpoint is change in agreement with the comparison structure.

| arm | disagreement decrease >10° | disagreement increase >10° | decrease >20° | increase >20° | decrease >40° | increase >40° |
|---|---|---|---|---|---|---|
| Phenix ×1 | 8.80 [7.04, 10.65] | 13.77 [11.50, 16.09] | 4.22 [3.08, 5.43] | 7.95 [6.41, 9.55] | 2.34 [1.64, 3.15] | 4.31 [3.11, 5.71] |
| Phenix ×2 | 10.90 [9.47, 12.49] | 17.14 [14.82, 19.49] | 4.97 [3.71, 6.28] | 8.47 [6.85, 10.11] | 2.78 [1.91, 3.75] | 4.67 [3.28, 6.23] |
| CryoNet.Refine | 8.14 [4.99, 11.63] | 7.26 [5.36, 9.25] | 1.33 [0.65, 2.09] | 0.84 [0.40, 1.32] | 0.00 [0.00, 0.00] | 0.00 [0.00, 0.00] |
| Servalcat | 14.25 [12.12, 16.39] | 27.83 [26.11, 29.58] | 4.78 [3.79, 5.77] | 11.19 [9.41, 12.73] | 0.94 [0.59, 1.37] | 1.49 [1.05, 1.95] |
| MEDIC external / Rosetta-relaxed | 17.93 [13.58, 22.19] | 10.16 [7.54, 12.96] | 8.99 [6.57, 11.45] | 5.82 [4.07, 7.70] | 6.20 [4.38, 8.10] | 4.53 [3.19, 5.93] |
| KNexPHENIX v1 | 3.37 [2.25, 4.61] | 2.12 [1.26, 3.02] | 0.78 [0.34, 1.30] | 0.55 [0.20, 0.97] | 0.11 [0.02, 0.22] | 0.10 [0.01, 0.21] |
| Phenix after MEDIC | 19.18 [15.87, 22.46] | 17.02 [15.04, 19.21] | 10.22 [8.15, 12.32] | 9.43 [7.84, 11.12] | 6.58 [4.75, 8.50] | 5.59 [4.42, 6.79] |

Fractions are calculated within case, averaged within shared-reference clusters and then averaged across 20 equal-weight clusters. These prespecified exploratory thresholds have no P values.

### Supplementary Table S19 — Exploratory $\chi_1$-class transitions and local side-chain RMSD fractions

Values are equal-reference-cluster mean percentages with unadjusted 95% cluster-bootstrap intervals. RMSD Δ is arm absolute fraction minus the raw-start absolute fraction on one eight-arm global RMSD support.

| arm | $\chi_1$-class agreement lost | $\chi_1$-class agreement gained | RMSD >0.5 Å absolute | RMSD >0.5 Å Δ | RMSD >1.0 Å absolute | RMSD >1.0 Å Δ |
|---|---|---|---|---|---|---|
| Phenix ×1 | 4.13 [2.89, 5.57] | 2.28 [1.60, 3.05] | 76.59 [71.68, 81.17] | 2.02 [-0.38, 4.77] | 53.45 [46.25, 60.91] | 3.49 [1.03, 5.96] |
| Phenix ×2 | 4.64 [3.23, 6.21] | 2.73 [1.87, 3.71] | 78.26 [73.70, 82.43] | 3.68 [0.46, 7.72] | 54.23 [47.49, 61.19] | 4.27 [1.57, 7.04] |
| CryoNet.Refine | 0.02 [0.00, 0.06] | 0.03 [0.00, 0.09] | 74.87 [69.46, 80.04] | 0.29 [-1.23, 2.15] | 48.76 [41.75, 55.92] | -1.20 [-2.58, -0.05] |
| Servalcat | 0.53 [0.35, 0.75] | 0.58 [0.32, 0.88] | 89.31 [85.82, 92.36] | 14.74 [11.00, 19.20] | 59.09 [52.13, 66.23] | 9.14 [5.91, 12.54] |
| MEDIC external / Rosetta-relaxed | 4.22 [2.96, 5.55] | 6.15 [4.36, 8.02] | 65.70 [59.44, 72.12] | -8.88 [-12.77, -4.84] | 41.97 [35.49, 49.01] | -7.99 [-11.45, -4.12] |
| KNexPHENIX v1 | 0.05 [0.01, 0.09] | 0.09 [0.03, 0.17] | 71.74 [65.63, 77.51] | -2.84 [-5.00, -0.57] | 48.47 [41.20, 55.74] | -1.49 [-2.95, -0.22] |
| Phenix after MEDIC | 5.25 [4.05, 6.46] | 6.47 [4.70, 8.35] | 68.87 [63.57, 74.03] | -5.71 [-10.07, -0.61] | 45.47 [39.62, 51.70] | -4.48 [-8.23, -0.59] |

The prespecified common RMSD support contains at least 28 and at most 482 residues per case. Individual residues are not treated as independent replicates.

### Supplementary Table S20 — AQuaRef public cohort: map fit and side-chain agreement

All values are arm-minus-regularized-start equal-reference-component means on the same 40 cases and 23 reference components. Same-map $CC_{mask}$ uses the seven-arm common heavy side-chain atoms from reportable $\chi_1$ residues, occupancy 1 and B=30 Å$^2$; the spatial mask remains coordinate-derived. Negative $\chi_1$ disagreement denotes increased agreement with the matched higher-resolution reference. Each endpoint uses Holm adjustment across six nonbaseline protocols.

| arm | Δ same-map $CC_{mask}$ | 95% interval | Holm-adjusted P | Δ $\chi_1$ disagreement (°) | 95% interval | Holm-adjusted P |
|---|---|---|---|---|---|---|
| AQuaRef | **−0.008684** | [−0.012705, −0.004397] | **0.001279** | **−0.931** | [−1.356, −0.508] | **0.002040** |
| Phenix V0 | +0.029555 | [+0.018655, +0.045805] | 0.00005397 | +2.716 | [+0.712, +4.716] | 0.02744 |
| Phenix V1 | +0.036377 | [+0.027521, +0.046579] | 0.000001431 | +1.259 | [+0.426, +2.063] | 0.01736 |
| Phenix V2 | +0.025649 | [+0.019200, +0.032774] | 0.000001431 | +0.243 | [−0.333, +0.860] | 0.5399 |
| Phenix V3 | +0.033136 | [+0.025198, +0.042090] | 0.000001431 | +0.839 | [−0.332, +1.868] | 0.03911 |
| Servalcat | **+0.038435** | [+0.027757, +0.051158] | **0.000001431** | **+3.165** | [+2.012, +4.219] | **0.0003619** |

The AQuaRef developers selected this public cohort and generated its refined coordinates. The timing of the map-fit analysis is described in Supplementary Note 2. The mean-aligned exact sign-flip sensitivity in Supplementary Table S22 preserves the AQuaRef and Servalcat conclusions but does not preserve the Wilcoxon-based Phenix V3 threshold decision.

### Supplementary Table S21 — AQuaRef map-correlation and mapping sensitivity analyses

Each entry is an equal-reference-component arm-minus-regularized-start $\Delta CC_{mask}$ with its unadjusted 95% component-bootstrap interval and a Holm-adjusted P value within that rule × support surface. The mapped-chain common-heavy-atom set is a secondary diagnostic; the scored side-chain set remains primary.

| rule / support | arm | $\Delta CC_{mask}$ [95% interval] | Holm-adjusted P |
|---|---|---|---|
| primary 40-case / mapped-chain global common heavy | AQuaRef | −0.010316 [−0.014649, −0.005836] | 0.0001814 |
| primary 40-case / mapped-chain global common heavy | Phenix V0 | +0.028898 [+0.019826, +0.039885] | 0.000001431 |
| primary 40-case / mapped-chain global common heavy | Phenix V1 | +0.049844 [+0.038242, +0.063509] | 0.000001431 |
| primary 40-case / mapped-chain global common heavy | Phenix V2 | +0.031123 [+0.023296, +0.040057] | 0.000001431 |
| primary 40-case / mapped-chain global common heavy | Phenix V3 | +0.044594 [+0.033389, +0.057433] | 0.000001431 |
| primary 40-case / mapped-chain global common heavy | Servalcat | +0.056796 [+0.044062, +0.072723] | 0.000001431 |
| adaptive 41-case / reportable $\chi_1$ side-chain | AQuaRef | −0.008716 [−0.012744, −0.004425] | 0.001279 |
| adaptive 41-case / reportable $\chi_1$ side-chain | Phenix V0 | +0.029382 [+0.018436, +0.045703] | 0.00005397 |
| adaptive 41-case / reportable $\chi_1$ side-chain | Phenix V1 | +0.036086 [+0.027147, +0.046300] | 0.000001431 |
| adaptive 41-case / reportable $\chi_1$ side-chain | Phenix V2 | +0.025411 [+0.018917, +0.032607] | 0.000001431 |
| adaptive 41-case / reportable $\chi_1$ side-chain | Phenix V3 | +0.032878 [+0.024904, +0.041837] | 0.000001431 |
| adaptive 41-case / reportable $\chi_1$ side-chain | Servalcat | +0.037915 [+0.027096, +0.050638] | 0.000001431 |
| adaptive 41-case / mapped-chain global common heavy | AQuaRef | −0.010274 [−0.014581, −0.005798] | 0.0001814 |
| adaptive 41-case / mapped-chain global common heavy | Phenix V0 | +0.028654 [+0.019570, +0.039571] | 0.000001431 |
| adaptive 41-case / mapped-chain global common heavy | Phenix V1 | +0.049429 [+0.037795, +0.063095] | 0.000001431 |
| adaptive 41-case / mapped-chain global common heavy | Phenix V2 | +0.030832 [+0.022996, +0.039834] | 0.000001431 |
| adaptive 41-case / mapped-chain global common heavy | Phenix V3 | +0.044228 [+0.032985, +0.057160] | 0.000001431 |
| adaptive 41-case / mapped-chain global common heavy | Servalcat | +0.056462 [+0.043645, +0.072373] | 0.000001431 |

The targeted primary and adaptive $\chi_1$ directions are displayed jointly in Supplementary Fig. S6. The 41-case adaptive structural $\chi_1$ sensitivity remains AQuaRef −0.927° [−1.353, −0.502] (Holm-adjusted P=0.002040) and Servalcat +3.150° [+2.006, +4.198] (Holm-adjusted P=0.0003619); it is a prespecified sensitivity analysis of the 40-case primary result.

### Supplementary Table S22 — Mean-aligned exact sign-flip sensitivity

The reported estimand is a cluster-weighted or component-weighted arithmetic mean, whereas the prespecified Wilcoxon test is rank based. As a post hoc sensitivity aligned to the mean estimand, all `2^n` one-sample sign assignments were exhaustively enumerated using the absolute arithmetic mean as the two-sided statistic; numerical ties were included and Holm adjustment remained confined to each existing family. Exactness is conditional on sign exchangeability, equivalently symmetry of complete unit-level differences about zero. The prespecified Wilcoxon tests remain the primary analysis.

| family | arm | arithmetic-mean change | exact sign-flip Holm P |
|---|---|---:|---:|
| MEDIC $\chi_1$, 20 clusters | Phenix ×1 | +2.929° | 0.0006008 |
| MEDIC $\chi_1$, 20 clusters | Phenix ×2 | +3.195° | 0.0005951 |
| MEDIC $\chi_1$, 20 clusters | Servalcat | +3.145° | 0.0000801 |
| MEDIC $\chi_1$, 20 clusters | CryoNet.Refine | −0.200° | 0.4582 |
| MEDIC $\chi_1$, 20 clusters | MEDIC external / Rosetta-relaxed | −3.240° | 0.01551 |
| MEDIC $\chi_1$, 20 clusters | KNexPHENIX | −0.303° | 0.07086 |
| MEDIC $\chi_1$, 20 clusters | Phenix after MEDIC | −1.308° | 0.4330 |
| full-map $CC_{mask}$, 20 clusters | Phenix ×1 | +0.01102 | 0.001026 |
| full-map $CC_{mask}$, 20 clusters | Phenix ×2 | +0.01865 | 0.0000744 |
| full-map $CC_{mask}$, 20 clusters | CryoNet.Refine | +0.01410 | 0.001026 |
| full-map $CC_{mask}$, 20 clusters | Servalcat | +0.05257 | 0.00000763 |
| bidirectional evaluation-half $CC_{mask}$, 9 clusters | Phenix ×1 | +0.01678 | 0.06250 |
| bidirectional evaluation-half $CC_{mask}$, 9 clusters | Servalcat | +0.04558 | 0.02344 |
| bidirectional evaluation-half $CC_{mask}$, 9 clusters | CryoNet.Refine | +0.02405 | 0.02344 |
| bidirectional $\chi_1$, 9 clusters | Phenix ×1 | +4.023° | 0.01563 |
| bidirectional $\chi_1$, 9 clusters | Servalcat | +2.767° | 0.01172 |
| bidirectional $\chi_1$, 9 clusters | CryoNet.Refine | +0.0368° | 0.9063 |
| AQuaRef structural primary, 23 components | AQuaRef | −0.9307° | 0.001558 |
| AQuaRef structural primary, 23 components | Phenix V0 | +2.7159° | 0.04427 |
| AQuaRef structural primary, 23 components | Phenix V1 | +1.2590° | 0.02914 |
| AQuaRef structural primary, 23 components | Phenix V2 | +0.2431° | 0.4774 |
| AQuaRef structural primary, 23 components | Phenix V3 | +0.8394° | 0.3320 |
| AQuaRef structural primary, 23 components | Servalcat | +3.1653° | 0.0004163 |
| AQuaRef targeted same-map CC, 23 components | AQuaRef | −0.008684 | 0.0008883 |
| AQuaRef targeted same-map CC, 23 components | Servalcat | +0.038435 | 0.000001431 |
| CERES structural, 15 clusters | CERES service workflow | +3.8148° | 0.002563 |
| CERES structural, 15 clusters | KNexPHENIX | −0.4554° | 0.05017 |

All central MEDIC, support-matched CC, bidirectional half-map, AQuaRef and Servalcat conclusions preserve their prespecified Wilcoxon decisions. Two peripheral threshold decisions are test-sensitive: public Phenix V3 in the AQuaRef structural family (Wilcoxon Holm P=0.0391; sign-flip Holm P=0.3320) and KNexPHENIX in the CERES family (0.0452 versus 0.0502). The complete 12-family result file, including the adaptive and global-support surfaces, will be included in the code and data archive indexed through the Primum repository (main-text Code availability).

### Supplementary Table S23 — Descriptive MEDIC resolution stratification

This post hoc exploratory sensitivity uses the symmetry-aware case means. Positive $\Delta\chi_1$ denotes reduced agreement with the separately determined comparison structure and negative $\Delta\chi_1$ denotes greater agreement. Only descriptive point estimates are reported.

| arm | ≤4.5 Å cluster-weighted $\Delta\chi_1$ | clusters + / − | >4.5 Å cluster-weighted $\Delta\chi_1$ | clusters + / − |
|---|---:|---:|---:|---:|
| MEDIC external / Rosetta-relaxed | −3.68° | 5 / 14 | −0.84° | 1 / 1 |
| Phenix ×1 | +2.65° | 14 / 5 | +5.03° | 2 / 0 |
| Phenix ×2 | +3.07° | 16 / 3 | +3.48° | 2 / 0 |
| Phenix after MEDIC | −1.64° | 7 / 12 | −1.94° | 1 / 1 |
| CryoNet.Refine | −0.17° | 9 / 10 | −0.67° | 0 / 2 |
| Servalcat | +3.08° | 17 / 2 | +1.64° | 1 / 1 |
| KNexPHENIX | −0.31° | 4 / 15 | −0.54° | 1 / 1 |

The ≤4.5 Å stratum contains 24 cases and 19 reference clusters; the >4.5 Å stratum contains only two cases and two clusters. Reference cluster 6WBF spans the split (6M66 at 4.1 Å and 6M68 at 4.6 Å), so the strata are not independent. The smaller stratum is insufficient for resolution-effect modification or equivalence inference.

### Supplementary Table S24 — Nine-cluster BCa interval sensitivity

For the nine-cluster bidirectional analyses, all 24,310 nine-part bootstrap count compositions were exactly enumerated and weighted by their multinomial multiplicities, representing all `9^9` ordered resamples without Monte Carlo error. Bias correction used a midrank treatment of ties and acceleration used delete-one-reference-cluster jackknife estimates. These post hoc BCa intervals supplement the prespecified percentile intervals.

| endpoint | arm | mean | original percentile 95% interval | exact-weighted BCa 95% interval |
|---|---|---:|---|---|
| evaluation-half $CC_{mask}$ | Phenix ×1 | +0.01678 | [+0.00293, +0.03227] | [+0.00423, +0.03441] |
| evaluation-half $CC_{mask}$ | Servalcat | +0.04558 | [+0.01654, +0.08244] | [+0.02021, +0.09365] |
| evaluation-half $CC_{mask}$ | CryoNet.Refine | +0.02405 | [+0.00684, +0.04672] | [+0.00920, +0.05417] |
| bidirectional $\chi_1$ disagreement | Phenix ×1 | +4.023° | [+2.077, +6.132] | [+2.170, +6.318] |
| bidirectional $\chi_1$ disagreement | Servalcat | +2.767° | [+1.899, +3.598] | [+1.881, +3.586] |
| bidirectional $\chi_1$ disagreement | CryoNet.Refine | +0.0368° | [−0.5469, +0.5923] | [−0.5666, +0.5561] |
| A→B minus B→A $\chi_1$ | Phenix ×1 | +0.0108° | [−0.3105, +0.3873] | [−0.2752, +0.4501] |
| A→B minus B→A $\chi_1$ | Servalcat | −0.0189° | [−0.2374, +0.2199] | [−0.2227, +0.2519] |
| A→B minus B→A $\chi_1$ | CryoNet.Refine | −0.0286° | [−0.1306, +0.0556] | [−0.1493, +0.0454] |

The BCa and percentile intervals have the same side-of-zero pattern for all nine rows. Because `n=9`, the BCa results remain small-sample sensitivities rather than additional multiplicity-adjusted tests. In particular, the unadjusted Phenix evaluation-half CC interval excludes zero while its Holm-adjusted exact sign-flip P is 0.0625; this P value exceeds the 0.05 threshold.

### Supplementary Table S25 — Relaxed buried-residue threshold sensitivity

The primary ≥18-neighbour analysis is unchanged. This post hoc exploratory sensitivity uses ≥12 starting-model C$\alpha$ neighbours within 10 Å, symmetry-aware $\chi_1$, arm-complete common support and the same 26 cases/20 reference clusters. It contains 12,108 residue keys; the excluded 6DJP case was not reconsidered. Positive $\Delta\chi_1$ denotes reduced agreement with the separately determined comparison structure. Confidence intervals are unadjusted and P values are Holm-adjusted within this sensitivity family.

| arm | cluster-weighted $\Delta\chi_1$ | unadjusted 95% interval | Holm-adjusted P |
|---|---|---|---|
| MEDIC external / Rosetta-relaxed | −3.01° | [−4.90, −1.09] | 0.05447 |
| Phenix ×1 | +3.14° | [+1.81, +4.71] | 0.0003147 |
| Phenix ×2 | +3.42° | [+2.09, +4.93] | 0.00005722 |
| Phenix after MEDIC | −0.94° | [−3.00, +1.19] | 1.000 |
| CryoNet.Refine | −0.22° | [−0.67, +0.17] | 1.000 |
| Servalcat | +3.20° | [+2.55, +3.86] | 0.00001335 |
| KNexPHENIX | −0.20° | [−0.49, +0.10] | 1.000 |

At the primary threshold of 18 neighbours the same scorer reproduces the primary analysis exactly. The three locally matched positive directions are robust to the relaxed threshold. The MEDIC external arm remains negative but no longer passes the Holm-adjusted threshold, so its adjusted evidence is threshold-sensitive.

### Supplementary Table S26 — Full MEDIC multiplicity family and provenance roles

The primary Holm family contains seven nonbaseline arms, but only the four locally executed matched protocols are shown in Supplementary Table S33. The three remaining rows are contextual comparisons with different input states or provenance. Intervals are unadjusted percentile cluster-bootstrap intervals; P values are two-sided Wilcoxon signed-rank values adjusted by Holm across the seven nonbaseline arms.

| arm | provenance role | cluster-weighted absolute mean (°) | cluster-weighted $\Delta\chi_1$ (°) | unadjusted 95% interval | Holm-adjusted P | missing |
|---|---|---|---|---|---|---|
| no operation | deposited model | 34.12 | — | — | — | 0 |
| Phenix ×1 | locally executed primary protocol | 37.05 | +2.93 | [+1.65, +4.46] | $6.68\times10^{-4}$ | 0 |
| Phenix ×2 | locally executed repeat pass | 37.31 | +3.19 | [+1.80, +4.75] | $6.29\times10^{-4}$ | 0 |
| Servalcat | locally executed primary protocol | 37.27 | +3.15 | [+2.26, +4.02] | $9.35\times10^{-5}$ | 0 |
| CryoNet.Refine | locally executed primary protocol | 33.92 | −0.20 | [−0.68, +0.20] | 0.674 | 0 |
| MEDIC-distributed external pipeline | external non-matched multi-stage output | 30.88 | −3.24 | [−5.09, −1.35] | 0.0169 | 0 |
| KNexPHENIX Workflow 1 | later version-mapped sensitivity workflow | 33.82 | −0.30 | [−0.54, −0.07] | 0.0577 | 0 |
| Phenix after MEDIC pipeline | composed workflow from another input state | 32.81 | −1.31 | [−3.27, +0.64] | 0.555 | 0 |

### Supplementary Table S27 — Independent replication cohort

Entries were selected from archive metadata alone (Methods). The search returned 2,758 low-resolution single-particle entries, 2,569 met the entry-level rules, 1,237 had an eligible later, higher-resolution comparison structure in the same 95% sequence-identity group, and 200 were accepted in identifier-derived order (at most two per group; 14 skipped for map size). 156 entries met the eligibility rules (≥95% aligned identity, ≥20 packed $\chi_1$ residues). Resolutions in Å; release = initial release date; group = RCSB 95% sequence-identity group of the target entity; half maps = two half maps listed in EMDB.

The full 200-entry table (rank, entry, EMDB map, resolution, release date, comparison structure, method, resolution and release date, 95% sequence group, half-map availability and status) is provided as Supplementary Data 1.

### Supplementary Table S28 — Refinement configurations in the sweep

All runs start from the deposited model. Options not listed keep the values of the original protocols (Phenix 2.2-6143: minimization_global+local_grid_search+nqh_flips with automatic weight; Servalcat 0.4.142: full map, ten cycles, automatic weight with adjustment).

| Refinement configuration | Program | Change from the original protocol | Cases |
|---|---|---|---|
| + ADP | Phenix | ADP refinement added (`refinement.run=...+adp`) | 26 |
| + reference restraints | Phenix | reference-model restraints to the starting model | 26 |
| tight geometry | Phenix | target bond and angle r.m.s.d. 0.005 Å and 0.5° | 26 |
| simulated annealing | Phenix | morphing and simulated annealing added; excluded because one run exceeded the 4-h limit | 26 |
| jelly-body | Servalcat | `--jellybody` (default $\sigma$ 0.01, $d_{max}$ 4.2 Å) | 26 |
| 0.5× weight | Servalcat | `--weight` 0.5× the case’s automatic weight, `--no_weight_adjust` | 26 |
| 2× weight | Servalcat | `--weight` 2× the case’s automatic weight, `--no_weight_adjust` | 26 |
| fixed B | Servalcat | `--adp fix` | 26 |
| half maps | Servalcat | `--halfmaps` with the two deposited half maps instead of `--map` | 10 |

### Supplementary Table S29 — Configuration sweep results

MEDIC, 26 cases and 20 clusters; cluster-weighted means with 95% percentile cluster-bootstrap intervals (10,000 draws, seed 20261014). Changes are relative to no operation; positive $\Delta\chi_1$ is worse agreement. The simulated-annealing configuration was excluded after one run exceeded the 4-h limit.

**a, Per refinement configuration.**

| Program | Refinement configuration | $\Delta CC_{mask}$ | $\Delta\chi_1$ (°) | Clusters worse |
|---|---|---|---|---|
| Phenix | ×1 (original) | +0.0110 [+0.0059, +0.0165] | +2.93 [+1.61, +4.42] | 16/20 |
| Phenix | ×2 (original) | +0.0186 [+0.0112, +0.0284] | +3.19 [+1.77, +4.75] | 17/20 |
| Phenix | + ADP | +0.0123 [+0.0074, +0.0175] | +2.99 [+1.70, +4.44] | 17/20 |
| Phenix | + reference restraints | +0.0131 [+0.0072, +0.0207] | +2.13 [+1.12, +3.16] | 15/20 |
| Phenix | tight geometry | +0.0091 [+0.0033, +0.0162] | +2.84 [+1.55, +4.26] | 16/20 |
| Servalcat | default (original) | +0.0526 [+0.0372, +0.0771] | +3.15 [+2.28, +4.00] | 18/20 |
| Servalcat | 0.5× weight | +0.0397 [+0.0269, +0.0598] | +1.55 [+0.81, +2.29] | 17/20 |
| Servalcat | 2× weight | +0.0683 [+0.0508, +0.0965] | +5.15 [+4.19, +6.15] | 20/20 |
| Servalcat | fixed B | +0.0512 [+0.0372, +0.0721] | +2.88 [+2.10, +3.68] | 18/20 |
| Servalcat | jelly-body | +0.0076 [+0.0058, +0.0098] | -0.03 [-0.15, +0.09] | 9/20 |
| CryoNet.Refine | original | +0.0141 [+0.0061, +0.0252] | -0.20 [-0.70, +0.18] | 9/20 |

**b, Highest-$CC_{mask}$ selection over action sets** (action counts include no operation; regret in degrees).

| Action set | Actions | Selection loss (°) | Worse | Regret | Selected actions |
|---|---|---|---|---|---|
| Original actions | 5 | +2.83 [+1.89, +3.74] | 17/20 | 3.39 | Servalcat default 25, Phenix ×2 1 |
| All actions | 12 | +5.05 [+4.10, +6.06] | 20/20 | 5.73 | Servalcat 2× 24, Servalcat default 1, Servalcat fixed B 1 |
| Without Servalcat | 7 | +2.32 [+1.14, +3.58] | 15/20 | 2.93 | Phenix ×2 11, Phenix + reference 7, CryoNet.Refine 4, Phenix + ADP 2, Phenix ×1 1, Phenix tight 1 |
| Without Phenix | 7 | +5.05 [+4.10, +6.06] | 20/20 | 5.58 | Servalcat 2× 24, Servalcat default 1, Servalcat fixed B 1 |
| Servalcat only | 6 | +5.05 [+4.10, +6.06] | 20/20 | 5.34 | Servalcat 2× 24, Servalcat default 1, Servalcat fixed B 1 |
| Phenix only | 6 | +2.59 [+1.43, +3.81] | 16/20 | 2.88 | Phenix ×2 13, Phenix + reference 8, Phenix + ADP 2, Phenix ×1 1, Phenix tight 1, no operation 1 |

**c, Servalcat half-map input** (10 cases, 9 clusters with deposited half maps): $\Delta\chi_1$ +3.19 [+2.33, +3.97]° with half maps and +3.72 [+2.61, +4.83]° with the full map on the same cases; difference -0.53° [-1.09, -0.13].

### Supplementary Table S30 — Residue-level effects of highest-$CC_{mask}$ selection

MEDIC, 6,480 buried side chains in 26 cases and 20 clusters. For the action selected by each rule, relative to no operation: side chains whose symmetry-aware $\chi_1$ disagreement with the comparison structure increased (moved away) or decreased (moved closer) by more than 20°, and side chains whose canonical $\chi_1$ class lost or gained agreement with the comparison structure. Counts are pooled over residues; percentages are cluster-weighted fractions with 95% cluster-bootstrap intervals (10,000 draws, seed 20261016). Values are descriptive.

| Action set | Moved away >20° | Moved closer >20° | $\chi_1$ class lost | $\chi_1$ class gained |
|---|---|---|---|---|
| Original actions (5) | 763 (10.7% [8.9, 12.2]) | 358 (4.9% [3.9, 6.0]) | 33 (0.5% [0.3, 0.7]) | 44 (0.7% [0.3, 1.3]) |
| All actions (12) | 1,132 (17.1% [15.1, 19.0]) | 450 (6.7% [5.5, 8.0]) | 66 (1.0% [0.7, 1.3]) | 57 (0.8% [0.4, 1.2]) |
| Without Servalcat (7) | 395 (6.9% [5.1, 8.8]) | 290 (4.4% [3.1, 5.7]) | 206 (3.6% [2.2, 5.0]) | 153 (2.2% [1.3, 3.2]) |

### Supplementary Table S31 — Selection by a half map withheld from re-refinement

Replication entries with two deposited half maps (70 entries; 70 scored). Each refinement configuration was run against each half map and scored against the other (evaluation half) and its own (work half). Scores and $\chi_1$ disagreement are averaged over both directions for each configuration before selecting one configuration per entry. The reported outcome is that configuration's averaged disagreement; selected-action counts sum to 69 entries. Selection-loss rows use the entries in which every action is valid. Cluster-weighted means with 95% cluster-bootstrap intervals (10,000 draws, seed 20261015); positive $\Delta\chi_1$ is worse agreement.

**a, Per action** (95% sequence groups).

| Action | $\Delta CC_{mask}$ work half | $\Delta CC_{mask}$ evaluation half | $\Delta\chi_1$ (°) | Worse |
|---|---|---|---|---|
| Phenix ×1 | +0.0174 [+0.0123, +0.0228] | +0.0038 [-0.0014, +0.0095] | +3.26 [+2.44, +4.14] | 59/63 |
| Phenix ×2 | +0.0211 [+0.0152, +0.0274] | +0.0055 [-0.0008, +0.0123] | +4.01 [+3.03, +5.05] | 59/63 |
| Servalcat | +0.0344 [+0.0282, +0.0409] | +0.0124 [+0.0070, +0.0183] | +2.07 [+1.55, +2.61] | 53/64 |

**b, Highest-$CC_{mask}$ selection** over no operation, Phenix ×1, Phenix ×2 and Servalcat.

| Scored on | Clustering unit | Selection loss (°) | Worse | Selected actions |
|---|---|---|---|---|
| evaluation half | sequence group | +1.83 [+1.27, +2.44] | 43/63 | Servalcat default 43, no operation 15, Phenix ×2 11 |
| work half | sequence group | +2.44 [+1.85, +3.07] | 53/63 | Servalcat default 56, Phenix ×2 11, no operation 2 |
| evaluation half | comparison structure | +1.82 [+1.26, +2.41] | 43/64 | Servalcat default 43, no operation 15, Phenix ×2 11 |
| work half | comparison structure | +2.51 [+1.89, +3.16] | 54/64 | Servalcat default 56, Phenix ×2 11, no operation 2 |

### Supplementary Table S32 — Prespecified replication results

Replication cohort: 156 eligible entries, 156 scored, 130 sequence groups. Every action was refined with the prespecified MEDIC protocols and scored exactly as in MEDIC; the 95% sequence-identity group is the primary clustering unit and comparison structures the sensitivity unit. Configurations excluded under the prespecified 10% failure rule: none. Cluster-weighted means with 95% percentile cluster-bootstrap intervals (10,000 draws, seed 20261015); changes are relative to no operation and positive $\Delta\chi_1$ is worse agreement. Phenix + ADP produced coordinates identical to Phenix ×1 in every entry (ADP refinement changes only B factors, which are fixed for scoring), so its values coincide with Phenix ×1. †Entries with two deposited half maps only.

**a, Per refinement configuration.**

| Program | Refinement configuration | $\Delta CC_{mask}$ | $\Delta\chi_1$ (°) | Groups worse | $\Delta\chi_1$, comparison clusters |
|---|---|---|---|---|---|
| Phenix | ×1 (original) | +0.0180 [+0.0136, +0.0228] | +2.84 [+2.20, +3.53] | 109/130 | +2.83 [+2.21, +3.49] |
| Phenix | ×2 (original) | +0.0217 [+0.0168, +0.0270] | +3.56 [+2.82, +4.36] | 107/130 | +3.54 [+2.81, +4.30] |
| Phenix | + ADP | +0.0180 [+0.0136, +0.0228] | +2.84 [+2.20, +3.53] | 109/130 | +2.83 [+2.21, +3.49] |
| Phenix | + reference restraints | +0.0148 [+0.0109, +0.0191] | +2.69 [+2.04, +3.43] | 104/130 | +2.70 [+2.06, +3.40] |
| Phenix | tight geometry | +0.0134 [+0.0092, +0.0180] | +2.77 [+2.10, +3.52] | 109/130 | +2.75 [+2.11, +3.46] |
| Phenix | simulated annealing | +0.0186 [+0.0131, +0.0244] | +4.22 [+3.41, +5.09] | 115/130 | +4.23 [+3.42, +5.07] |
| Servalcat | default (original) | +0.0472 [+0.0409, +0.0537] | +3.02 [+2.65, +3.40] | 120/130 | +3.00 [+2.62, +3.38] |
| Servalcat | 0.5× weight | +0.0364 [+0.0306, +0.0424] | +1.48 [+1.16, +1.81] | 99/130 | +1.47 [+1.15, +1.79] |
| Servalcat | 2× weight | +0.0625 [+0.0556, +0.0697] | +4.93 [+4.52, +5.35] | 127/130 | +4.90 [+4.47, +5.31] |
| Servalcat | fixed B | +0.0467 [+0.0409, +0.0529] | +2.56 [+2.20, +2.93] | 115/130 | +2.54 [+2.17, +2.91] |
| Servalcat | jelly-body | +0.0116 [+0.0082, +0.0155] | +0.03 [-0.02, +0.09] | 71/130 | +0.04 [-0.01, +0.10] |
| Servalcat | half maps† | +0.0287 [+0.0199, +0.0379] | +2.37 [+1.80, +2.93] | 53/64 | +2.34 [+1.75, +2.90] |
| CryoNet.Refine | original | +0.0205 [+0.0150, +0.0263] | -0.39 [-0.62, -0.17] | 49/130 | -0.37 [-0.59, -0.16] |

**b, Highest-$CC_{mask}$ selection over the prespecified action sets** (action counts include no operation; regret in degrees).

| Rule | Clustering unit | Actions | Selection loss (°) | Worse | Regret | Selected actions |
|---|---|---|---|---|---|---|
| P1 original actions | sequence group | 5 | +3.16 [+2.68, +3.68] | 116/130 | 4.09 | Servalcat default 135, Phenix ×2 7, CryoNet.Refine 6, Phenix ×1 4, no operation 4 |
| P2 all actions | sequence group | 13 | +4.82 [+4.30, +5.35] | 124/130 | 5.95 | Servalcat 2× 134, Servalcat fixed B 10, CryoNet.Refine 3, Phenix ×2 3, Phenix annealing 2, no operation 2, Phenix ×1 1, Servalcat default 1 |
| P3 without Servalcat | sequence group | 8 | +2.14 [+1.44, +2.91] | 82/130 | 3.20 | CryoNet.Refine 47, Phenix ×2 35, Phenix annealing 22, Phenix ×1 21, no operation 16, Phenix + reference 9, Phenix tight 6 |
| P1 original actions | comparison structure | 5 | +3.13 [+2.65, +3.65] | 116/133 | 4.03 | Servalcat default 135, Phenix ×2 7, CryoNet.Refine 6, Phenix ×1 4, no operation 4 |
| P2 all actions | comparison structure | 13 | +4.77 [+4.25, +5.31] | 124/133 | 5.86 | Servalcat 2× 134, Servalcat fixed B 10, CryoNet.Refine 3, Phenix ×2 3, Phenix annealing 2, no operation 2, Phenix ×1 1, Servalcat default 1 |
| P3 without Servalcat | comparison structure | 8 | +2.18 [+1.50, +2.94] | 83/133 | 3.20 | CryoNet.Refine 47, Phenix ×2 35, Phenix annealing 22, Phenix ×1 21, no operation 16, Phenix + reference 9, Phenix tight 6 |

### Supplementary Figure S1 — Numerically selected examples across the four MEDIC protocols

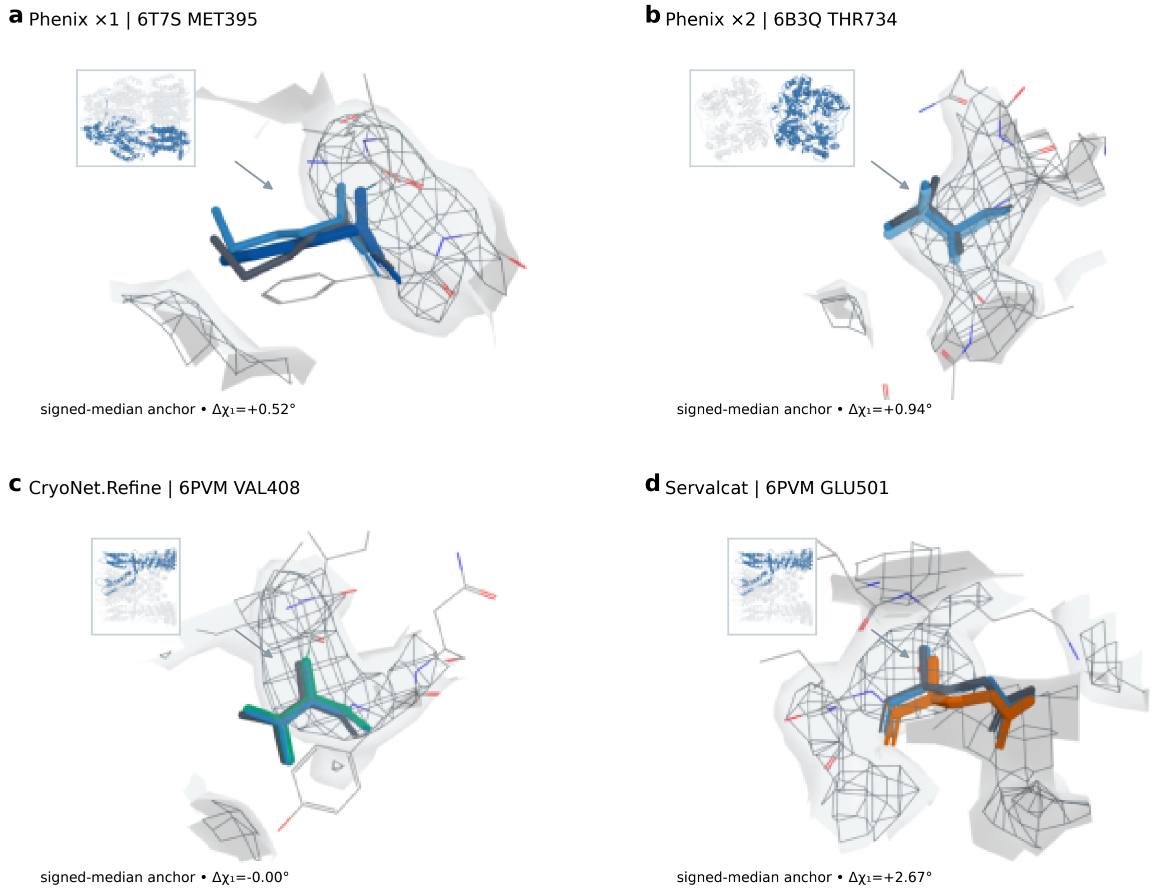

Four residues closest to each protocol's median signed $\Delta\chi_1$ were selected by numeric rank. Panels a–d show Phenix ×1, Phenix ×2, CryoNet.Refine and Servalcat, respectively. Each panel contains a whole-model locator, starting residue in blue, selected arm in its fixed method color, five-residue-Kabsch-aligned comparison residue in charcoal, and the original deposited full map as a pale surface plus higher-contour mesh. The signed $\Delta\chi_1$ value is printed on each panel. Coordinates were not realigned for display. These examples, which also appear in Extended Data Fig. 4, are descriptive.

### Supplementary Figure S2 — Matched MEDIC local atlas

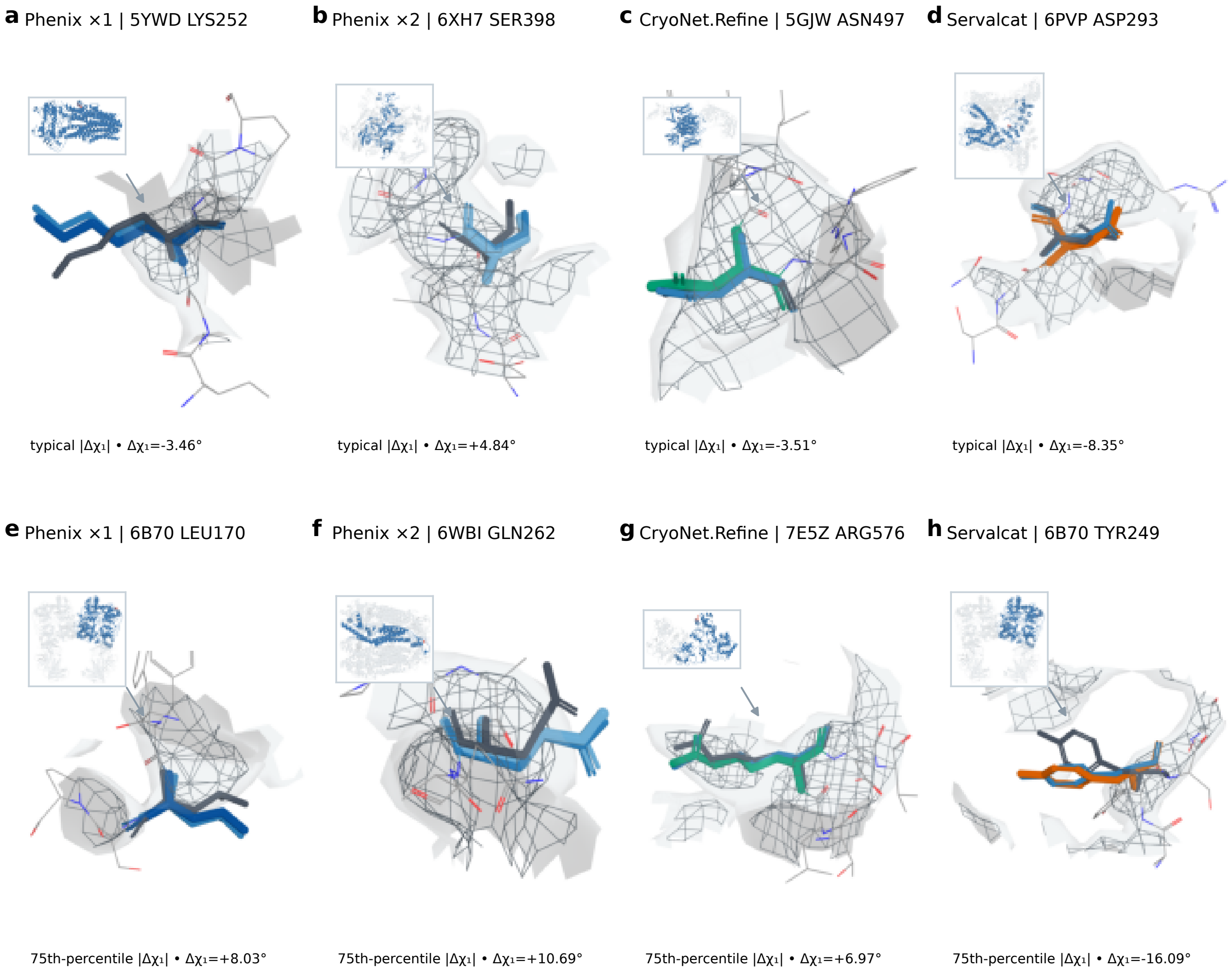

Eight additional, non-overlapping MEDIC examples cover all four matched methods. Panels a–d are closest to each arm's median absolute $\Delta\chi_1$; panels e–h are closest to its 75th-percentile absolute change and exclude the maximum extreme by construction. Rendering and color conventions match Supplementary Figure S1. Selection rankings, superposition transforms and contour levels will be included in the public data archive. The examples illustrate local heterogeneity.

### Supplementary Figure S3 — Bidirectional half-map local atlas

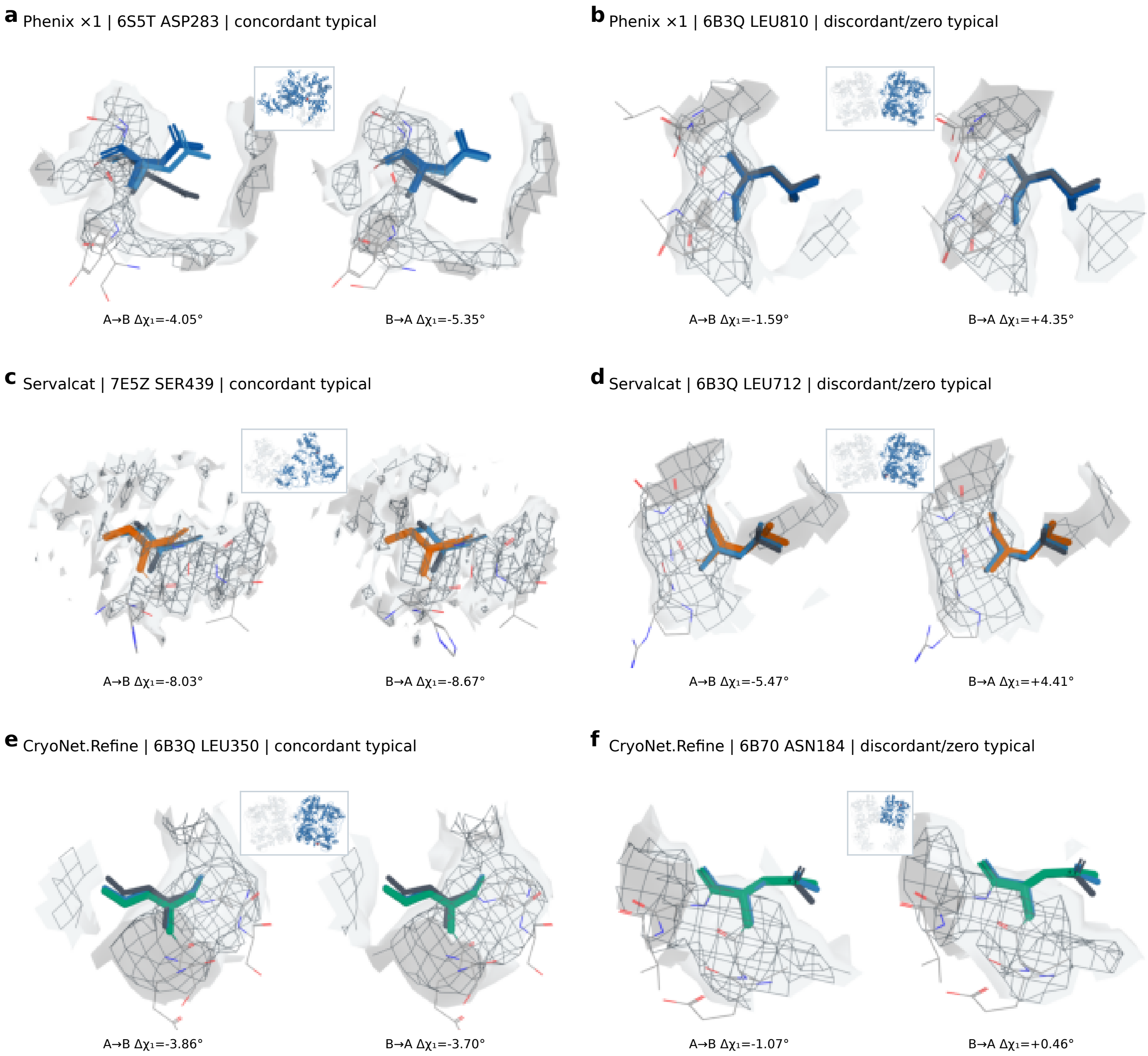

Six numerically selected examples show A⟶B and B⟶A local views with a shared camera view. Rows correspond to Phenix ×1, Servalcat and CryoNet.Refine; left panels are directionally concordant typical examples and right panels are discordant-or-zero typical examples. Labels below each image report the signed $\Delta\chi_1$ for A⟶B and B⟶A. Each pair uses the opposite evaluation half map at fixed absolute contour levels and one shared whole-model locator. Examples were selected by numeric rank and exclude residues shown in Supplementary Figs. S1 and S2. Both directions are shown for descriptive comparison; statistical analyses average directions within each case.

### Supplementary Figure S4 — Practical effects of the MEDIC protocols

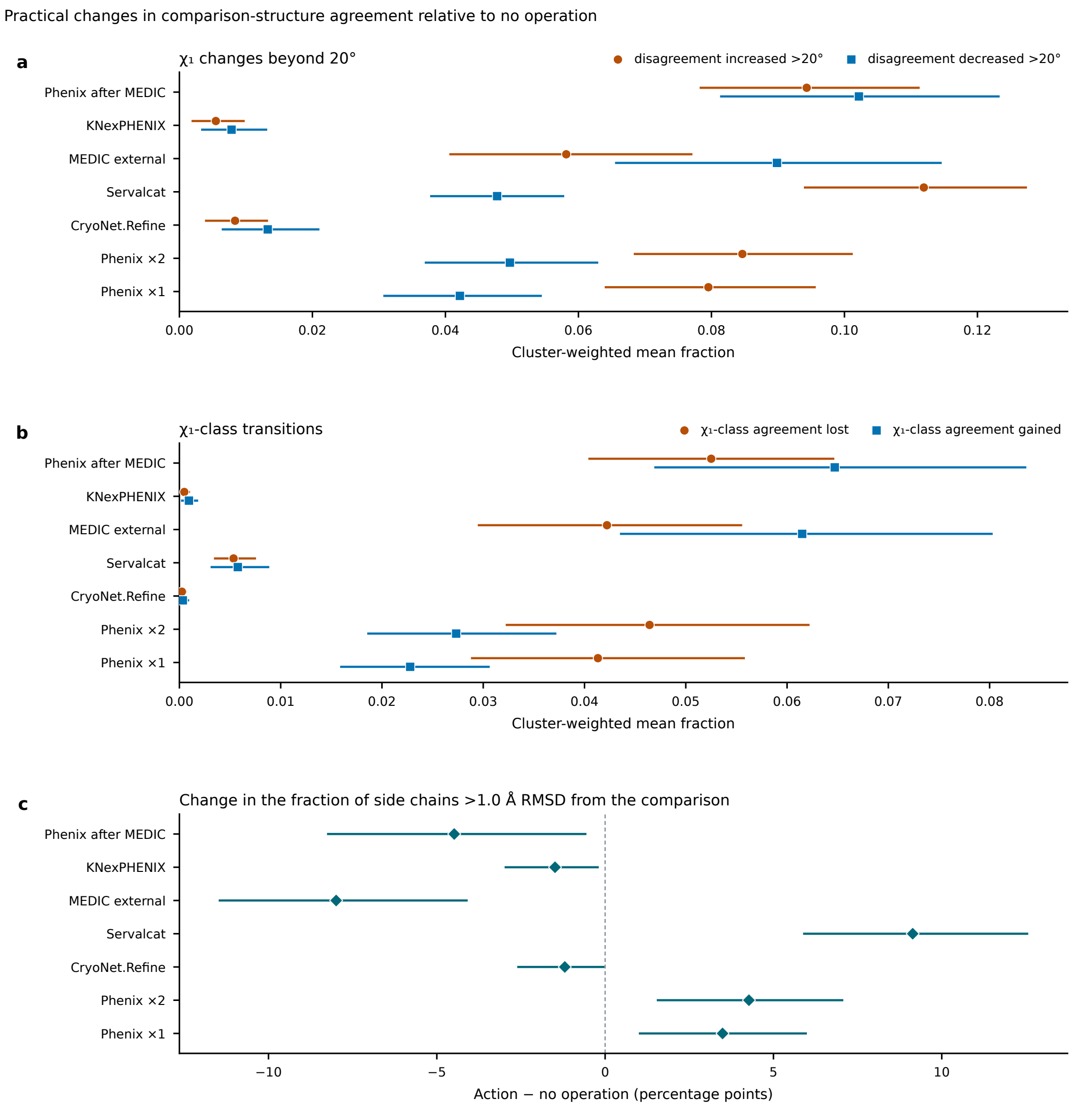

Fractions of packed side chains whose $\chi_1$ disagreement with the comparison structure increased or decreased by more than 20°, and of residues with local side-chain RMSD above 1.0 Å, for all seven MEDIC arms. Fractions are averaged within case and cluster, with unadjusted 95% cluster-bootstrap intervals. Exploratory summaries without P values (Supplementary Tables S18 and S19).

### Supplementary Figure S5 — CERES service-workflow sensitivity

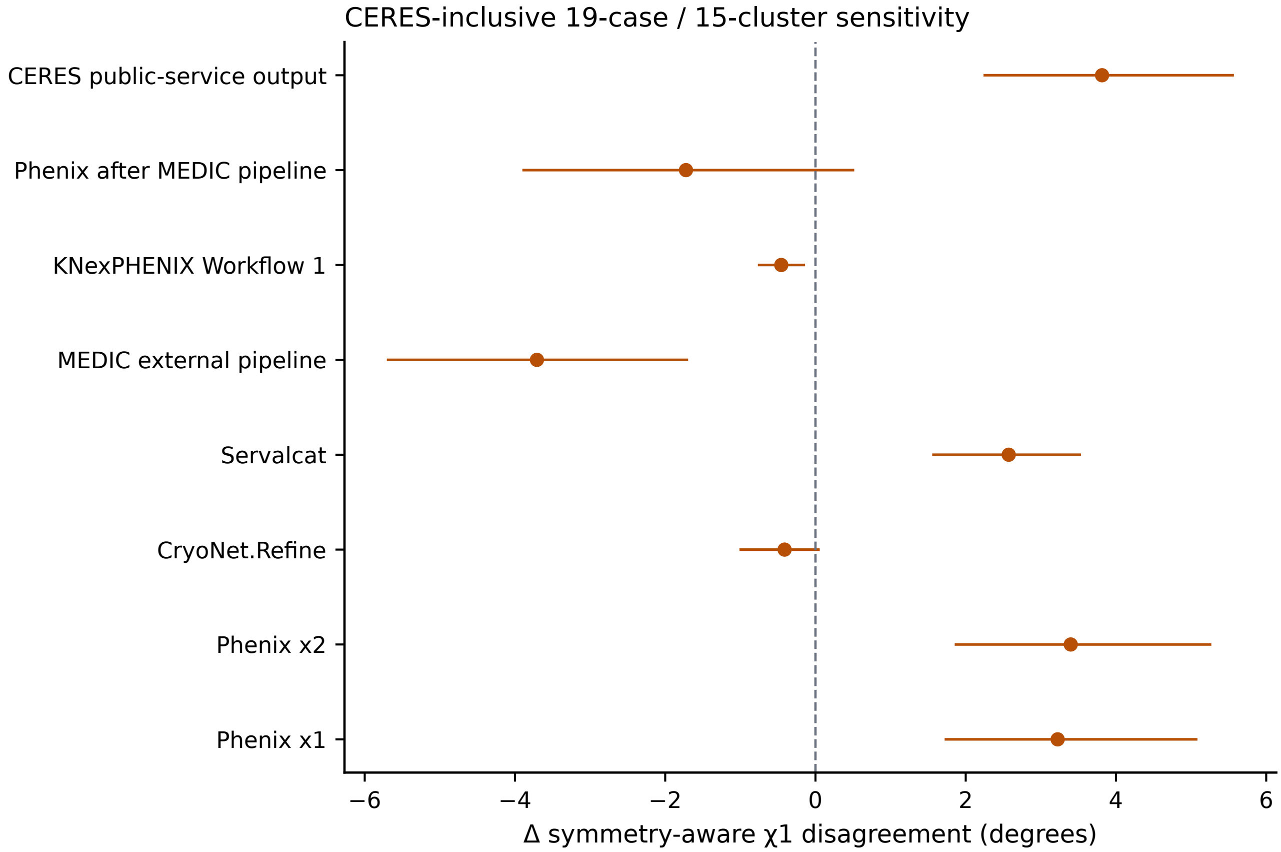

Nine arms on the common support of 19 cases and 15 reference clusters with public CERES service outputs. CERES is a developer-operated service workflow; this family is analysed separately and not pooled with MEDIC, AQuaRef or the replication cohort (Supplementary Table S17).

### Supplementary Figure S6 — AQuaRef mapping sensitivity

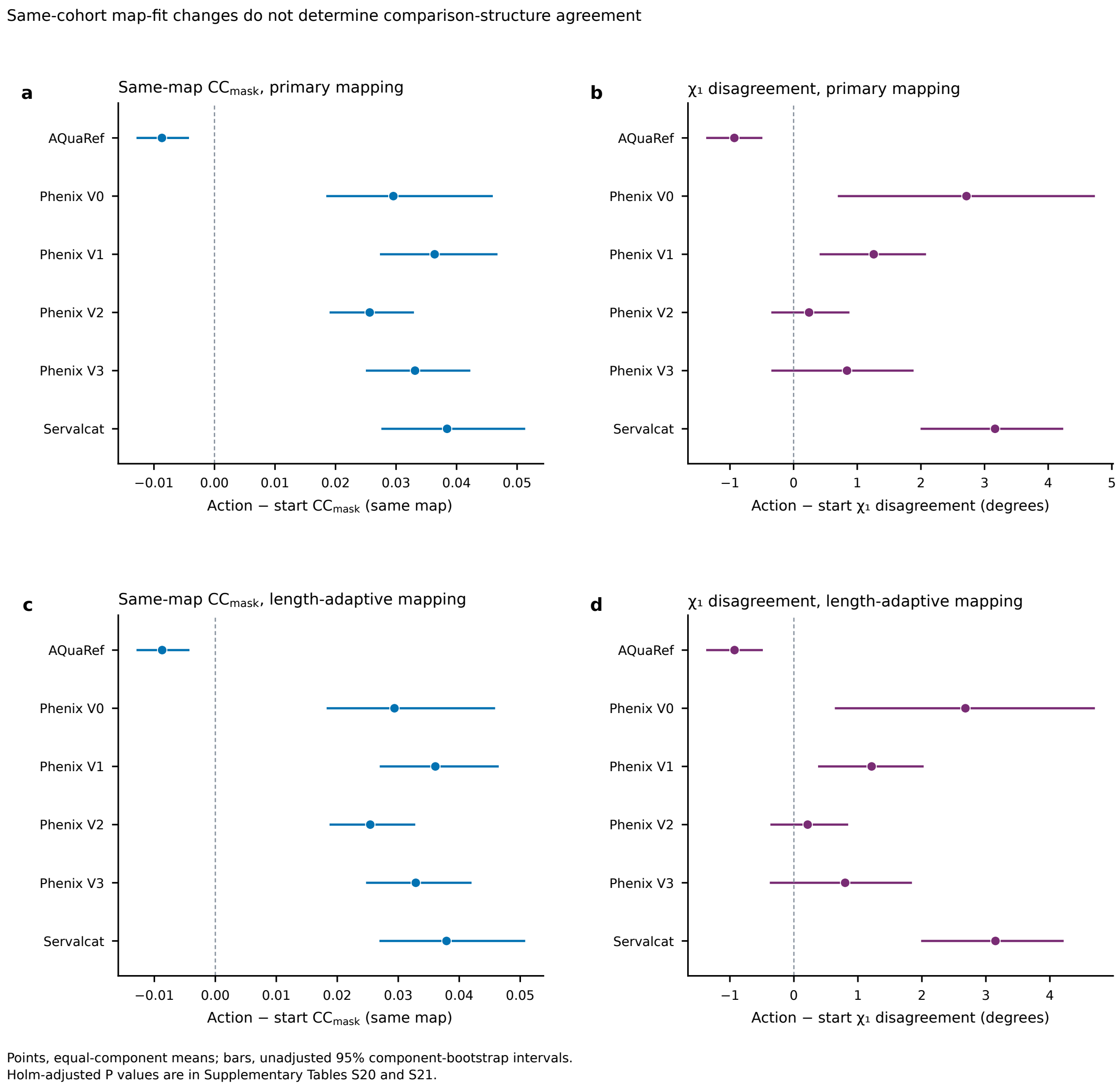

Primary 40-case and length-adaptive 41-case chain-mapping rules for same-map $CC_{mask}$ and $\chi_1$ disagreement in the developer-generated AQuaRef cohort. Bars are unadjusted 95% component-bootstrap intervals; Holm-adjusted results are in Supplementary Tables S20 and S21.

### Supplementary Note 1 — Comparison structures and residue support

Comparison structures are used to assess coordinate agreement; acquisition relationships for the within-study pairs are listed in Supplementary Table S43. Component-distance, five-residue backbone and modelled-sequence filters address measurable context differences. Crystal contacts, biological interfaces, unresolved constructs or post-translational modifications, ensemble differences, prior templates and shared rotamer-library priors remain potential sources of disagreement. Within-structure experimental-support quartiles stratify MEDIC comparison residues by their local experimental support. The primary ≥18-neighbour criterion restricts the endpoint to packed residues.

### Supplementary Note 2 — AQuaRef and CERES analyses

The developers' published AQuaRef coordinates were re-scored for a public cohort they selected and screened for comparison structures (Supplementary Note 5). The joint analysis compares same-map, coordinate-derived $CC_{mask}$ with comparison-structure $\chi_1$ agreement for the same cases and reference components. The CC definitions and tests were specified before CC computation, after the $\chi_1$ results were known. CERES results come from a developer-operated Phenix service on a 19-case subset of MEDIC; 13 service inputs include additional components from the full PDB entry. AQuaRef and CERES are analysed separately from the other cohorts.

### Supplementary Note 3 — Coordinate-derived spatial support

All full-map and half-map fit analyses use standard coordinate-derived $CC_{mask}$. Molecular identity, target chain, residue and atom support, occupancy and B factor are fixed within each comparison, but the evaluated voxel region is generated separately from each model's coordinates. EMRinger was evaluated in the MEDIC decision analysis and in acceptance-rule analyses of the 3.0–3.6 Å and within-study cohorts (Supplementary Tables S38 and S44).

### Supplementary Note 4 — Scope of the residue-level summaries

The $\chi_1$, $\chi_1$-class and local-RMSD threshold summaries describe residue-level changes relative to the comparison structures. Thresholds were specified for exploratory analysis. No P values were computed.

### Supplementary Note 5 — AQuaRef outputs

The AQuaRef analysis re-scores the developers' published coordinates for their public cohort; AQuaRef was not run locally. The published pipeline could not be run in our environment, so only the developers' published outputs are scored.

### Supplementary Note 6 — Analyses not reported

Two analyses were excluded after failing their prespecified validation criteria. A three-weight PyRosetta density sensitivity completed all 81 refinements, but one output (6DJP) exceeded the prespecified relative clashscore tolerance, so the sweep as a whole was not analysed. Coordinate-independent fixed-voxel scoring did not pass its prespecified validation against the native implementation, so no fixed-spatial result is reported.

### Supplementary Note 7 — Primum benchmark specification and access

Primum v1.0 includes the 26-case MEDIC cohort, the 156-entry replication cohort and its 70-entry half-map subset. Scoring and selection definitions are in Methods, and reference results are in Extended Data Table 1 and Supplementary Table S34. The half-map subset's common four-action selection sample contains 69 entries in 63 sequence groups. The 13-action full-map replication set contains no operation and 12 refinement configurations. Its 1,942 refined models comprise 1,872 full-map configurations and 70 Servalcat models refined with both half maps as input. The additional directional half-map analysis contributes 416 models (69 entries for each of two Phenix configurations in two directions, and 70 entries for Servalcat in two directions), giving 2,358 refined models in the replication resource.

Primum v2 adds a separate 200-entry candidate list at $3.0 \leq d < 3.6$ Å, with 168 eligible and 167 analysed entries; its four-action half-map selection sample contains 99 entries in 82 groups. The earlier cohort versions retain their original reference results. Reference strategies for this cohort are in Extended Data Table 2.

Primum v2 also includes the 22-pair within-study pilot and the further 53-pair cohort with prespecified coordinate and model-building screening. The within-study cohort contains 53 pairs with five full-map actions and 44 pairs with five half-map actions; its reference results, exclusions and pilot context are in Supplementary Tables S41–S44. Screening and acquisition relationships are part of the cohort definition.

For a new refiner, the benchmark record consists of the input coordinates and map, the candidate coordinates, case and sequence-group identifiers, residue correspondence and retained scoring support. Failed cases and missing required atoms are recorded. For a new selection rule, the case-level action table supplies the selection scores and coordinate endpoints. Candidate sets and sample exclusions are recorded with every result; the complete risk–coverage curve describes thresholds. The five reporting items in the Discussion provide a common result schema.

The project and resource-access entry point is the Primum repository listed in the main-text Code availability statement. The repository will index the cohort documentation, coordinates, residue supports, scoring software and reference-result tables. New method results can be recorded against a specified cohort, support and scorer version. Cohort versions retain their query and exclusion rules: the archive cohorts exclude previously evaluated sequence groups, and the within-study cohort excludes the pilot studies and entries. Earlier versions retain their input identifiers and reference results.

### Supplementary Table S33 — Original MEDIC protocol effects

**Locally executed MEDIC protocols, 26 cases and 20 reference clusters.**
Intervals are unadjusted percentile cluster-bootstrap estimation intervals. Two-sided Wilcoxon P values are Holm-adjusted across all seven nonbaseline MEDIC arms.

| Arm | Mean (°) | Δ (°) | Unadjusted 95% interval | Holm-adjusted P | Missing |
|---|---|---|---|---|---|
| No operation (deposited model) | 34.12 | — | — | — | 0 |
| Phenix ×1 | 37.05 | +2.93 | [+1.65, +4.46] | $6.68 \times 10^{-4}$ | 0 |
| Phenix ×2 | 37.31 | +3.19 | [+1.80, +4.75] | $6.29 \times 10^{-4}$ | 0 |
| Servalcat | 37.27 | +3.15 | [+2.26, +4.02] | $9.35 \times 10^{-5}$ | 0 |
| CryoNet.Refine | 33.92 | -0.20 | [-0.68, +0.20] | 0.674 | 0 |

### Supplementary Table S34 — Earlier Primum reference strategies

Loss is the change in $\chi_1$ disagreement relative to no operation; lower values indicate better agreement. The no-operation endpoint in the earlier replication cohort is 35.44°. Each block fixes its candidate set and evaluation sample.

| Strategy | Loss [95% interval] (°) | Groups worse | Coverage (%) | Regret (°) | Selection input |
|---|---|---|---|---|---|
| **a, Replication cohort: original five actions, 156 entries and 130 sequence groups** | | | | | |
| No operation | 0.00 [0.00, 0.00] | 0/130 | 0.0 | 0.94 | None |
| Highest full-map $CC_{mask}$ | +3.16 [+2.68, +3.68] | 116/130 | 97.4 | 4.09 | Full map |
| Always CryoNet.Refine | -0.39 [-0.62, -0.17] | 49/130 | 100.0 | 0.55 | None |
| Per-case oracle | -0.94 [-1.14, -0.75] | 0/130 | 70.5 | 0.00 | Outcome |
| **b, Replication cohort: expanded 13 actions, the same 156 entries and 130 groups** | | | | | |
| No operation | 0.00 [0.00, 0.00] | 0/130 | 0.0 | 1.12 | None |
| Highest full-map $CC_{mask}$ | +4.82 [+4.30, +5.35] | 124/130 | 98.7 | 5.95 | Full map |
| Always Servalcat jelly-body | +0.03 [-0.02, +0.09] | 71/130 | 100.0 | 1.16 | None |
| Always Servalcat 0.5× | +1.48 [+1.16, +1.81] | 99/130 | 100.0 | 2.61 | None |
| Per-case oracle | -1.12 [-1.37, -0.90] | 0/130 | 81.4 | 0.00 | Outcome |
| **c, Half-map subset: four actions, common 69 entries and 63 sequence groups** | | | | | |
| Highest work-half CC | +2.44 [+1.85, +3.07] | 53/63 | 97.1 | — | Work half |
| Highest evaluation-half CC | +1.83 [+1.27, +2.44] | 43/63 | 78.3 | — | Evaluation half |

Cluster-weighted means and 95% percentile bootstrap intervals use 10,000 draws and seed 20261015. Coverage is the pooled fraction of entries assigned a refinement; "Groups worse" counts sequence groups with positive mean loss. Regret is measured against the per-case minimum within the action set of that block. The oracle uses the observed comparison-structure outcome to give a hindsight lower bound. Fixed-program rows apply one configuration to every entry. Selection-input requirements apply after candidate models have been generated; refinement itself may require maps. An em dash denotes an unreported value. The half-map block is evaluated on its own common subset.

The original five actions are no operation, Phenix ×1/×2, CryoNet.Refine and Servalcat. The expanded set adds the eight configurations in Supplementary Table S28. MEDIC results for EMRinger, geometry and combined-signal rules are reported separately in main Fig. 3 because they use different cohorts and candidate availability.

### Supplementary Table S35 — 3.0–3.6 Å cohort membership

The 200-entry roster was selected from archive metadata before refinement. There were 168 eligible entries; 8EAP was excluded at scoring because the model did not overlap the map. The analysed sample contains 167 entries in 131 sequence groups. Resolution is in Å; "preparation" denotes an entry outside the eligible set. Detailed inclusion and file records are retained in the cohort documentation. The full 200-entry roster (rank, entry, map, resolution, comparison structure and resolution, 95% sequence group and analysis membership) is provided as Supplementary Data 2.

### Supplementary Table S36 — Half-map protocol effects at 3.0–3.6 Å

For each configuration, both directions are averaged before selection or aggregation within sequence groups (Methods). Phenix effects use 99 entries and 82 groups; Servalcat uses 102 entries and 85 groups. Changes are relative to no operation; intervals are 95% group-bootstrap intervals (10,000 draws, seed 20261101).

| Action | $\Delta\chi_1$ (°) | Work-half $\Delta CC_{mask}$ | Evaluation-half $\Delta CC_{mask}$ |
|---|---|---|---|
| Phenix ×1 | +3.25 [+2.56, +3.97] | +0.0260 | +0.0142 |
| Phenix ×2 | +3.85 [+3.10, +4.64] | +0.0290 | +0.0156 |
| Servalcat | +2.18 [+1.75, +2.64] | +0.0325 | +0.0146 |

### Supplementary Figure S7 — Original MEDIC map-fit analysis

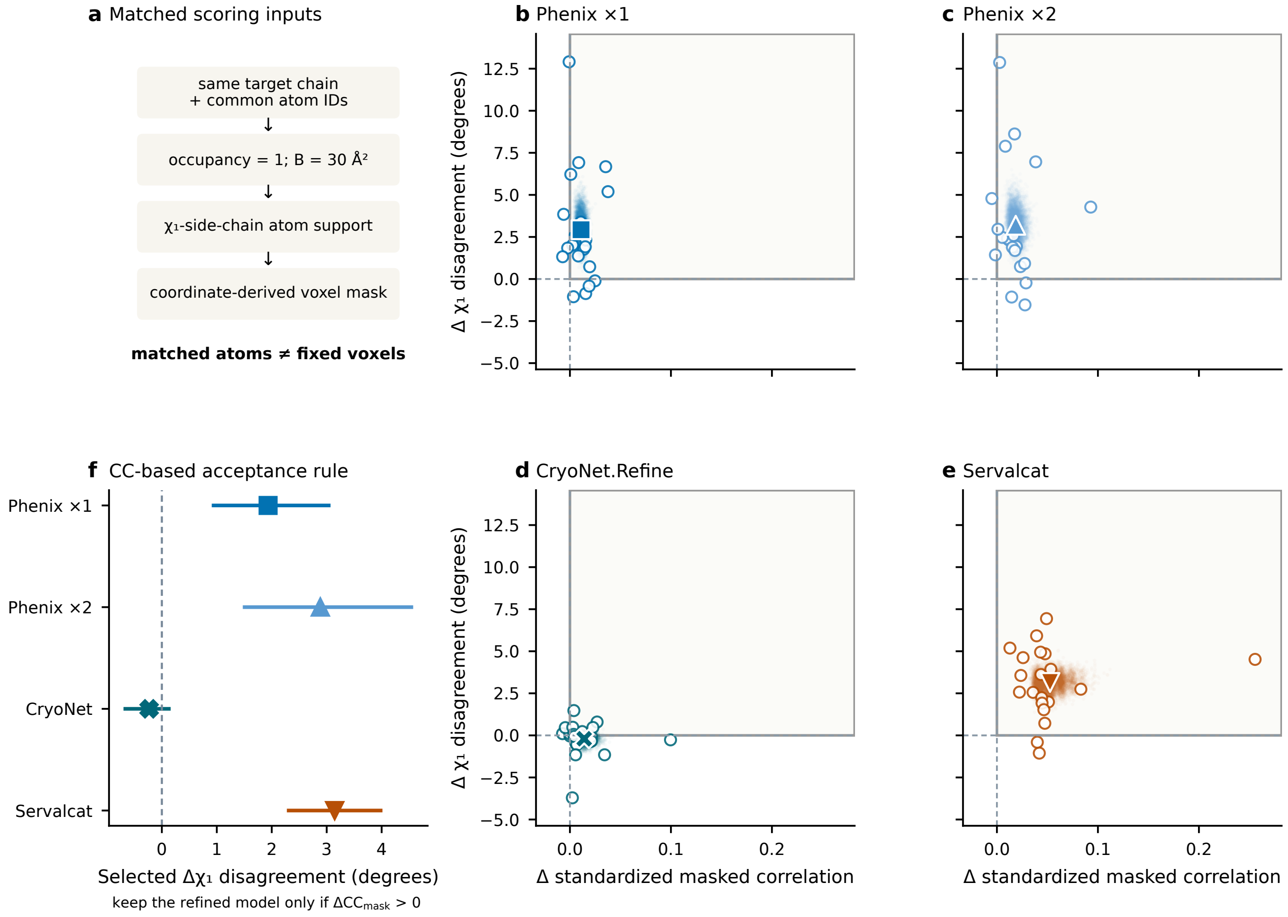

**Supplementary Figure S7 | Same-map fit and side-chain agreement on matched atoms.** **a**, Atom identity, occupancy, B factor and side-chain support are matched across actions; the voxel mask remains coordinate-derived. **b–e**, All 20 cluster values for Phenix ×1, Phenix ×2, CryoNet.Refine and Servalcat; large symbols are cluster-weighted means and pale clouds are 10,000 paired bootstrap means (bootstrap distribution of the mean). **f**, Descriptive cluster-level acceptance rule: for each refiner, average gains and outcomes within clusters; retain the cluster's $\chi_1$ change if mean $CC_{mask}$ gain is positive, otherwise zero; then average clusters equally.

### Supplementary Note 8 — Cohort protocols and scoring

The protocol, candidate list and analysis code for the 3.0–3.6 Å cohort were fixed before any model was scored; the archive snapshot used for the query is recorded in the cohort documentation. Prespecified analyses are supported by dated internal protocol documents. No external preregistration timestamp was obtained. The residue-level summaries and single-program acceptance analysis for this cohort in Supplementary Tables S37 and S39 are post hoc. Entries were ordered reproducibly from their PDB identifiers using the algorithm recorded with the analysis code. One CryoNet.Refine log (entry 7VGH) was inspected before scoring; it reports only the program's internal loss and model–map correlation.

The five full-map actions and the directional Phenix/Servalcat half-map actions used the commands of the earlier replication. 8EAP had failed Phenix alignment and Servalcat matrix checks, and both remaining candidates lacked valid baseline map correlation. No configuration exceeded the 10% failure threshold. For 6HBJ, 6TTF and 6WXH only the Phenix half-map branch failed alignment checks. Estimates for this cohort average cases within 95% sequence groups; comparison-structure clusters form a sensitivity analysis (167 entries, 135 clusters; selection loss +2.41°, 95% interval +2.12 to +2.70°). No resolution subdivision within this cohort was prespecified. This cohort is a single sample selected in a fixed identifier-derived order.

The within-study protocol and 90-study roster were recorded internally before downloading any within-study data. They specified the metadata pool, one pair per study, pilot exclusions, coordinate and model-building criteria and the primary and secondary analyses. Coordinate screening preceded refinement. At that stage, the reference coordinates were used for eligibility, alignment and baseline screening. Candidate outputs were completed and the candidate set was fixed before candidate $\chi_1$ scoring. Selection-rule thresholds remained those of the earlier calibration. Coordinate and model-building criteria were motivated by the pilot and fixed prospectively for the within-study cohort. Acquisition relationships and state equivalence were assessed from archive metadata; study identifiers, titles, major-ligand checks and reconstruction relationships are supplied in Supplementary Data 4. Shared external starting models are listed in Supplementary Table S43.

### Supplementary Note 9 — Original MEDIC selection and residue-level results

The initial cohort comprised 26 deposited cryo-EM models at 3.6–6.2 Å from the MEDIC validation set[30], each paired with a separately determined comparison structure of the same protein at higher resolution; cases that share a comparison structure form 20 equal-weight clusters (main Fig. 1a). Every case offers the deposited model as a no-operation action alongside models re-refined from it. The endpoint is chemical-symmetry-aware $\chi_1$ disagreement with the comparison structure over packed side chains (at least 18 C$\alpha$ neighbours within 10 Å; 6,480 residues). Outcomes were available for every included action on this common residue set.

Relative to no operation, Phenix ×1, Phenix ×2 and Servalcat increased $\chi_1$ disagreement by 2.93°, 3.19° and 3.15°, whereas CryoNet.Refine produced no detected change (−0.20°; Supplementary Table S33). On matched atoms with fixed occupancy and B factors, all four increased same-map $CC_{mask}$ (Phenix ×1 +0.011, Phenix ×2 +0.019, CryoNet.Refine +0.014, Servalcat +0.053; Holm-adjusted $P \leq 0.0023$; Supplementary Fig. S7). The increases in disagreement persisted after excluding valine or all atom-name-ambiguous residues, for $\chi_2$, after excluding residues near non-polymer components, with poor local backbone agreement or near sequence differences, and at a relaxed packing threshold (12,108 residues; Extended Data Figs. 1 and 2).

The highest-map-fit selection rule takes, within each case, the action with the highest same-map $CC_{mask}$, with ties resolved in favour of no operation. The rule chose Servalcat in 25 cases and Phenix ×2 in one and increased $\chi_1$ disagreement by 2.83° (95% cluster-bootstrap interval 1.85–3.74°) relative to no operation (Extended Data Table 1). In 19 of 26 cases, at least one refined action agreed better with the comparison structure than did the starting model. The rule's observed regret (the selected action's disagreement minus the lowest disagreement among the available actions) was 3.39° (2.56–4.19°). The selected action raised $CC_{mask}$ yet increased disagreement in 85% of cases (cluster-weighted average; 70–100%). The direction held when any one cluster (20/20) or any pair of clusters (190/190) was removed, and raising the required $CC_{mask}$ gain reduced coverage without identifying a threshold that eliminated the observed mean increase in disagreement among accepted refinements (source data available from the corresponding author on request).

At the level of individual side chains, following the rule increased $\chi_1$ disagreement by more than 20° for 763 of the 6,480 packed side chains and decreased it by more than 20° for 358, about two increases for each decrease (cluster-weighted fractions 10.7%, 8.9–12.2%, and 4.9%, 3.9–6.0%; Supplementary Table S30). The cluster-weighted mean local side-chain RMSD from the comparison structure increased by 0.15 Å, and the cluster-weighted fraction above 1 Å rose by 8.9 percentage points. In contrast, only 33 side chains lost agreement with the comparison structure's canonical $\chi_1$ class and 44 gained it. These counts measure reference-class agreement, as distinct from changes in the starting model's class (Supplementary Table S37).

### Supplementary Table S37 — Residue-level and complementary effects at 3.0–3.6 Å

Post hoc summaries of the 3.0–3.6 Å cohort: 167 entries in 131 sequence groups. Counts pool residues; fractions and changes average residues within entry, entries within group and groups equally. Intervals are 95% percentile group-bootstrap intervals (10,000 draws, seed 20261101); no P values were computed.

**a, Residue events under highest-map-fit selection.**

| Event | Count | Residues | Fraction [95% interval] (%) |
|---|---|---|---|
| $\chi_1$ disagreement increased by >20° | 2,806 | 37,504 | 7.20 [6.61, 7.79] |
| $\chi_1$ disagreement decreased by >20° | 1,339 | 37,504 | 3.74 [3.37, 4.12] |
| Reference $\chi_1$-class agreement lost | 189 | 37,504 | 0.43 [0.30, 0.58] |
| Reference $\chi_1$-class agreement gained | 126 | 37,504 | 0.34 [0.23, 0.47] |
| $\chi_1$ class changed from start (non-valine) | 316 | 33,153 | 0.87 [0.64, 1.19] |
| Local side-chain RMSD crossed above 1 Å | 2,495 | 33,387 | 7.30 [6.56, 8.02] |
| Local side-chain RMSD crossed to ≤1 Å | 1,540 | 33,387 | 4.91 [4.41, 5.44] |

The 37,504-residue $\chi_1$ support is common across all five actions. Reference-class agreement uses the original symmetry-aware definition. Direct class changes from the starting model use the nearest of −60°, +60° and 180°; valine is excluded from this count because CG1/CG2 atom-name interchange changes the canonical torsion. Complete rotamer assignments additionally depend on later torsions.

**b, Complementary effects relative to no operation.**

| Strategy | $\Delta\chi_2$ (°) | Local ΔRMSD (Å) | Δ fraction >1 Å (pp) |
|---|---|---|---|
| Phenix ×1 | +1.27 [+0.87, +1.68] | +0.014 [+0.005, +0.023] | +1.09 [+0.50, +1.72] |
| Phenix ×2 | +1.55 [+1.07, +2.04] | +0.018 [+0.008, +0.029] | +1.09 [+0.43, +1.79] |
| Servalcat | +2.15 [+1.77, +2.53] | +0.082 [+0.069, +0.095] | +2.65 [+1.79, +3.51] |
| CryoNet.Refine | −1.74 [−2.01, −1.47] | 0.000 [−0.008, +0.009] | −0.23 [−0.72, +0.32] |
| Highest map fit | +1.97 [+1.60, +2.34] | +0.074 [+0.062, +0.087] | +2.39 [+1.53, +3.27] |
| Calibrated EMRinger | −1.16 [−1.40, −0.92] | −0.003 [−0.008, +0.003] | −0.10 [−0.44, +0.24] |

$\chi_2$ uses 26,487 residues with defined torsions in all five actions. Local side-chain RMSD uses 33,387 residues with valid values in every action, with the original five-residue backbone Kabsch frame and symmetry/flip handling. Each endpoint uses common support across actions. Calibrated-EMRinger rows include retained no-operation choices. pp, percentage points. The coordinate endpoints describe agreement with the comparison structure.

### Supplementary Table S38 — Calibrated acceptance of re-refined models

**a, Fixed rule applied to the 3.0–3.6 Å cohort.** The calibration set contained 182 entries (26 MEDIC, 156 earlier replication) in 149 sequence groups. The selected EMRinger threshold was 0.881229, with calibration coverage 30.3%, accepted-set loss −0.42° and one-sided 90% upper bound −0.12°. The same rule was then applied to the 3.0–3.6 Å cohort (167 entries in 131 sequence groups).

| Strategy | Accepted | Coverage (%) | Overall loss (°) | Accepted-set loss (°) |
|---|---|---|---|---|
| Highest map fit | 161 | 96.2 | +2.41 [+2.12, +2.71] | +2.50 [+2.21, +2.80] |
| Always CryoNet.Refine | 167 | 100.0 | −0.09 [−0.31, +0.13] | −0.09 [−0.31, +0.13] |
| Calibrated EMRinger | 107 | 65.3 | −0.06 [−0.23, +0.11] | −0.09 [−0.34, +0.18] |
| Per-case oracle | 106 | 64.5 | −0.77 [−0.91, −0.63] | −1.20 [−1.36, −1.03] |

Values in brackets are 95% group-bootstrap intervals, added as descriptive summaries for this report. Coverage is group-weighted and therefore differs from the pooled accepted fraction. Overall loss includes zero for retained no-operation choices; accepted-set loss divides the weighted accepted loss by weighted acceptance. The oracle uses the observed comparison outcomes.

The EMRinger rule accepted 106 CryoNet.Refine and one Servalcat candidate, retaining no operation in 60 entries. Its one-sided 90% upper bound on accepted-set loss was +0.084°. The prespecified criteria for the 3.0–3.6 Å cohort were a non-positive accepted-set mean with upper bound ≤+0.5°, coverage ≥50%, and regret below half the highest-map-fit regret with a supporting 90% bootstrap comparison. All three were met: regret was 0.714° versus 3.179° for map-fit selection; the paired difference from half that regret was −0.875° (90% interval −1.042 to −0.701°). Always CryoNet.Refine had regret 0.682°.

**b, Other calibrated rules at target coverage 0.30.**

| Rule | Coverage (%) | Accepted loss (°) | Upper 90% bound | Regret (°) |
|---|---|---|---|---|
| Movement cap | 84.4 | +2.47 | +2.68 | 2.86 |
| EMRinger | 65.3 | −0.09 | +0.08 | 0.71 |
| Q-score | 9.9 | +2.59 | +3.26 | 1.03 |
| Masked FSC | 11.5 | +2.14 | +2.65 | 1.02 |
| Combined veto | 29.4 | +1.94 | +2.28 | 1.34 |
| Logistic (secondary) | 44.3 | −0.57 | −0.36 | 0.52 |

Candidate rules were calibrated over target coverages 0.30, 0.40, 0.50, 0.60, 0.70 and 0.80. Within each target, the attained coverage closest from above was chosen; ties used accepted loss, regret and grid order. Admissibility required a one-sided 90% percentile bootstrap upper bound at or below zero. The rule family was chosen by largest admissible target coverage, then smaller regret and fewer tuned parameters. EMRinger at 0.30 was selected by this procedure; the logistic model was secondary. The upper bounds are bootstrap estimates. All thresholds and program assignments were carried forward from calibration. The full calibration grids and analysis records are available from the corresponding author and will be indexed through the Primum repository.

**Application.** The four documented refinement candidates are generated and scored with `phenix.emringer` against the same full map, together with the deposited model. The refined candidate with the largest score is selected, with exact ties broken by program identifier, and accepted only when its score exceeds the deposited-model score by more than 0.881229; otherwise no operation is retained. Candidates with unavailable scores are ignored; if the deposited score or every candidate score is unavailable, no operation is retained. Reference coordinates are used to calibrate and evaluate this rule, while application uses only candidate coordinates and the map.

### Supplementary Table S39 — Accepting one refiner when its map correlation increases

Each rule offers one refinement configuration and no operation on the same 167 entries in 131 sequence groups from the 3.0–3.6 Å cohort. The refined model is accepted when its support-matched $CC_{mask}$ exceeds the deposited model's value by more than $10^{-12}$; otherwise the deposited model is retained. This post hoc analysis uses the original scores and candidate models.

| Refiner | Accepted | Worse | Overall loss (°) | Accepted-set loss (°) |
|---|---|---|---|---|
| Phenix ×1 | 124 | 101 | +1.28 [+0.92, +1.66] | +1.73 [+1.27, +2.21] |
| Phenix ×2 | 127 | 103 | +1.54 [+1.14, +1.95] | +2.02 [+1.53, +2.53] |
| Servalcat | 157 | 147 | +2.42 [+2.11, +2.75] | +2.57 [+2.25, +2.91] |
| CryoNet.Refine | 128 | 68 | +0.04 [−0.14, +0.22] | +0.05 [−0.19, +0.30] |

Accepted is the number of accepted entries out of 167. Worse counts accepted entries with greater mean packed-side-chain $\chi_1$ disagreement than no operation. Group-weighted coverage was 74.0%, 76.3%, 94.3% and 75.6%, respectively. Overall loss includes zero for retained no-operation choices. Accepted-set loss divides group-weighted accepted loss by group-weighted coverage. Brackets give unadjusted 95% percentile intervals from 10,000 sequence-group bootstrap draws, seed 20261101. Cases are averaged within groups before groups receive equal weight.

This tests the use of a positive map-fit change as an acceptance criterion within one program. The three Phenix/Servalcat rules have positive mean losses and intervals above zero; the CryoNet.Refine interval spans zero.

### Supplementary Table S40 — Release-date sensitivities and CryoNet.Refine training provenance

CryoNet.Refine was initialized from Boltz-2. Boltz-2's experimental PDB training structures were released before 1 June 2023 (Passaro et al., Boltz-2, bioRxiv preprint (2025), section 2). CryoNet.Refine's public training script specifies AFDB-aligned structures, EMDB map crops and a Boltz-2 checkpoint, but gives no release-date cutoff for refinement training (accessed 29 October 2026). The date analysis addresses pretraining chronology; establishing overlap with refinement training requires its dated training-entry list.

The 3.0–3.6 Å cohort sensitivity analysis retained entries by the *starting entry's* initial PDB release date, strictly after each listed date. Their comparison structures were also released later than each date. Three release-date subsets were evaluated, with the calibrated threshold unchanged at 0.881229. The 2022 and January 2023 cuts are date sensitivities; June 2023 is the documented Boltz-2 cutoff.

| Strategy | Accepted | Coverage (%) | Accepted-set loss (°) | Upper 90% bound |
|---|---|---|---|---|
| **Released after 1 January 2022: 90 entries, 76 groups** | | | | |
| Highest map fit | 90 | 100.0 | +2.57 [+2.15, +3.00] | +2.84 |
| Always CryoNet.Refine | 90 | 100.0 | −0.03 [−0.30, +0.23] | +0.14 |
| Calibrated EMRinger | 66 | 74.3 | +0.07 [−0.23, +0.36] | +0.26 |
| **Released after 1 January 2023: 40 entries, 36 groups** | | | | |
| Highest map fit | 40 | 100.0 | +2.14 [+1.58, +2.73] | +2.52 |
| Always CryoNet.Refine | 40 | 100.0 | −0.07 [−0.48, +0.34] | +0.20 |
| Calibrated EMRinger | 32 | 80.6 | +0.08 [−0.38, +0.52] | +0.36 |
| **Released after 1 June 2023: 21 entries, 20 groups** | | | | |
| Highest map fit | 21 | 100.0 | +1.91 [+1.06, +2.85] | +2.49 |
| Always CryoNet.Refine | 21 | 100.0 | −0.20 [−0.76, +0.35] | +0.17 |
| Calibrated EMRinger | 17 | 80.0 | −0.18 [−0.84, +0.46] | +0.25 |

Coverage is group-weighted. Group membership and equal-group weights are recomputed within each date subset. Brackets are descriptive 95% percentile intervals added for this report; the one-sided 90% bounds reproduce the stored analysis (10,000 group-bootstrap draws, seed 20261101). The EMRinger rule met the three stated test criteria in the June 2023 subset; in the two earlier date subsets its accepted-set mean was positive, so the non-positive-mean criterion was not met.

### Supplementary Table S41 — Prespecified within-study paired-map endpoints

The within-study protocol and 90-study roster were fixed before downloading any within-study data. Screening retained 53 pairs from 53 studies, including 52 separate-acquisition pairs and one shared-dataset pair. Four refinements and no operation were scored against the paired higher-resolution model. Fifty of 53 full-map selection losses were positive, two negative and one zero. The half-map analysis contains 44 pairs, with refinement against half 1 and selection by half 2; 31/44 losses were positive. Each study contributes one equally weighted pair.

**a, Decision endpoints.**

| Endpoint (°) | Full map (53 pairs) | Evaluation half (44 pairs) |
|---|---|---|
| Selection loss | +3.43 [+2.85, +4.02] | +2.35 [+1.60, +3.21] |
| Observed regret | +3.87 [+3.38, +4.37] | +2.61 [+1.87, +3.45] |
| Always CryoNet.Refine | +0.57 [+0.23, +0.93] | +0.69 [+0.32, +1.08] |
| Per-case oracle | −0.44 [−0.65, −0.26] | −0.26 [−0.41, −0.13] |

Full-map selection chose Servalcat in 49 pairs, CryoNet.Refine in three and no operation in one. Evaluation-half selection chose Servalcat in 22, CryoNet.Refine in five, Phenix ×1 in one, Phenix ×2 in three and no operation in 13. The oracle selects the smallest observed disagreement among the same five actions.

**b, Individual refinement effects.**

| Refiner | Full-map $\Delta\chi_1$ (°) | Half-map $\Delta\chi_1$ (°) |
|---|---|---|
| Phenix ×1 | +2.11 [+1.38, +2.91] | +3.72 [+2.89, +4.64] |
| Phenix ×2 | +2.70 [+1.87, +3.61] | +4.79 [+3.80, +5.86] |
| Servalcat | +3.57 [+3.00, +4.14] | +3.01 [+2.39, +3.65] |
| CryoNet.Refine | +0.57 [+0.23, +0.93] | +0.69 [+0.32, +1.08] |

Values are means with unadjusted 95% percentile intervals from 10,000 pair-bootstrap draws (PCG64 seed 20261101). The prespecified primary interpretation was A when the selection-loss interval lay above zero and the mean was at least 1.5°; B when the interval lay above zero and the mean was below 1.5°; C when the lower interval bound was at or below zero. The full-map result met A. Half-map and subgroup analyses were secondary. No P values were computed.

### Supplementary Table S42 — Within-study sensitivities and pilot motivation

**a, Prespecified within-study sensitivities.** The primary unit is the study pair. Sequence-group sensitivity first averages within 95% sequence-identity groups; the 53 pairs comprise 50 such groups. Reconstruction and resolution subsets were specified before refinement. Single-pair results below are descriptive values without inferential intervals.

| Subset | Pairs | Full loss (°) | Half pairs | Half loss (°) |
|---|---|---|---|---|
| Separate acquisitions | 52 | +3.50 [+2.93, +4.10] | 43 | +2.40 [+1.65, +3.27] |
| Shared dataset | 1 | −0.07 | 1 | +0.00 |
| Resolution <4.5 Å | 52 | +3.50 [+2.90, +4.08] | 44 | +2.35 [+1.60, +3.21] |
| Resolution ≥4.5 Å | 1 | +0.00 | 0 | — |
| Whole-chain C$\alpha$ RMSD ≤3 Å | 49 | +3.52 [+2.90, +4.13] | 40 | +2.49 [+1.67, +3.43] |

Sequence-group weighting gave full-map selection loss +3.43 [+2.86, +3.98]°, with 48/50 groups positive. Selection using work-half correlation on the same 44 half-map pairs was exploratory and gave +3.52 [+2.72, +4.42]°.

**b, Descriptive baseline and influence checks.** Within-study baseline disagreement ranged from 13.10° to 54.49°, with median 24.99°; the 3.0–3.6 Å cohort and earlier replication medians were 26.5° and 35.9°. Scored-residue C$\alpha$ RMSD ranged from 0.19 to 2.69 Å. All 53 baseline residue counts and means were independently reproduced from the input coordinates. The median selection loss was 3.44°; omitting the largest loss (6CV3, +7.89°) gave +3.35°. These influence summaries are post hoc.

| Descriptive subgroup | Pairs | Selection loss (°) | Always CryoNet (°) |
|---|---|---|---|
| Baseline below 3.0–3.6 Å cohort median | 30 | +3.95 [+3.12, +4.74] | +0.95 [+0.47, +1.44] |
| Baseline at or above 3.0–3.6 Å cohort median | 23 | +2.76 [+1.98, +3.50] | +0.07 [−0.33, +0.47] |

The within-study baseline threshold was fixed at 12.95° before refinement. It was motivated by the pilot split at the 3.0–3.6 Å cohort's exact fifth percentile, 12.946717°. This defines the evaluated starting-agreement range.

**c, pilot endpoints and post hoc stratification.** The pilot cohort contained 22 pairs, 21 sharing a dataset and one using separate acquisitions. The cohort and primary analysis were prespecified; its baseline and provenance stratification was post hoc. These observations motivated the within-study screening criteria. The within-study cohort excludes all studies and entries on the pilot roster. The prespecified five-action analysis includes 22 pairs from 22 studies and 22 sequence groups. Each low-resolution deposited model was refined against its own map and compared with the paired higher-resolution model. All 22 full-map selection losses were positive. The half-map analysis includes 15 pairs and refines against half 1, selecting by half 2. Each pair contributes one independent group.

**Pilot decision endpoints.**

| Endpoint (°) | Full map (22 pairs) | Evaluation half (15 pairs) |
|---|---|---|
| Selection loss | +8.79 [+5.81, +12.50] | +5.02 [+2.48, +7.63] |
| Regret | +8.97 [+6.08, +12.64] | +5.28 [+2.78, +7.79] |
| Always CryoNet.Refine | +3.43 [+1.94, +4.95] | +3.27 [+1.42, +5.16] |
| Per-case oracle | −0.18 [−0.52, 0.00] | −0.26 [−0.74, 0.00] |

Full-map selections were Servalcat 18, Phenix ×1 one, Phenix ×2 two and CryoNet.Refine one. Half-map selections were Servalcat nine, CryoNet.Refine two and no operation four; 11/15 half-map groups had positive loss. The oracle is the per-case minimum among the five offered actions.

**Pilot refinement effects.**

| Refiner | Full-map $\Delta\chi_1$ (°) | Half-map $\Delta\chi_1$ (°) |
|---|---|---|
| Phenix ×1 | +6.78 [+3.95, +10.37] | +8.92 [+5.27, +13.17] |
| Phenix ×2 | +8.09 [+4.83, +12.09] | +10.36 [+6.35, +14.80] |
| Servalcat | +7.67 [+5.81, +9.47] | +7.42 [+5.06, +9.66] |
| CryoNet.Refine | +3.43 [+1.94, +4.95] | +3.27 [+1.42, +5.16] |

Values are equal-group means with unadjusted 95% percentile intervals (10,000 draws, PCG64 seed 20261101). The prespecified primary decision was A when the selection-loss interval lay above zero and the mean was at least 1.5°; B when the interval lay above zero and the mean was below 1.5°; C when the lower interval bound was at or below zero. The full-map result met A. The half-map result is secondary. Twenty-one pairs share a reconstruction dataset; the single separate-acquisition pair (9C0Q/9C0R) had full-map loss +1.79° and half-map loss 0.00°. Its one-case estimates are descriptive.

**Supplementary Table S42 (continued): d, Pilot baseline-disagreement strata (post hoc).** The baseline is no-operation $\chi_1$ disagreement on the fixed common residue set. The threshold is the fifth percentile across the 167 case baselines in the 3.0–3.6 Å cohort, 12.95° (exact value above). Strata were defined after the paired-map results were available. All values below re-aggregate the recorded per-case outputs; they are descriptive sensitivities.

| Stratum | Pairs | Full-map loss (°) | Evaluation-half loss (°; n) |
|---|---|---|---|
| All pairs | 22 | +8.79 [+5.81, +12.50] | +5.02 [+2.48, +7.63]; 15 |
| Baseline <12.95° | 13 | +12.14 [+8.24, +17.48] | +7.26 [+3.59, +10.58]; 9 |
| Baseline ≥12.95° | 9 | +3.95 [+2.31, +5.90] | +1.66 [+0.44, +3.47]; 6 |
| Baseline below the 3.0–3.6 Å cohort minimum (5.39°) | 7 | +16.09 [+11.81, +23.92] | +6.99 [+1.87, +12.11]; 5 |
| Baseline at or above the 3.0–3.6 Å cohort minimum | 15 | +5.38 [+3.53, +7.32] | +4.04 [+1.50, +6.85]; 10 |
| Low entry names high as initial model | 2 | +12.11 [+10.87, +13.35] | +11.26 [+11.15, +11.37]; 2 |
| Other pairs | 20 | +8.46 [+5.33, +12.53] | +4.06 [+1.61, +6.80]; 13 |

| Stratum | Baseline median (°) | Oracle (°) | Always CryoNet (°) |
|---|---|---|---|
| All pairs | 7.4 | −0.18 [−0.52, 0.00] | +3.43 [+1.94, +4.95] |
| Baseline <12.95° | 2.8 | 0.00 [−0.01, 0.00] | +5.11 [+3.30, +6.92] |
| Baseline ≥12.95° | 23.7 | −0.43 [−1.22, 0.00] | +1.01 [−0.50, +2.49] |

All nine higher-baseline pairs had positive full-map loss; five of their six half-map cases had positive loss. Their baseline range was 14.77–35.90°. The 13 lower-baseline pairs include five with scored-C$\alpha$ RMSD below 0.1 Å; the +12.14° estimate refers to the complete 13-pair stratum. The no-operation baseline and the change score share the baseline term, and very small initial disagreement leaves little room for improvement. Baseline strata therefore characterize starting agreement; model provenance is reported separately in Supplementary Table S43. The 3.0–3.6 Å cohort and the earlier replication had median baselines 26.5° and 35.9°, compared with 7.4° in the within-study pairs. Intervals use the same group bootstrap and seed as Supplementary Table S41.

### Supplementary Table S43 — Within-study rosters and model provenance

**a, The 53 within-study pairs.** Baseline and C$\alpha$ RMSD use the fixed residue set. Acquisition relationships are archive-metadata classifications: S denotes separate acquisitions; D denotes a shared dataset. Neither entry in a retained pair names its partner as a starting model. Listed external starting models may be shared. Case-level scores and exclusions are in Supplementary Data 3; study identifiers, map identifiers and reconstruction metadata are in Supplementary Data 4.

| Low | High | Low (Å) | High (Å) | Baseline (°) | Residues | C$\alpha$ RMSD (Å) | Data | Selected | Full loss (°) | Half loss (°) |
|---|---|---|---|---|---|---|---|---|---|---|
| 9E06 | 9E02 | 3.10 | 2.50 | 13.10 | 56 | 0.30 | S | Servalcat | +3.26 | +1.05 |
| 11BR | 10UP | 3.37 | 2.86 | 13.22 | 147 | 0.55 | S | Servalcat | +5.65 | +1.34 |
| 9GW1 | 9GVC | 3.10 | 1.82 | 13.29 | 33 | 0.19 | S | Servalcat | +1.29 | +0.00 |
| 6C06 | 6BZO | 5.15 | 3.38 | 13.56 | 522 | 1.26 | S | No operation | +0.00 | — |
| 6CV3 | 6CV4 | 3.56 | 3.03 | 13.95 | 91 | 0.33 | S | Servalcat | +7.89 | +0.00 |
| 9BML | 9BM0 | 3.30 | 2.80 | 14.04 | 1508 | 0.34 | S | Servalcat | +4.99 | +1.90 |
| 9XHY | 9XI1 | 3.84 | 2.58 | 14.89 | 271 | 0.41 | S | Servalcat | +6.53 | +6.34 |
| 8E79 | 8E8M | 3.71 | 3.13 | 15.05 | 488 | 0.62 | S | Servalcat | +7.85 | +7.09 |
| 8SQH | 8SRS | 3.69 | 2.79 | 15.48 | 503 | 0.34 | S | Servalcat | +5.20 | +5.50 |
| 7CXN | 7CXM | 3.84 | 2.90 | 16.08 | 405 | 0.33 | S | Servalcat | +2.11 | — |
| 9QQK | 9H8V | 3.14 | 2.41 | 16.38 | 491 | 0.84 | S | Servalcat | +3.96 | +0.00 |
| 8VB8 | 8VBH | 3.00 | 2.20 | 17.00 | 218 | 0.38 | S | CryoNet | +0.36 | +0.00 |
| 9I9J | 9I9K | 3.10 | 2.40 | 17.80 | 123 | 0.35 | S | Servalcat | +3.30 | +1.67 |
| 8IU2 | 8Y01 | 3.35 | 2.48 | 17.90 | 166 | 0.38 | S | Servalcat | +4.83 | +1.07 |
| 9VUK | 9VUJ | 3.17 | 2.60 | 18.83 | 694 | 0.64 | S | Servalcat | +3.20 | +3.60 |
| 7XUM | 7XUK | 3.80 | 3.30 | 19.31 | 337 | 0.58 | S | Servalcat | +6.79 | +6.53 |
| 9CDI | 9CDJ | 3.24 | 2.49 | 19.34 | 208 | 1.40 | S | Servalcat | +0.15 | +1.19 |
| 9COH | 9COE | 3.10 | 2.45 | 19.40 | 136 | 0.37 | S | Servalcat | +1.84 | +2.43 |
| 8XM6 | 8XMA | 3.48 | 2.65 | 21.09 | 106 | 0.35 | S | Servalcat | +3.55 | +0.22 |
| 7W5Y | 7W5X | 4.20 | 3.40 | 21.34 | 523 | 0.58 | S | Servalcat | +0.45 | — |
| 7TYW | 7TYF | 3.00 | 2.20 | 21.81 | 112 | 2.09 | S | Servalcat | +1.61 | +0.80 |
| 8WZH | 8WZM | 3.70 | 2.70 | 21.99 | 181 | 0.45 | S | Servalcat | +2.44 | +2.09 |
| 8JKZ | 8JL0 | 3.60 | 3.10 | 22.79 | 207 | 1.27 | S | Servalcat | +5.73 | +0.00 |
| 5ONV | 6T1Y | 4.10 | 3.30 | 22.87 | 172 | 0.51 | S | Servalcat | +6.62 | +4.85 |
| 7T22 | 9ECO | 4.20 | 2.83 | 22.92 | 198 | 1.44 | S | Servalcat | +5.52 | — |
| 7OSJ | 7ZNQ | 3.80 | 3.04 | 24.72 | 235 | 0.50 | S | Servalcat | +3.27 | — |
| 7TKU | 7TI8 | 4.00 | 3.50 | 24.99 | 155 | 0.64 | S | Servalcat | +5.03 | — |
| 7ADM | 7ADJ | 3.50 | 2.80 | 25.16 | 295 | 0.91 | S | Servalcat | +5.88 | +13.12 |
| 9CNY | 9CNW | 4.00 | 2.60 | 25.94 | 288 | 0.96 | S | Servalcat | +5.03 | +5.50 |
| 8V0Q | 8V0N | 3.90 | 3.40 | 26.22 | 375 | 1.32 | S | Servalcat | +4.22 | +0.00 |
| 8Q7H | 8RHZ | 4.10 | 3.37 | 26.53 | 137 | 2.69 | D | CryoNet | −0.07 | +0.00 |
| 7AHQ | 7AHN | 3.60 | 2.90 | 26.82 | 155 | 0.42 | S | Servalcat | +3.84 | +3.67 |
| 9U6S | 9U6I | 4.20 | 3.00 | 27.52 | 105 | 1.11 | S | Servalcat | +1.09 | +0.00 |
| 9G0S | 9G0P | 3.60 | 3.00 | 29.63 | 224 | 0.56 | S | Servalcat | +5.54 | +4.20 |
| 9DI3 | 9DKD | 4.10 | 3.30 | 29.75 | 915 | 0.99 | S | Servalcat | +4.41 | +0.00 |
| 7W0O | 7W0P | 3.78 | 3.16 | 30.35 | 132 | 0.83 | S | Servalcat | +6.19 | +5.77 |
| 6WI9 | 6WZG | 4.30 | 2.30 | 31.01 | 76 | 1.53 | S | Servalcat | +4.56 | +0.00 |
| 7TKT | 7TJ5 | 3.60 | 2.41 | 31.04 | 87 | 0.56 | S | Servalcat | +4.11 | +4.97 |
| 9M84 | 9ISN | 3.61 | 2.97 | 32.01 | 491 | 1.29 | S | Servalcat | +3.31 | +0.00 |
| 8Z6H | 8ZKH | 3.10 | 2.30 | 33.04 | 165 | 0.88 | S | Servalcat | +1.77 | +0.85 |

| Low | High | Low (Å) | High (Å) | Baseline (°) | Residues | Cα RMSD (Å) | Data | Selected | Full loss (°) | Half loss (°) |
|---|---|---|---|---|---|---|---|---|---|---|
| 3J5R | 3J9J | 4.20 | 3.27 | 33.42 | 81 | 0.74 | S | CryoNet | −2.26 | — |
| 9FSS | 9FSP | 4.14 | 3.39 | 34.54 | 514 | 1.14 | S | Servalcat | +3.61 | +0.00 |
| 9PT5 | 9IFO | 3.74 | 2.10 | 35.37 | 592 | 2.27 | S | Servalcat | +2.90 | +0.45 |
| 7EU1 | 7EU0 | 3.86 | 3.16 | 35.44 | 404 | 0.59 | S | Servalcat | +4.62 | +3.02 |
| 8GH9 | 8GHF | 3.80 | 2.70 | 37.51 | 344 | 1.80 | S | Servalcat | +1.05 | +0.00 |
| 8I41 | 8I42 | 3.42 | 2.92 | 37.57 | 95 | 1.04 | S | Servalcat | +2.27 | +0.79 |
| 9YMA | 9YMC | 3.87 | 3.28 | 37.66 | 292 | 2.50 | S | Servalcat | +3.44 | +2.82 |
| 7ZRM | 7ZRK | 3.70 | 3.10 | 38.03 | 308 | 0.76 | S | Servalcat | +3.68 | +2.64 |
| 8A5T | 8PE4 | 3.78 | 3.21 | 38.94 | 307 | 0.96 | S | Servalcat | +1.14 | +0.83 |
| 7YP9 | 7YPA | 3.58 | 3.05 | 40.55 | 563 | 2.17 | S | Servalcat | +2.30 | +1.75 |
| 7PKO | 7PKP | 3.90 | 3.10 | 44.80 | 117 | 0.60 | S | Servalcat | +1.37 | — |
| 8BX5 | 8BXF | 4.20 | 3.40 | 45.15 | 100 | 0.72 | S | Servalcat | +2.93 | +5.28 |
| 7K8A | 7K8B | 3.65 | 2.94 | 54.49 | 299 | 1.04 | S | Servalcat | +1.66 | — |

**b, Recorded external starting models and deposition order.** PDB starting-model fields and the reconstruction relationships were checked before refinement; the study-metadata roster is Supplementary Data 4. Blank fields are shown as a dash. Deposition order is provided for provenance; the within-study design did not require the higher-resolution entry to have a later release date.

| Low | High | Low deposited | High deposited | Low entry: initial models | High entry: initial models |
|---|---|---|---|---|---|
| 11BR | 10UP | 2026-02-16 | 2026-02-09 | — | — |
| 3J5R | 3J9J | 2013-10-28 | 2015-02-02 | — | — |
| 5ONV | 6T1Y | 2017-08-04 | 2019-10-07 | — | 5OOD |
| 6C06 | 6BZO | 2017-12-27 | 2017-12-25 | — | — |
| 6CV3 | 6CV4 | 2018-03-27 | 2018-03-27 | — | — |
| 6WI9 | 6WZG | 2020-04-09 | 2020-05-13 | — | — |
| 7ADM | 7ADJ | 2020-09-15 | 2020-09-15 | — | — |
| 7AHQ | 7AHN | 2020-09-25 | 2020-09-24 | 5OOD | 6FHL |
| 7CXN | 7CXM | 2020-09-02 | 2020-09-02 | — | — |
| 7EU1 | 7EU0 | 2021-05-15 | 2021-05-15 | — | — |
| 7K8A | 7K8B | 2020-09-26 | 2020-09-26 | — | — |
| 7OSJ | 7ZNQ | 2021-06-08 | 2022-04-21 | — | — |
| 7PKO | 7PKP | 2021-08-26 | 2021-08-26 | — | — |
| 7T22 | 9ECO | 2021-12-02 | 2024-11-14 | — | — |
| 7TKT | 7TJ5 | 2022-01-17 | 2022-01-14 | 6CJU | 6CJQ |
| 7TKU | 7TI8 | 2022-01-17 | 2022-01-13 | 1SXJ | 1SXJ |
| 7TYW | 7TYF | 2022-02-14 | 2022-02-13 | — | — |
| 7W0O | 7W0P | 2021-11-18 | 2021-11-18 | — | — |
| 7W5Y | 7W5X | 2021-11-30 | 2021-11-30 | — | — |
| 7XUM | 7XUK | 2022-05-19 | 2022-05-18 | — | — |
| 7YP9 | 7YPA | 2022-08-03 | 2022-08-03 | 6ALF | 6ASX |
| 7ZRM | 7ZRK | 2022-05-04 | 2022-05-04 | 7NNL | 7NNL |
| 8A5T | 8PE4 | 2022-06-16 | 2023-06-13 | 1M1C | 1M1C |
| 8BX5 | 8BXF | 2022-12-07 | 2022-12-08 | — | — |
| 8E79 | 8E8M | 2022-08-23 | 2022-08-25 | — | — |
| 8GH9 | 8GHF | 2023-03-09 | 2023-03-10 | — | — |

| Low | High | Low deposited | High deposited | Low entry: initial models | High entry: initial models |
|---|---|---|---|---|---|
| 8I41 | 8I42 | 2023-01-18 | 2023-01-18 | — | — |
| 8IU2 | 8Y01 | 2023-03-23 | 2024-01-21 | — | — |
| 8JKZ | 8JL0 | 2023-06-02 | 2023-06-02 | — | — |
| 8Q7H | 8RHZ | 2023-08-16 | 2023-12-17 | — | — |
| 8SQH | 8SRS | 2023-05-04 | 2023-05-06 | — | — |
| 8V0Q | 8V0N | 2023-11-17 | 2023-11-17 | — | — |
| 8VB8 | 8VBH | 2023-12-12 | 2023-12-12 | — | — |
| 8WZH | 8WZM | 2023-11-01 | 2023-11-02 | 1DGF | 1DGF |
| 8XM6 | 8XMA | 2023-12-27 | 2023-12-27 | — | — |
| 8Z6H | 8ZKH | 2024-04-19 | 2024-05-16 | — | — |
| 9BML | 9BM0 | 2024-05-02 | 2024-05-02 | — | — |
| 9CDI | 9CDJ | 2024-06-25 | 2024-06-25 | 5OF9 | 5OF9 |
| 9CNY | 9CNW | 2024-07-15 | 2024-07-15 | — | — |
| 9COH | 9COE | 2024-07-16 | 2024-07-16 | 5YXW, 8UUP | 8UUP |
| 9DI3 | 9DKD | 2024-09-05 | 2024-09-08 | — | — |
| 9E06 | 9E02 | 2024-10-17 | 2024-10-17 | — | — |
| 9FSS | 9FSP | 2024-06-21 | 2024-06-21 | — | — |
| 9G0S | 9G0P | 2024-07-08 | 2024-07-08 | — | — |
| 9GW1 | 9GVC | 2024-09-26 | 2024-09-23 | 8B11 | — |
| 9I9J | 9I9K | 2025-02-06 | 2025-02-06 | 1OKV, 8P6Y | 1OKV, 8P6Y |
| 9M84 | 9ISN | 2025-03-11 | 2024-07-18 | — | — |
| 9PT5 | 9IFO | 2025-07-27 | 2025-02-18 | — | — |
| 9QQK | 9H8V | 2025-04-01 | 2024-10-29 | 9H2R | — |
| 9U6S | 9U6I | 2025-03-23 | 2025-03-23 | 6QUM | 6QUM |
| 9VUK | 9VUJ | 2025-07-13 | 2025-07-13 | — | — |
| 9XHY | 9XI1 | 2025-11-02 | 2025-11-02 | — | — |
| 9YMA | 9YMC | 2025-10-09 | 2025-10-09 | 4W5N | 4W5N |

**Supplementary Table S43 (continued): c, Within-study screening and exclusions.** The applied order was coordinate availability, recorded direct starting-model links, sequence/residue eligibility, scored-residue C$\alpha$ RMSD, baseline $\chi_1$ disagreement, and primary-map validity. Counts after each stage were 90, 82, 80, 78, 67, 53 and 53. Each pair below is assigned its first failing criterion. All exclusions preceded refinement.

| Low | High | First exclusion reason |
|---|---|---|
| 8HHB | 8HH6 | Baseline $\chi_1$ disagreement below 12.95° |
| 8DBC | 8DBL | Baseline $\chi_1$ disagreement below 12.95° |
| 8E2I | 8E2D | Coordinates unavailable in the required PDB format |
| 9DB1 | 9DBZ | Coordinates unavailable in the required PDB format |
| 7NV1 | 7NV0 | Scored-residue C$\alpha$ RMSD outside 0.1–3.0 Å |
| 27ZK | 27ZR | Scored-residue C$\alpha$ RMSD outside 0.1–3.0 Å |
| 8AZ0 | 8AZ4 | One entry records the other as an initial model |
| 8XWQ | 8XWP | Baseline $\chi_1$ disagreement below 12.95° |
| 5U07 | 5U0A | Scored-residue C$\alpha$ RMSD outside 0.1–3.0 Å |
| 9IK9 | 9IK8 | Coordinates unavailable in the required PDB format |
| 9HGA | 9HG6 | Coordinates unavailable in the required PDB format |
| 8WQG | 8WQA | Scored-residue C$\alpha$ RMSD outside 0.1–3.0 Å |
| 6VWH | 6VWG | Identity below 95% or fewer than 20 packed $\chi_1$ residues |
| 9MGG | 9MGF | Scored-residue C$\alpha$ RMSD outside 0.1–3.0 Å |
| 8VAL | 8VAT | Coordinates unavailable in the required PDB format |
| 7X05 | 7X06 | Baseline $\chi_1$ disagreement below 12.95° |
| 9HXE | 9HXG | Baseline $\chi_1$ disagreement below 12.95° |
| 7OXR | 7OXP | Coordinates unavailable in the required PDB format |
| 8G9N | 8UCU | One entry records the other as an initial model |
| 9L8N | 9L8M | Baseline $\chi_1$ disagreement below 12.95° |
| 9BDM | 9ZLD | Identity below 95% or fewer than 20 packed $\chi_1$ residues |
| 8ONA | 8ON9 | Baseline $\chi_1$ disagreement below 12.95° |
| 8PFG | 8PIB | Baseline $\chi_1$ disagreement below 12.95° |
| 7OO0 | 7ONJ | Baseline $\chi_1$ disagreement below 12.95° |
| 7ZJ8 | 7ZJ6 | Baseline $\chi_1$ disagreement below 12.95° |
| 7ORJ | 7ORN | Scored-residue C$\alpha$ RMSD outside 0.1–3.0 Å |
| 8EA3 | 8EA4 | Baseline $\chi_1$ disagreement below 12.95° |
| 7R16 | 7R1B | Scored-residue C$\alpha$ RMSD outside 0.1–3.0 Å |
| 8W9R | 8K4A | Scored-residue C$\alpha$ RMSD outside 0.1–3.0 Å |
| 7BAM | 7BAN | Baseline $\chi_1$ disagreement below 12.95° |
| 9BUC | 9BUD | Coordinates unavailable in the required PDB format |
| 7Z7T | 7P64 | Baseline $\chi_1$ disagreement below 12.95° |
| 7KFU | 7KFT | Scored-residue C$\alpha$ RMSD outside 0.1–3.0 Å |
| 7XNH | 8XGD | Baseline $\chi_1$ disagreement below 12.95° |
| 9M2P | 9M2R | Scored-residue C$\alpha$ RMSD outside 0.1–3.0 Å |
| 9GXP | 9GXK | Scored-residue C$\alpha$ RMSD outside 0.1–3.0 Å |
| 7M3L | 7M3N | Coordinates unavailable in the required PDB format |

Of the 53 retained pairs, eight lacked usable deposited half maps and 7TKU failed the prespecified half-map frame check, leaving 44 half-map pairs. The 7TKU half-map cell differed from the primary map by 87.5%; its full-map analysis was retained. All generated full- and half-map candidates passed technical checks; no configuration was excluded. The paired higher-resolution map was not used for refinement, selection or scoring; the reference coordinates supplied the structural endpoint.

**Supplementary Table S43 (continued): d, pilot roster and provenance. The 22 pilot pairs.** Baseline values use the fixed common residue set. C$\alpha$ RMSD uses the scored-residue superposition. Asterisks mark RMSD below 0.1 Å. Dates are PDB deposition dates; initial-model relations are from PDB starting-model fields.

| Low | High | Baseline (°) | Residues | C$\alpha$ RMSD (Å) | Low deposited | High deposited | Recorded initial-model relation | Data relation | Selected | Loss (°) |
|---|---|---|---|---|---|---|---|---|---|---|
| 7ZKI | 7ZAH | 0.02 | 236 | 0.0005* | 2022-04-13 | 2022-03-22 | Unrecorded | Shared dataset | Servalcat | +12.61 |
| 8WK4 | 8WJR | 0.02 | 30 | 0.0005* | 2023-09-27 | 2023-09-26 | Low lists: 7CBL | Shared dataset | Phenix ×2 | +39.31 |
| 9BXS | 9BXC | 0.02 | 403 | 0.0005* | 2024-05-22 | 2024-05-22 | Low names high | Shared dataset | Phenix ×1 | +13.35 |
| 9DVU | 9DVS | 0.12 | 501 | 0.1674 | 2024-10-08 | 2024-10-08 | Common parent: 8SYI | Shared dataset | Servalcat | +12.71 |
| 9SWO | 9SWN | 1.21 | 335 | 0.0259* | 2025-10-07 | 2025-10-07 | Unrecorded | Shared dataset | Servalcat | +11.26 |
| 9EUJ | 9EUK | 2.69 | 380 | 0.0512* | 2024-03-27 | 2024-03-27 | Unrecorded | Shared dataset | Servalcat | +12.01 |
| 7MWD | 7JQ9 | 2.80 | 1271 | 0.2695 | 2021-05-16 | 2020-08-10 | Unrecorded | Shared dataset | Servalcat | +11.40 |
| 9ZUB | 9ZQY | 7.13 | 178 | 0.1668 | 2025-12-23 | 2025-12-19 | Low names high; common external models | Shared dataset | Servalcat | +10.87 |
| 8GAU | 8GAT | 7.14 | 202 | 0.3454 | 2023-02-23 | 2023-02-23 | Unrecorded | Shared dataset | Servalcat | +8.14 |
| 8CT2 | 7UZE | 7.18 | 114 | 0.1414 | 2022-05-13 | 2022-05-09 | Unrecorded | Shared dataset | Servalcat | +5.92 |
| 7UQI | 7UQK | 7.18 | 206 | 0.1318 | 2022-04-19 | 2022-04-19 | Unrecorded | Shared dataset | Servalcat | +7.09 |
| 8AG5 | 8AG4 | 7.65 | 163 | 0.1253 | 2022-07-19 | 2022-07-19 | Common parent: 1JEY | Shared dataset | Phenix ×2 | +0.05 |
| 7NYU | 7NZ1 | 8.24 | 461 | 0.4171 | 2021-03-23 | 2021-03-23 | Common parent: 4HEA | Shared dataset | Servalcat | +13.09 |
| 9TEP | 9TEQ | 14.77 | 171 | 1.6982 | 2025-11-25 | 2025-11-25 | Unrecorded | Shared dataset | Servalcat | +5.00 |
| 6XLA | 6XL9 | 17.05 | 60 | 0.3870 | 2020-06-28 | 2020-06-28 | Low lists: 6WL5 | Shared dataset | Servalcat | +10.23 |
| 9C0Q | 9C0R | 18.15 | 406 | 0.3165 | 2024-05-27 | 2024-05-27 | Unrecorded | Separate acquisition | Servalcat | +1.79 |
| 7K9X | 7KA4 | 21.25 | 160 | 0.2289 | 2020-09-29 | 2020-09-29 | Unrecorded | Shared dataset | Servalcat | +4.83 |
| 35VF | 35VC | 23.71 | 103 | 0.6575 | 2026-05-19 | 2026-05-19 | Low lists: 6CJQ | Shared dataset | Servalcat | +5.95 |
| 8G9B | 8FUZ | 24.55 | 172 | 1.2512 | 2023-02-21 | 2023-01-18 | Unrecorded | Shared dataset | Servalcat | +2.87 |
| 6POE | 6UUZ | 26.07 | 469 | 2.4335 | 2019-07-03 | 2019-11-01 | Common parent: 3MWD | Shared dataset | Servalcat | +2.83 |
| 8DRQ | 8DS3 | 30.11 | 231 | 1.0897 | 2022-07-21 | 2022-07-21 | Unrecorded | Shared dataset | CryoNet | +0.72 |
| 9MYJ | 9P6H | 35.90 | 173 | 0.5969 | 2025-01-21 | 2025-06-19 | High names low | Shared dataset | Servalcat | +1.30 |

Two low entries name their high-resolution partner as an initial model; one high entry names the low-resolution partner; five pairs record a common external parent, including 9ZUB/9ZQY, which also has a direct link (shared models: 1NL0, 6I58 and AF-F2RM35-F1-V4). Thirteen low entries provide no starting-model identifier. The audit combines coordinate comparisons with PDB metadata; unrecorded links remain unresolved. Of the 22 pairs, 14 were deposited on the same day, six high-resolution entries first and two low-resolution entries first. Reconstruction relationships include shared-particle maps and consensus/focused-map pairs.

**Supplementary Table S43 (continued): e, Eight excluded pilot pairs.**

| Low | High | Exclusion reason |
|---|---|---|
| 9R6O | 9R6P | Scored-C$\alpha$ RMSD above 3 Å |
| 8G8B | 8G86 | Aligned sequence identity below 95% |
| 9CAZ | 9C5Y | Scored-C$\alpha$ RMSD above 3 Å |
| 7QO9 | 7QO7 | Scored-C$\alpha$ RMSD above 3 Å |
| 9C45 | 9C44 | Scored-C$\alpha$ RMSD above 3 Å |
| 7UNT | 7UNU | Reference-coordinate download failed |
| 7SN2 | 7SN3 | Aligned sequence identity below 95% |
| 9CWP | 9CWR | Fewer than 20 packed residues with defined $\chi_1$ |

The metadata roster contained 30 studies; eight download/preparation exclusions left 22 analysed pairs. No additional full-map pair was excluded after refinement. Entry 7MWD remained in the full-map analysis but its half-map branch was excluded before refinement because the half-map cell differed by 3.03% from the primary map. Fifteen pairs had usable half maps in the primary-map frame. Candidate order was determined reproducibly from the low-resolution PDB identifiers using the algorithm recorded with the analysis code.

### Supplementary Table S44 — Fixed acceptance rules on the within-study cohorts

**a, Prespecified within-study evaluation.** The rules calibrated on MEDIC and the earlier replication were applied unchanged to all 53 within-study pairs. Starting entry 7TKU also occurs in the calibration cohort, paired there with 8DQX and here with 7TI8. The designated EMRinger rule used the exact gain threshold 0.881229, accepted 23 CryoNet.Refine outputs and one Servalcat output, and retained no operation for 29 pairs.

| Rule | Accepted | Coverage (%) | Accepted loss (°) | Upper 90% bound | Regret (°) |
|---|---|---|---|---|---|
| EMRinger (designated) | 24 | 45.3 | +1.00 | +1.31 | 0.89 |
| Highest map fit | 52 | 98.1 | +3.50 | +3.88 | 3.87 |
| Always CryoNet.Refine | 53 | 100.0 | +0.57 | +0.80 | 1.01 |
| Always Servalcat | 53 | 100.0 | +3.57 | +3.94 | 4.00 |
| Movement cap | 29 | 54.7 | +3.19 | +3.67 | 2.19 |
| Q-score | 7 | 13.2 | +3.97 | +4.87 | 0.96 |
| Masked FSC | 8 | 15.1 | +3.27 | +4.53 | 0.93 |
| Combined veto | 9 | 17.0 | +1.66 | +2.85 | 0.72 |
| Logistic (secondary) | 25 | 47.2 | +0.57 | +1.02 | 0.71 |

Coverage is the fraction of accepted pairs. The designated rule's overall loss, including no-operation choices, was +0.45°, while its accepted-set loss was +1.00°. The loss criterion required a non-positive accepted-set mean and a one-sided 90% upper bound no greater than +0.5°; the mean and +1.31° bound failed this criterion. Coverage of 45.3% failed the 50% coverage requirement. The regret criterion was met: regret was 0.89° versus half the highest-map-fit regret of 1.94°, with paired 90% difference interval −1.36 to −0.73°. The designated rule therefore failed the joint test. Other calibrated rules are shown at their calibration target coverage of 0.30. No operation has overall loss zero and regret 0.44°; the oracle accepted 24 cases, with overall loss −0.44° and zero regret. Highest-map-fit accepted-set loss is +3.50° over 52 accepted pairs; its overall loss is +3.43° over all 53.

**b, Within-study release-date sensitivities.**

| Date filter | Released after | Pairs | Coverage (%) | Accepted loss (°) | Always CryoNet (°) |
|---|---|---|---|---|---|
| Low entry | 2022-01-01 | 42 | 54.8 | +1.00 | +0.77 |
| Low entry | 2023-01-01 | 34 | 50.0 | +0.96 | +0.74 |
| Low entry | 2023-06-01 | 30 | 50.0 | +0.92 | +0.78 |
| High entry | 2022-01-01 | 42 | 54.8 | +1.00 | +0.77 |
| High entry | 2023-01-01 | 34 | 50.0 | +0.96 | +0.74 |
| High entry | 2023-06-01 | 31 | 48.4 | +0.92 | +0.78 |

The June 2023 date concerns Boltz-2 pretraining, as described in Supplementary Table S40. These filters do not establish CryoNet.Refine's refinement-training membership. The designated rule failed the joint test in every release-date subset. Intervals and bounds use 10,000 pair-bootstrap draws, seed 20261101.

**c, Pilot evaluation.** The thresholds calibrated on MEDIC and the earlier replication were applied unchanged to the 22 pairs. The designated EMRinger rule used a gain threshold of 0.881229 and accepted 11 CryoNet.Refine outputs, retaining no operation for the remaining 11 cases.

| Rule | Accepted | Coverage (%) | Accepted loss (°) | Upper 90% bound | Regret (°) |
|---|---|---|---|---|---|
| EMRinger (designated) | 11 | 50.0 | +4.38 | +5.77 | 2.37 |
| Highest map fit | 22 | 100.0 | +8.79 | +11.08 | 8.97 |
| Always CryoNet.Refine | 22 | 100.0 | +3.43 | +4.42 | 3.61 |
| Always Servalcat | 22 | 100.0 | +7.67 | +8.87 | 7.85 |
| Movement cap | 13 | 59.1 | +6.16 | +7.71 | 3.82 |
| Q-score | 5 | 22.7 | +9.01 | +12.90 | 2.23 |
| Masked FSC | 4 | 18.2 | +8.01 | +11.14 | 1.64 |
| Combined veto | 4 | 18.2 | +2.38 | +4.07 | 0.61 |
| Logistic (secondary) | 10 | 45.5 | +3.25 | +4.98 | 1.66 |

Coverage is equal-group weighted. Accepted-set loss is conditional on acceptance; overall loss for the designated rule was +2.19°. The loss criterion required a non-positive accepted-set mean and a one-sided 90% upper bound no greater than +0.5°; its +4.38° mean and +5.77° bound failed this criterion. The 50% coverage requirement and the regret criterion were met: regret was 2.37° versus half the highest-map-fit regret of 4.48°, with paired 90% difference interval −3.18 to −1.13°. The fixed rule therefore failed the prespecified joint test. All other rules above are shown at their calibration target coverage of 0.30. No operation has overall loss zero and regret 0.18°; the oracle accepted three cases, with overall loss −0.18° and zero regret.

**Release-date sensitivities.**

| Entries released after | Cases | EMRinger coverage (%) | Accepted loss (°) | Always CryoNet (°) |
|---|---|---|---|---|
| 2022-01-01 | 17 | 52.9 | +4.55 | +3.37 |
| 2023-01-01 | 14 | 50.0 | +4.61 | +3.47 |
| 2023-06-01 | 11 | 45.5 | +5.44 | +3.89 |

The low- and high-entry date filters gave identical memberships for these cuts. The June 2023 date concerns Boltz-2 pretraining, as described in Supplementary Table S40. Intervals and one-sided bounds use 10,000 group-bootstrap draws, seed 20261101.
